# Ground nesting of soft eggs by extinct birds and a new parity mode switch hypothesis for the evolution of animal reproduction

**DOI:** 10.1101/2024.08.15.607182

**Authors:** M. Jorge Guimarães, Junyou Wang, Xuemin Zhang, Qiang Sun, M. Fátima Cerqueira, Yi-Hsiu Chung, Richard Deng, Bin Guo, Pedro Alpuim, Feimin Ma, Xiaobing Wang, Tzu-Chen Yen

## Abstract

Nearshore ground nesting of soft eggs by extinct birds is demonstrated here, providing a new explanation for the abundance of bird fossils in early Cretaceous lacustrine environments, where humidity conditions required for soft egg incubation would have been present. This reinforces recent findings of Archaeopteryx soft eggs near Jurassic marine environments, the possibility that wings and elongated feathers developed primarily in association with nest protection on the ground and only secondarily with flight, and the origin of flight from the ground up. Notably, soft eggs preceded rigid eggs in evolution, but both crocodiles, whose ancestors seem to have antedated bird precursors, and extant birds reproduce exclusively via hard-shelled eggs. Therefore, an explanation is in order for how reproduction via soft eggs could have occurred in the bird lineage in-between two evolutionary moments of reproduction via rigid eggs. In alternative to the commonly accepted convergent evolution of viviparity and rigid eggshells, a parity mode switch hypothesis is presented here. It postulates the existence, since the rise of animals, of an inherited ancestral parity mode switch between viviparity and oviparity. This switch would have evolved to embrace hard-shelled oviparity after rigid eggshells appeared in evolution. Commitment to a particular parity mode or eggshell type may have conditioned survival of entire animal groups, especially during major extinction events, explaining, among others, the extinction of all birds that reproduced via soft eggshells.

## INTRODUCTION

All extant birds are oviparous and lay hard-shelled eggs. However, little is known on the reproduction of their extinct ancestors^1^. Certain species of turtles and geckos reproduce via hard-shelled eggs whereas others reproduce via soft eggs^2,3^. These two types of eggs were also shown to have co-existed among pterosaurs^4,5^ and non-avian dinosaurs^6^. Incidentally, a soft egg was discovered in association with marine reptiles^7^, a group of extinct animals that had been previously been associated with viviparity^8^. Following the discovery of soft eggs in *Archaeopteryx*, a bird ancestor^9–12^, we report here, for the first time, the discovery of soft eggs among extinct birds.

The discovery of soft eggs in the bird lineage requires revisiting several published reports, as explained in the discussion. In addition, a new hypothesis is presented here for the evolution of eggshell types in birds, which, if correct, will have profound implications in our understanding of not only bird reproduction evolution but of all animals, and is likely to explain the extinction of major animal groups.

## RESULTS

NUS_A6_2016 is a well preserved and fully articulated specimen of the ornithuromorph *Iteravis huchzermeyeri*, from the Lower Cretaceous Yixian Formation of the Sihedang locality, near Lingyuan, Liaoning, in Northeast China^13^. It fossilized with a globular 3D structure, designated NUS_A6_2016 E1 (Egg E1 from specimen NUS_A6_2016), ensnared in its right wing bones, which structure is approximately 1 cm in size (9.68 x 6.58 x 3.58 mm; Figures 1 and S1-S3).

**Figure 1.**
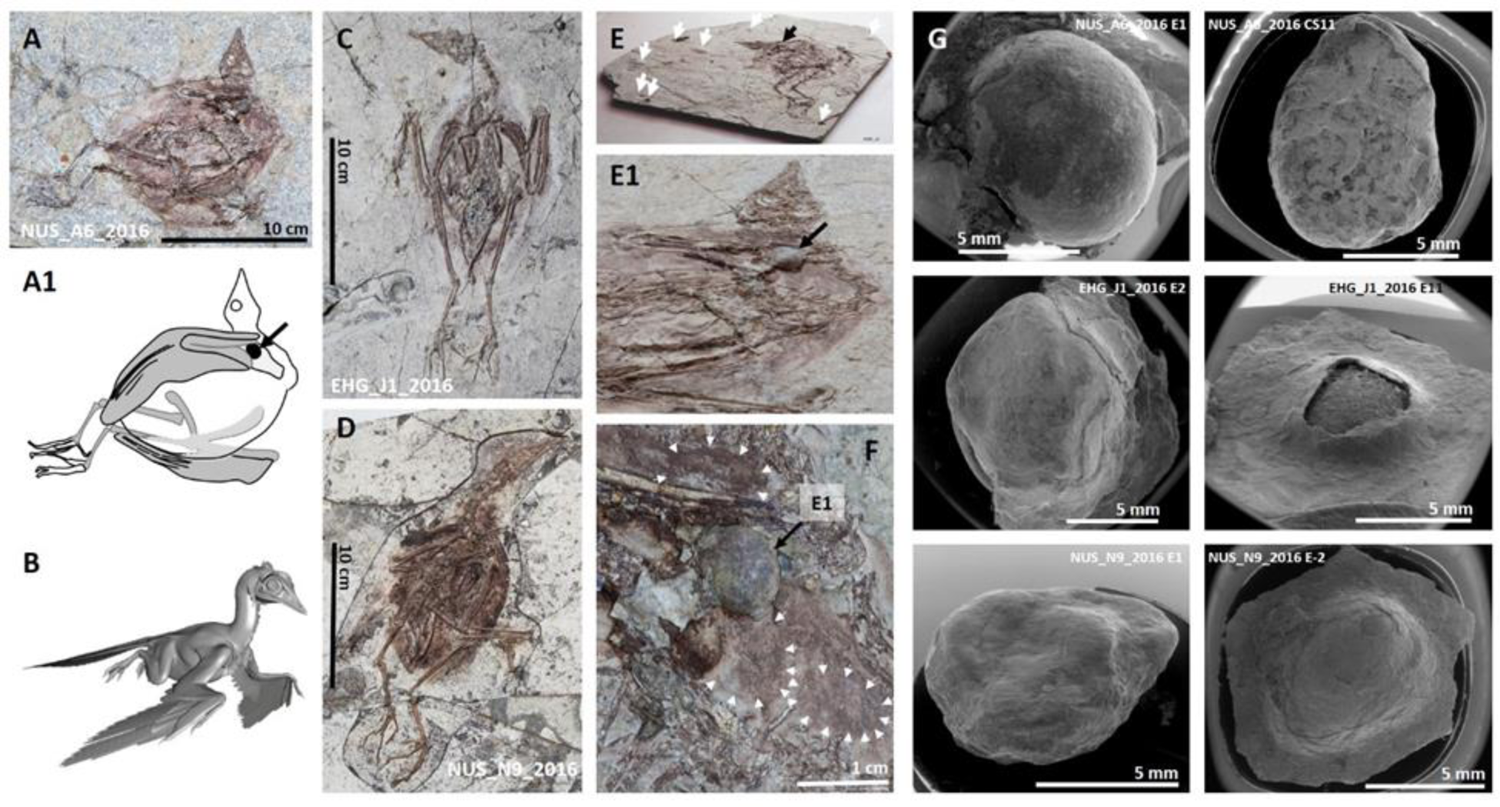
Several *I. huchzermeyeri* specimens fossilized in the same pose and surrounded by similarly sized globular structures of different 3D shape. A: specimen NUS_A6_2016 fossilized in ventro-lateral view (illustrated in **A1** where the NUS_A6_2016 E1 egg is indicated by a black arrow) of the nesting posture modeled in **B**. **C** and **D** are specimens EHG_J1_2016 and NUS_N9_2016 of the same ornithuromorph, which fossilized, respectively, in ventral and dorsolateral view of the same posture. **E**: side view of NUS_A6_2016’s main slab, where the NUS_A6_2016 E1 egg (black arrow) is located. White arrows indicate slab surface convexities and concavities caused by structures similar in size to NUS_A6_2016 E1. The latter is observable in a side view of the right wing area in **E1**. **F:** NUS_A6_2016 E1, surface exposed against the folded right wing of NUS_A6_2016, is surrounded by bi-dimensional impressions of structures with compatible 3D characteristics to its own. **G**: Globular structures extracted under Computerized Tomography (CT) guidance from the slabs of the three fossilized birds shown in A, B and C are shown here under Scanning Electron Microscopy (SEM). NUS_A6_2016 E1 is shown on the top left and slab-embedded NUS_A6_2016 CS11 from the same fossil is shown on the top right. EHG_J1_2016 and EHG_J1_2016 E11 shown in the center were slab-embedded in the EHG_J1_2016 fossil. In turn, NUS_N9_2016 E1 and NUS_N9_2016 E-2 shown at the bottom were slab-embedded in the NUS_N9_2016 fossil (see also Figures S17-S26).

This specimen’s fossilized pose fits into a hypothesized nesting posture characterized by symmetrical flexion and moderate abduction of both wings, symmetrical flexion of the hindlimbs and slight flexion of the vertebral column first identified elsewhere (Figure S4) ^9–12^. Six more fully articulated *I. huchzermeyeri* specimens fossilized in different views of the same posture were also identified and studied here (Figures 1C, 1D, and S5-S9).

Surface convexities and concavities caused by globular radiodense structures of similar size to NUS_A6_2016 E1 (Figures S10-S12), as well as bi-dimensional impressions of similar size to its sagittal, coronal and transversal outlines, were detected in both slabs of the NUS_A6_2016 fossil (Figures 1E-1F and S13).

NUS_A6_2016 E1 was extracted from its fossil slab. It has smooth but unleveled surface with pinpoint depressions and 3D asymmetric shape (Figures 1G and S14 – S15). This and other globular structures derived from all seven *I. huchzermeyeri* specimens reported here (Figure S16) measure approximately 0.6 to 1.0 cm in their longer axis, similar to the smallest hummingbird eggs^14^, but, unlike the latter, which are rigid and have only two 2D outlines (coronal and sagittal outlines are coincident but differ from the transversal view), present various 3D shapes that account for entirely different 2D outlines in coronal, sagittal, and transversal views (Figures 1G and S17-S26).

The already mentioned NUS_A6_2016 E1, EHG_J1_2016 E11 (Figure 1G) extracted from specimen EHG_J1_2016 (Figure 1C), NUS_N9_2016 E-1 (Figure 1G) extracted from specimen NUS_N9_2016 (Figure 1D), NUS_A6_2016 CS1, like NUS_A6_2016 E1 extracted from specimen NUS_A6_2016, and other similar structures from these and other ornithuromorph specimens, when submitted to micro-CT scans at 3–10 μm resolution, show a continuous, radiodense, thin layer that delimits their periphery, approximately 0.1 mm or less thick (Figures 2A-2D, S27, and S28).

**Figure 2.**
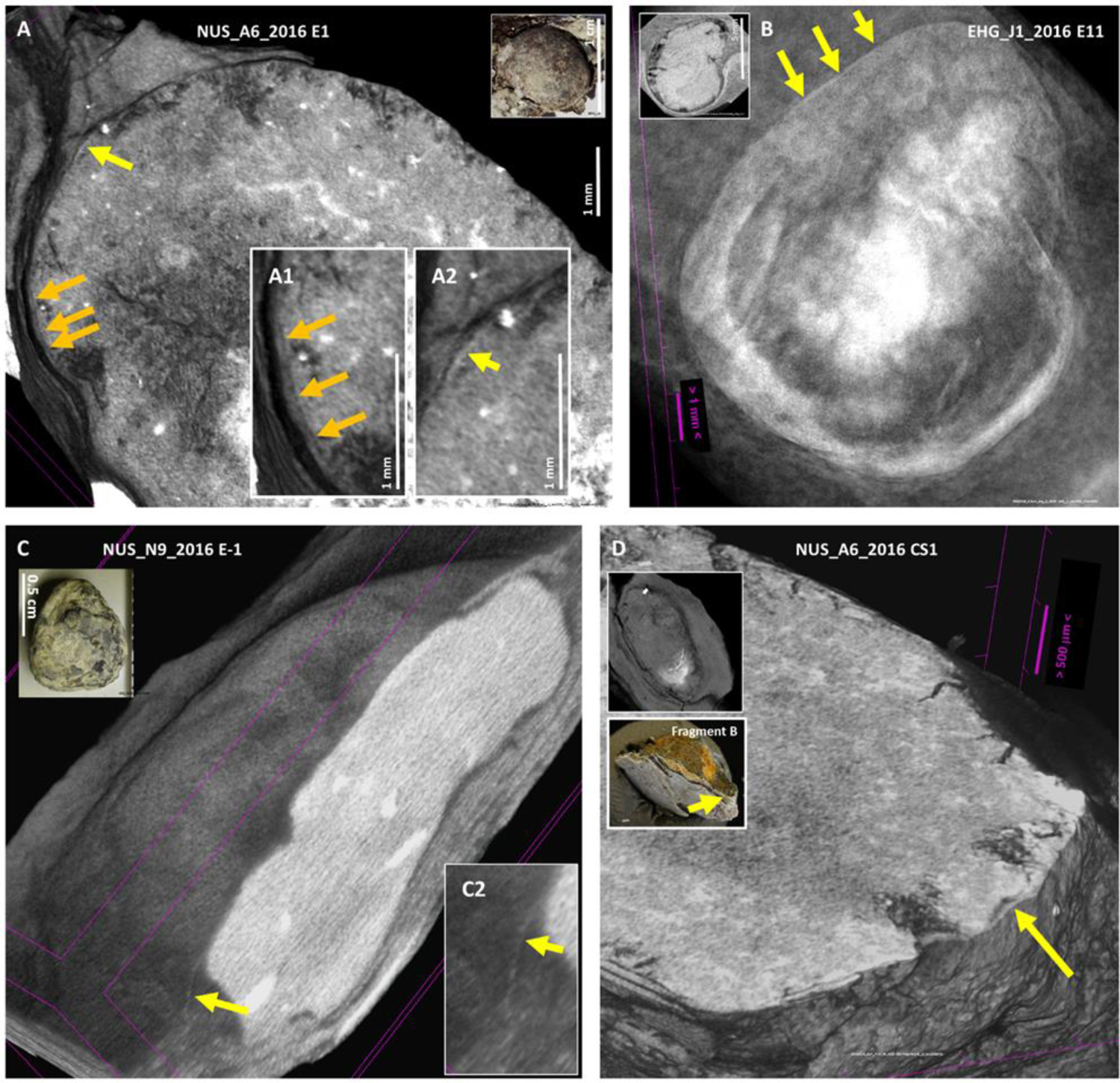
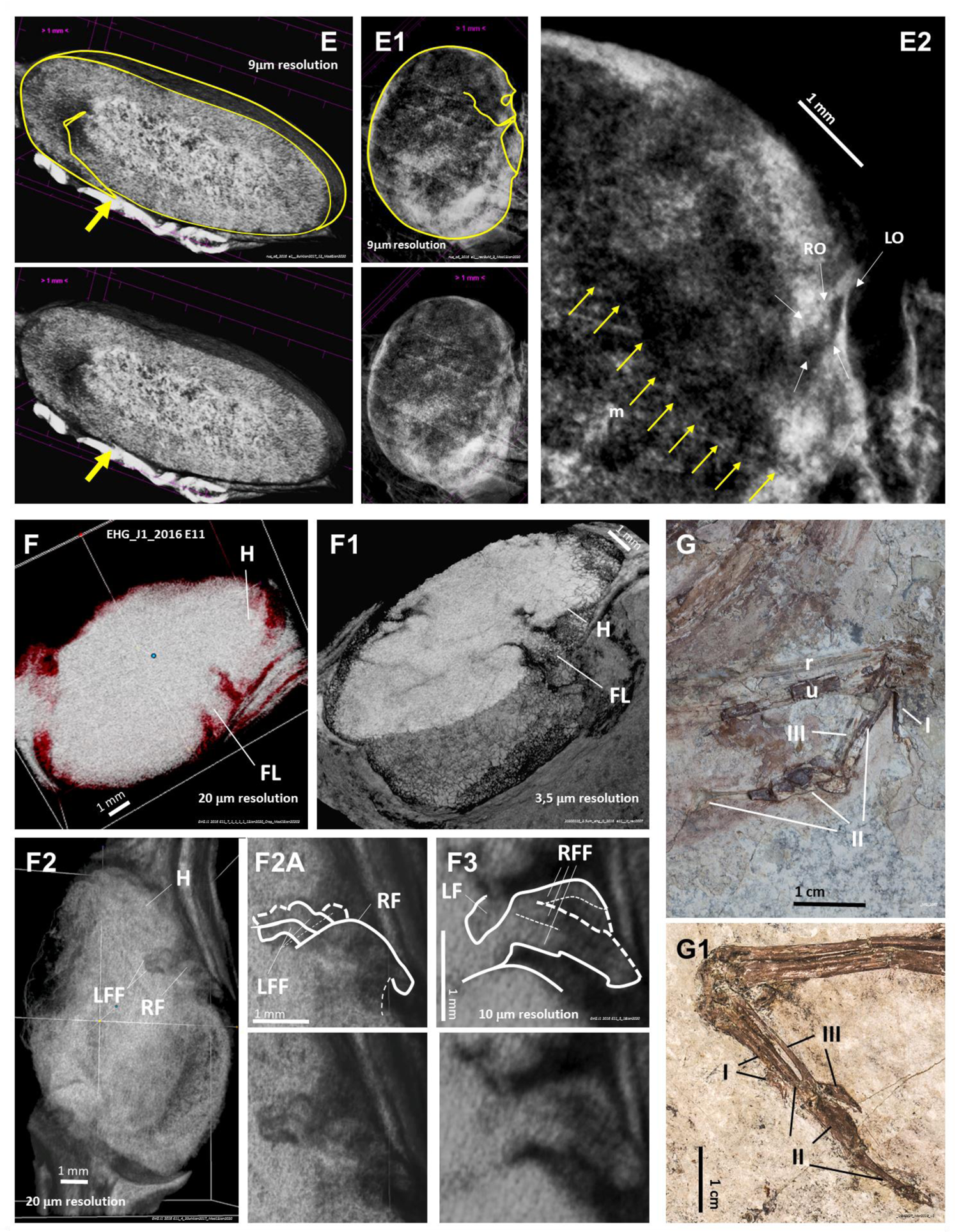
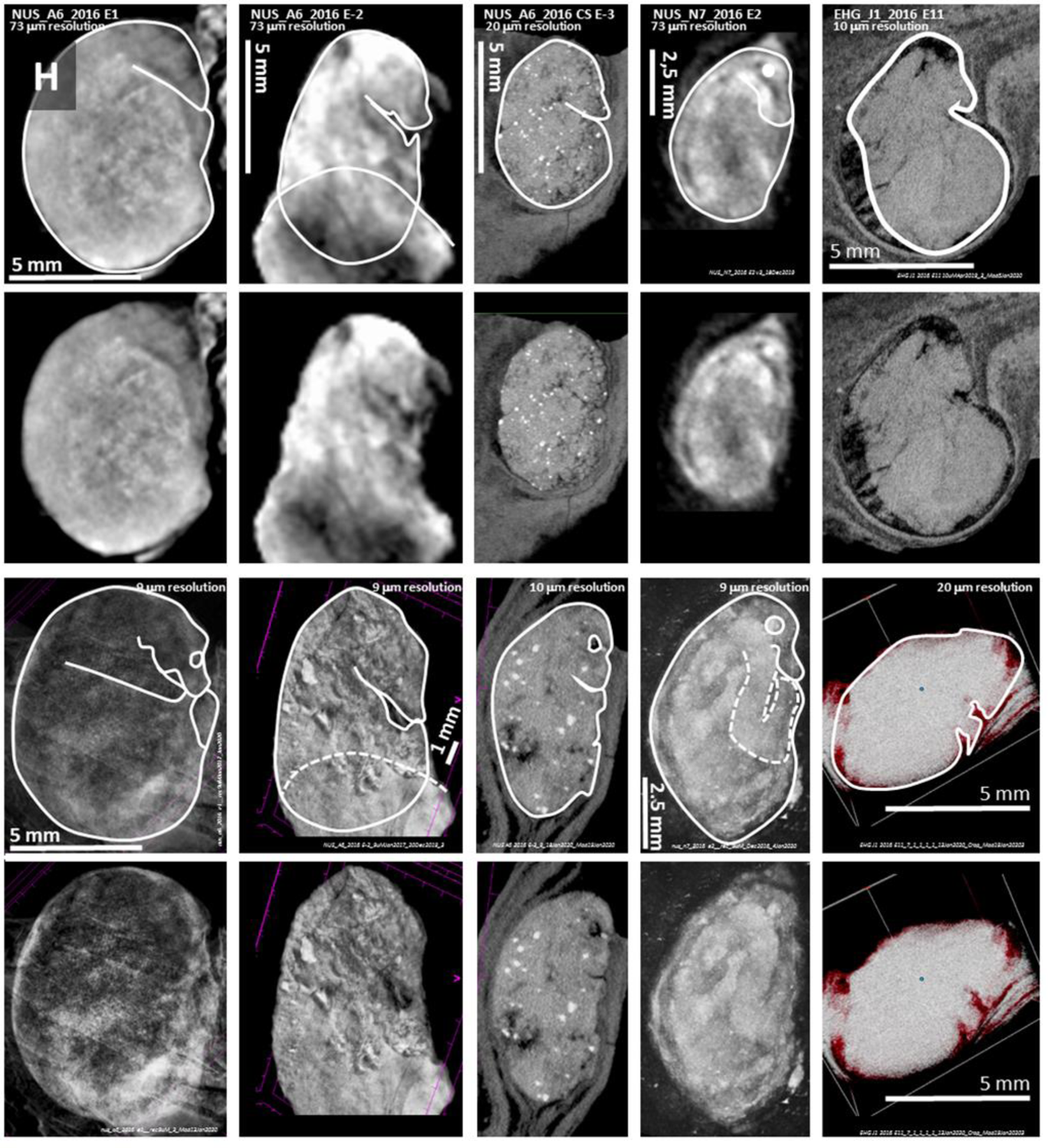
Egg periphery identification and evidence suggestive of egg-enclosed fetuses with surface morphology preservation. Virtual cuts of NUS_A6_2016 E1 extracted from specimen NUS_A6_2016 shown in Figure 1A (**A**), EHG_J1_2016 E1 extracted from specimen EHG_J1_2016 shown in Figure 1C (**B**), NUS_N9_2016 E-1 extracted from specimen NUS_N9_2016 shown in Figure 1D (**C**), and NUS_A6_2016 CS1, which, like NUS_A6_2016 E1, derived from specimen NUS_A6_2016 (**D**). Each egg has an indication of its periphery (arrows). NUS_A6_2016 E1, also shown in A, is shown under a virtual sagittal cut (**E**) and a 3D coronal view at 9 μm resolution (**E1** magnified in **E2**). The yellow arrow in E points at what appears to be a beaked fetal head outlined in E1 and amplified in E2 (yellow thin arrows point at the putative lower margin of the fetal mandible in E2; the left (LO) and right (RO) fetal orbits are also putatively indicated). The EHG_J1_2016 E11 fetus that resides within the EHG_J1_2016 egg shown in B is shown under 3D CT scanning in sagittal view at 20 μm resolution in **F**, in a virtual coronal cut at 3.5 μm resolution (**F1**), and in a 3D coronal view at 20 μm resolution (**F2**). Details of what appear to be its crossed left and right forelimbs are shown at 20 μm and 10 μm resolution, respectively, in **F2A** and **F3**. **G** and **G1:** Respectively, macrophotography of NUS_A6_2016’s left and EHG_J2_2016’s right wing. Five examples of what appears to be surface morphology preservation of egg-enclosed fetuses are shown in **H**. Eggs are shown in coronal CT view (top rows) and under 3D CT scan view (bottom rows), outlining what appear to be fetuses fossilized with only surface morphology preservation. I, II and III – 1^st^, 2^nd^ and 3^rd^ wing fingers; FL – Forelimb; H – Head of the fetus; LFF – Left forelimb fingers; LO – Left Orbit; lw – left wing; r – radius; RF – Right forelimb; RFF – Right forelimb fingers; RO – Right Orbit; u – ulna.

Importantly, despite the lack of evidence of fetal bone fossilization, the interior of the NUS_A6_2016 E1 egg appears to be entirely occupied by a beaked fetus in head tucked position with surface morphology preservation (Figures 2E-2E2, S29, and S30).

In turn, the EHG_J1_2016 E11 egg (Figures 1G and 2B), also devoid of fossilized fetal bones, seems to display surface morphology preservation not only of a fetal head in tucked position, as seen in NUS_A6_2016 E1 (Figures 2F and 2F1), but also of anteriorly crossed forelimbs, including what appear to be left and right wing fingers (Figures 2F2, 2F2A, and 2F3) that would be compatible with the 3-finger wing anatomy of adult specimens from this species (Figures 2G and 2G1, and S31-S33).

In total, five examples of what appear to be egg-enclosed fetuses with surface morphology preservation and no evidence of fetal bones are provided in Figure 2H. An additional egg that may have fossilized at the hatching stage with some degree of surface morphology preservation of the hatching animal is shown in Supplementary Material (Figures S34-S37).

Under guidance provided by micro-CT scan images, the eggshell of these eggs was confirmed via Raman spectroscopy due to its calcite signature modes at ≈156-160, ≈283-285, and ≈1091 cm^-1^, and via SEM with Electron Dispersive Spectroscopy (EDS) mapping mostly as a continuous, calcium-rich prismatic layer (see below).

As seen in Figures 3A, 3B, and S38-S41, the eggshell identified via CT scanning on the surface of two different broken quadrants of a 3D preserved egg that had been fractured into four segments was confirmed to be calcium rich under SEM with EDS mapping.

**Figure 3.**
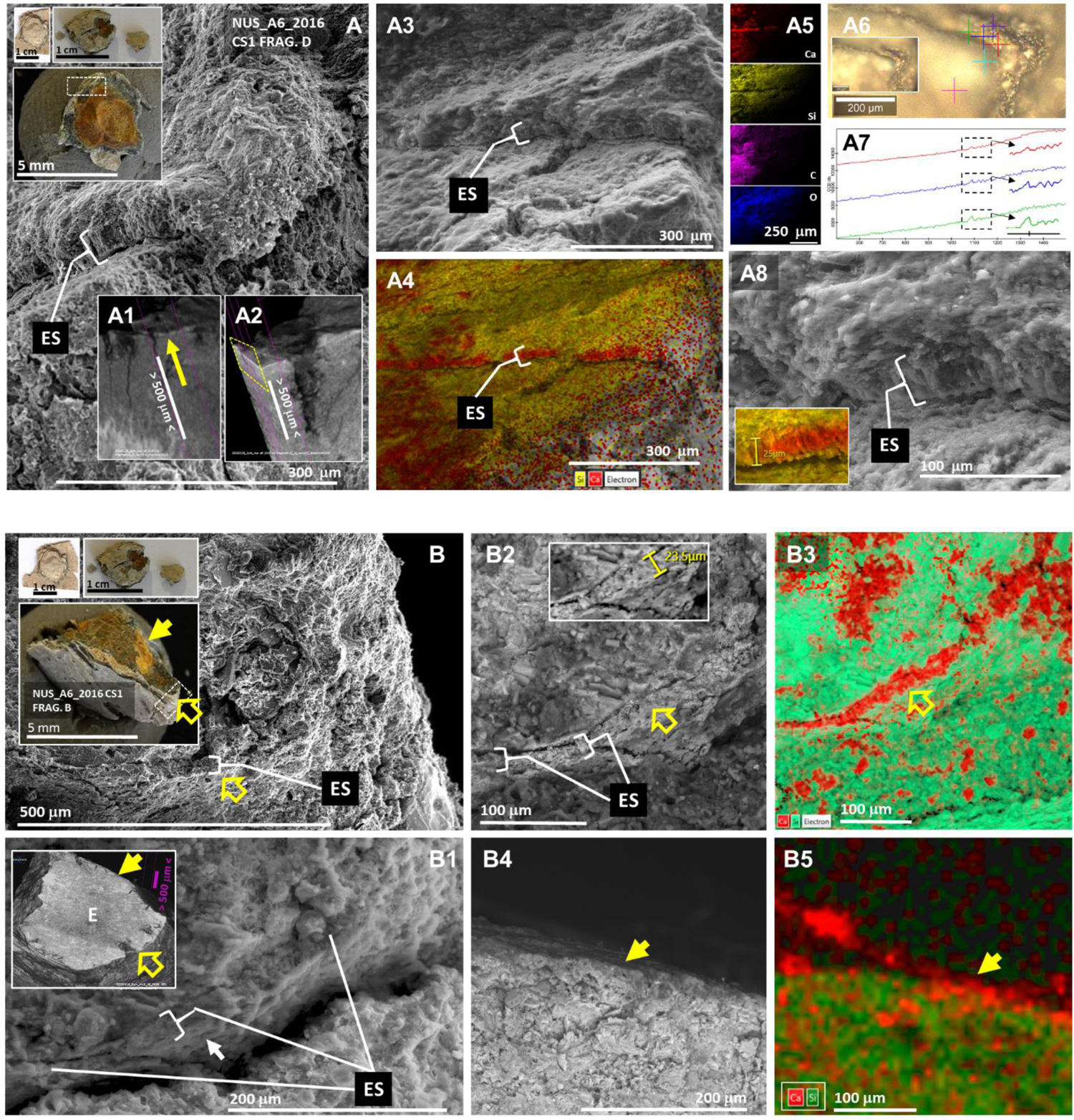
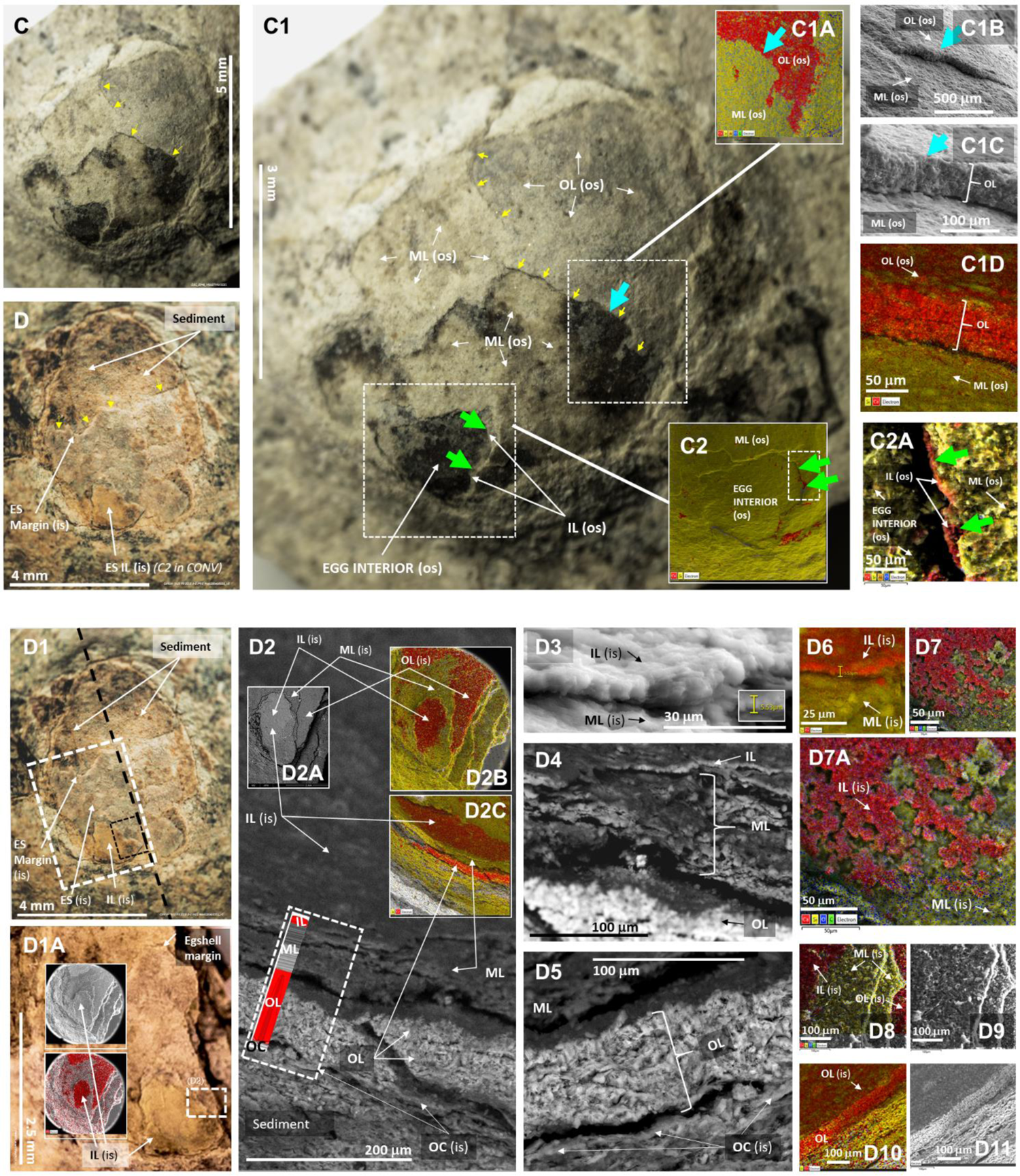
The ornithuromorph soft eggshell. **A** and **B**: SEM and EDS analysis of two quadrants of the fragmented NUS_A6_2016 CS1 egg. A: The rectangle in the lager of the upper left corner insets that show different phases of the NUS_A6_2016 CS1 egg fragmentation indicates the area of fragment D shown, respectively, under SEM and 3D views of virtual cuts of a CT scan at 3 μm resolution in **A1** and **A2**. The eggshell (ES) is indicated. The yellow arrow in A1 indicates the position at which a virtual cut was performed under CT scan analysis to expose the eggshell in perpendicular orientations, as in A2. The same area was subjected to SEM with EDS mapping (**A3-A5**) exposing the eggshell as a calcium rich layer, and RAMAN spectroscopy (**A6** and **A7;** calcite modes are shown with an amplified view at near 1 100 cm^-1^). The columnar appearance of the eggshell under SEM is shown in **A8** (EDS mapping in the inset). **B**: The rectangle in the larger inset indicates the area of fragment B where the eggshell was identified. The eggshell position indicated by the open yellow arrow is shown in a virtual cut of the egg (E) fragment under a CT scan at 3 μm resolution in the inset of **B1**, which is further exposed via EDS mapping in **B2** and **B3**. The position indicated by the yellow full arrow in the insets of B and B1 is situated on the opposite side of fragment B from that indicated by the open arrow. The former is further explored in **B4** and **B5**, respectively via SEM and EDS mapping. Green areas in B3 and B5 are silica rich. The NUS_N9_2016 E-2 eggshell is exposed on the convex egg surface and on the concave inner surface of the fragment that separated from NUS_N9_2016 E-2 after preparation, respectively shown in **C** and **D**. The eggshell edge on the convex outer surface of NUS_N9_2016 E-2, first identified via RAMAN spectroscopy (Figure S42), is indicated by small yellow arrows in **C1**. **C1A**: area of the outer surface of the eggshell and egg studied via SEM and EDS mapping in top view. The area indicated by a light blue arrow is seen in radial view in **C1B**, amplified in **C1C** and studied via EDS mapping in **C1D** (red and yellow correspond, respectively, to calcium and silica rich areas). Another area of the convex surface of the egg (**C2**), which corresponds to a depression in its surface, exposed the margin of the inner layer (IL os) of the eggshell (green arrows in C1, C2 and **C2A;** shown in red by EDS mapping in C2A). The fragment detached from the surface of NUS_N9_2016 E-2 (D) was fractured alongside the black dotted line shown in **D1** to expose its radial surface. The rectangle in D1 was amplified in **D1A** and its fractured surface is shown in radial view in **D2**. The calcified outer layer (OL), a non-calcified middle layer (ML) and a calcified thin inner layer (IL) were identified via EDS mapping (see diagram within the dashed rectangle that encompasses the outer cuticle (OC), OL, ML and IL (this dashed area is also found in Figure 4) and shown in top (**D2A** and **D2B**) and radial (**D2C**) views. The inner eggshell layer is much thinner than the OL and the ML (**D3**, **D4** and **D5**). **D6** and **D7**: respectively, radial and top (amplified in **D7A**) views of an EDS map of the IL. **D8** and **D9**: Top view of OL, ML and IL under EDS mapping and SEM. **D10-D11**: Radial view of the OL under EDS mapping and SEM.

Importantly, the eggshell was also identified on the top surface and at the edge of a thin superficial layer in a 3D-preserved egg (Figures 3C and S42-S45), but not in areas of its surface where that same layer was not present. Notably however, it was detected at the surface and edge of the same superficial layer present in the concave surface of the fossil fragment counterpart that detached from the surface of the 3D-preserved egg after preparation, precisely matching the areas that are missing in the latter (Figure 3D).

The eggshell was radially exposed in the inner concave surface of the fragment that separated from the convex egg, revealing an eggshell structure composed of the following four continuous layers (Figures 3D-D11 and S46-S48): (1) the non-calcified outer cuticle, approximately 2-5 μm thick, which is the outermost eggshell layer; (2) the outer layer, which is 20-70 μm thick and has, among other characteristics (see below), a columnar appearance derived from prismatic calcite crystals; (3) the non-calcified middle layer, which is more difficult to observe, possibly due to lack of preservation in isolated eggshells, appears to be of similar thickness to the outer layer; and (4) the inner layer, which is the innermost eggshell layer, is only 5-6 μm thick, and, like the outer layer, is calcium-rich.

Not all eggshell layers are consistently preserved, even in a given egg. When preserved, the entire eggshell thickness appears to vary from approximately 50 or less to approximately 130 μm. As shown in Figure 2D1, the inner layer was only preserved in part of the exposed inner concave surface of the NUS_N9_2016 E-2 eggshell. This layer will go unnoticed, even if preserved, unless a radial view of the eggshell is investigated with a technique such as EDS mapping.

The outer cuticle shows donut-like impressions of delineated circular areas, approximately 20 μm in diameter, with a central perforation. The outer layer appears to have an arrangement in sublayers with separation between the outer sublayer characterized by columnar prismatic calcite crystals, 20-70 μm tall and 5-25 μm wide, arranged perpendicularly to the outer surface, from one or maybe two inner sublayers of shorter crystals with globular appearance. The prismatic calcite crystals of the upper sublayer of the outer eggshell layer funnel in their inner extremity and extend through the putative globular sublayer(s) into the outer surface of the (middle) layer that lies below, where their nucleation centers appear to reside, as in the chicken and probably also in the dinosaur eggshells (Figures 4C, 4D, 4E, and S49).

**Figure 4.**
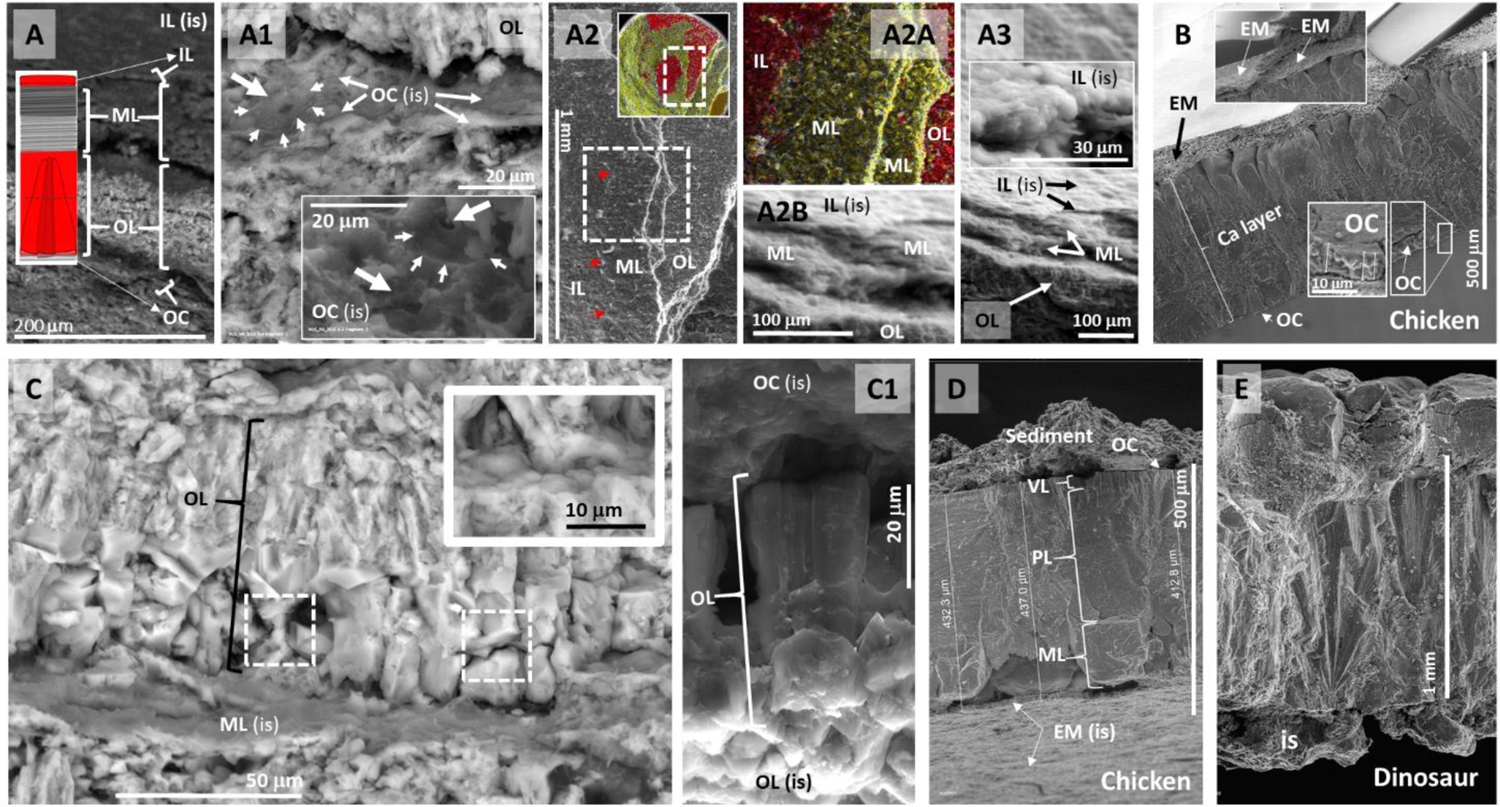
Details of the ornithuromorph eggshell. **A:** Inner (IL), middle (ML) and outer (OL) layers, and outer cuticle (OC) observable in the concave inner surface (is) of the eggshell of the NUS_N9_2016 E-2 egg observable in the fragment that was detached from the 3D-preserved egg, shown here under SEM. The diagram identifies the different eggshell layers, and represents the calcified layers in red and the prismatic crystals of the outer layer (see below). **A1**: The inner surface of the outer cuticle shows, under SEM, perforations at the center of delimited areas (small areas) that resemble a ring- or donut-like structure (thick arrows). The middle layer appears to be organized in two non-calcified layers (**A2** and **A2A**). The calcified inner layer (A2A) is only 5-6 μm thick (**A2B** and **A3**). **B**: Chicken eggshell shown for comparison, with its outer cuticle (OC), calcitic layer (Ca layer) and eggshell membranes, which are shown to detach from each other under dry conditions (inset). **C**: The outer layer of the ornithuromorph eggshell appears to be composed of sublayers. The outer columnar prismatic sublayer seems to penetrate (inset) one or two globular inner sublayers of the outer layer (dashed squares indicate equivalent areas to that in the inset). These sublayers are also apparent in a view from the inner surface of the outer layer (**C1**). **D**: Calcitic layer of the chicken eggshell shown for comparison in regards to its sublayers characterized by mammillae (ML), palisades (PL) and vertical crystals (VL)^15^. **E**: Dinosaur eggshell shown for comparison of the calcitic layer and location of the nucleation center.

In turn, the middle non-calcified layer may have two sublayers, possibly equivalent to the inner and outer eggshell membranes of the chicken eggshell (Figures 4 and S49-S53)^15^.

Under SEM, the outer surface of the outer cuticle and outer layer, as well as the inner surface of the outer, middle and inner layers are densely populated by perforations, ranging in diameter from 2-7 μm (Figures S54-S56).

Under stereomicroscopy, donut-shaped or ring-like structures, ranging from approximately 20-100 μm in diameter, with a central aperture, were observed on the outer surface of some ornithuromorph eggs, inclusively accompanied by an underlying layer with circular impressions matching the donut-shaped structures that reside above them (data not shown), but these structures were not yet confirmed under SEM and EDS mapping except, as mentioned above, in the inner surface of the outer cuticle (Figure 4A1; see Discussion).

The possibility of a nest environment was investigated in all seven *Iteravis huchzermeyeri* ornithuromorph fossils from this study (Figure S5) and other museum fossils of extinct birds from the same location beyond those from the ornithuromorph.

The area that corresponds to the soft parts and immediate body surroundings of each of the seven complete ornithuromorph specimens can be differentiated from the surrounding environment using soft tissue software settings in 3D analysis of CT scans, making the possibility of *in loco* decomposition and fossilization worthy of further investigation (Figure S16; see Discussion).

The possibility of a hidden nest environment^17^ was explored by removing sediment from the front surface of the EHG_J1_2016 fossil slab until egg detachment levels were reached, 2-3 mm below the slab surface, exposing juxtaposed layers of thin greenish material compatible with fossilized vegetation (Figures 5A-5A5 and S57 – S60).

**Figure 5.**
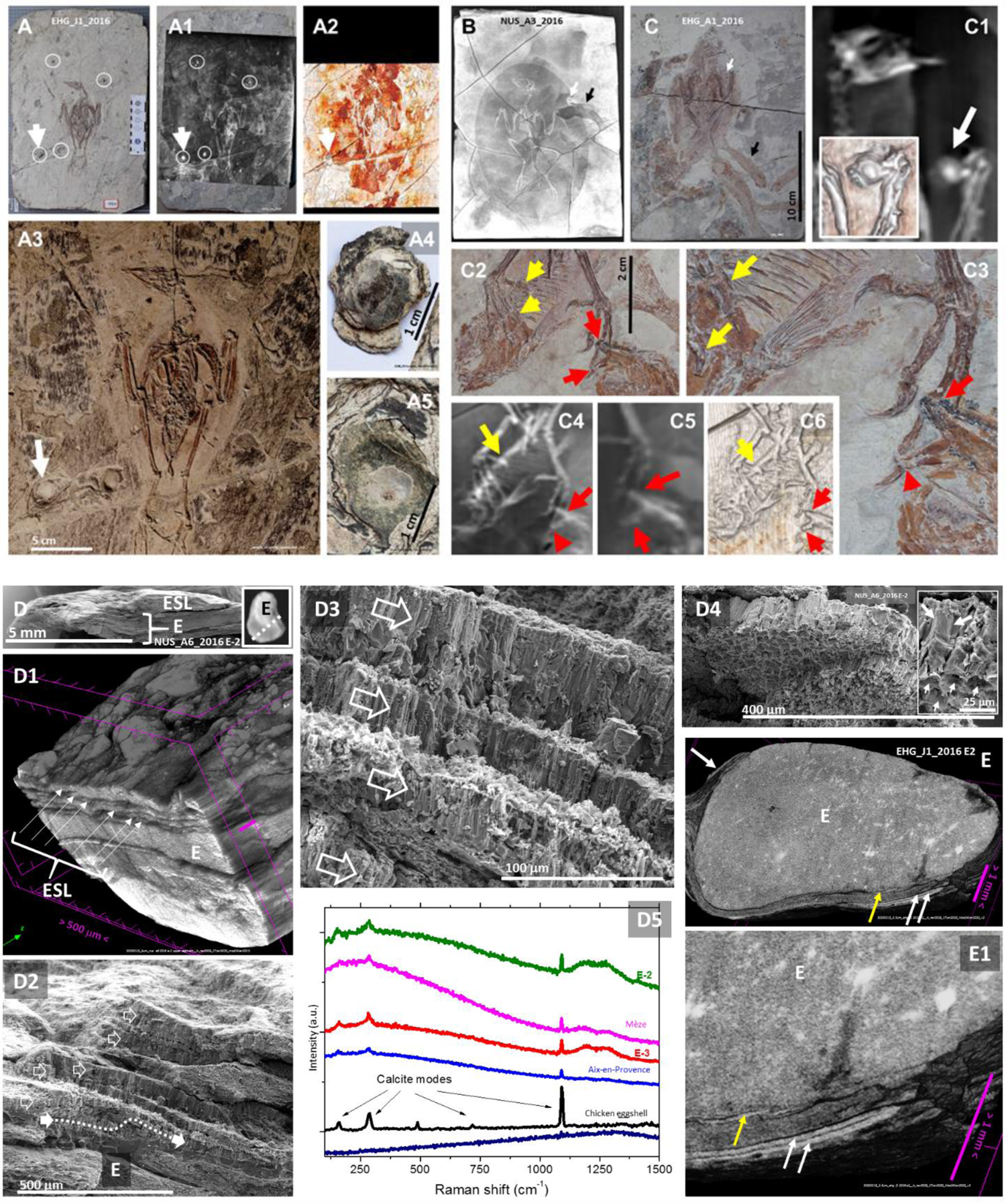
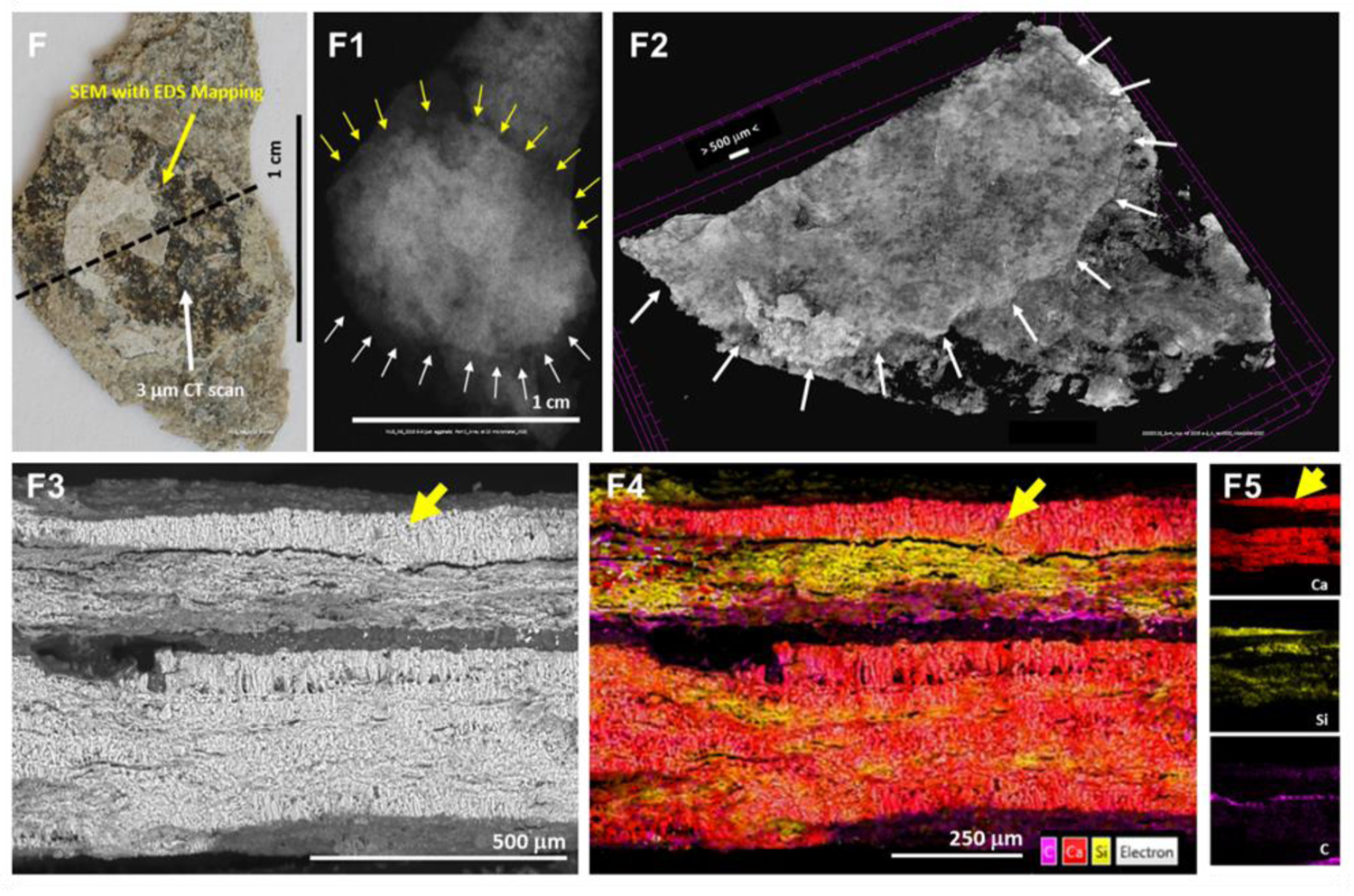
Fossilized birds and eggs atop vegetation, fish skeletons and extra eggshells. **(D-F)**. Fossil slab of ornithuromorph EHG_J1_2016 under macrophotography before preparation (**A**), under X-ray (**A1**) and 3D CT scan analysis (**A2**), and after additional preparation oriented by X-rays and CT scans to the level of egg detachment (**A3**; white arrow in A, A1 and A2 indicates egg EHG_J1_2016 E2 before its detachment; its position after detachment is shown in A3, already detached in **A4** and its fossil bed in **A5**). Enantiornithe fossils NUS_A3_2016 (**B**) and EHG_A1_2016 (**C)** are both associated with eggs (white arrows; EHG_A1_2016’s egg is shown under 2D and 3D CT scan images in **C1**; see also Figures S61 and S63-S66). Both fossilized birds are also associated with fish skeletons (black arrows in B and C). EHG_A1_2016’s left foot fingers (yellow arrows) are dorsal to a fish skeleton whereas the 3^rd^ digit of its right foot (red arrows) is ventral to another (**C2** and **C3**). A 2D CT image of the left foot (**C4**) confirms its dorsal position relative to the fish skeleton; 2D and 3D images of the right foot confirm its ventral position to another fish skeleton (**C5** and **C6**). **D**: NUS_A6_2016 E-2 egg, shown also in the inset, has extra eggshell layers (ESL) adjacent to it, which are visible under CT (**D1**), SEM (**D2-D4**; open arrows; the egg periphery is putatively indicated by the full white arrow and dotted line in D2). A RAMAN calcite signature is shown in E-2, isolated eggshell NUS_A9_2016 E-3, dinosaur eggshells from Mèze and Aix-en-Provence, and in an extant chicken eggshell, but not in a negative control (**D5**). The EHG_J1_2016 E2 egg (shown in A4) under a virtual sagittal 3D CT scan view at 3.5 μm resolution (**E**), puts into evidence how its eggshell (yellow arrow) is separated by sediment from extra eggshells (white arrows), as magnified in **E1** and confirmed by EDS (data not shown and Figures S71 and S72). An isolated eggshell, designated NUS_N9_2016 E-3 (**F**) was identified in the fossil of ornithuromorph specimen NUS_N9_2016. Its outline was confirmed via X-Ray and CT scanning (**F1** and **F2**; in the latter image after breakage along the dashed black line shown in F), and subjected to RAMAN spectroscopy and SEM with EDS mapping as shown in **F3** and **F4**. The isolated eggshell, indicated by the yellow arrow, shows as a calcium-rich layer (red in F4) separated by sediment (yellow represents silica and purple carbon) from several compressed layers of similar eggshells (thicker area of red layers interspersed with silica-rich sediment). Red areas coincide with calcite modes under RAMAN spectroscopy (see also Figures S73-S77).

Two museum fossils of unspecified enantiornithes, NUS_A3_2016 and EHG_A1_2016, also derived from Sihedang, Liaoning, and also fossilized in the same nesting position described above, were investigated and found to be associated with eggs and fish skeletons (Figures 5B, 5C-5C1 and S61-S66). The EHG_A1_2016 bird’s left foot lies dorsal to a fish skeleton whereas its right foot is ventral to another (Figure 5C2-5C6). Therefore, the bird lies in intimate association with fish skeletons, both ventral and dorsally to it. Both NUS_A3_2016 and EHG_A1_2016 have 3D-preserved eggs in or near their shoulder area (Figures S61, S62 and S64).

In addition to the association of bird skeletons with vegetation and eggs (and fish skeletons in the case of the unspecified enantiornithes), extra layers of eggshells were found, via RAMAN spectroscopy, SEM and EDS mapping, to be associated with extracted eggs and eggshells from all three ornithuromorph specimens shown in Figure 1 (Figures 5D-5F5 and S67-S77).

## DISCUSSION

### Why these findings were missed earlier

The lacustrine formation environment where these fossils were found, the small size of the eggs relative to that of the birds they are associated with, the absence of fossilized bones or other recognizable skeletal elements beyond fetal surface morphology preservation, and enhanced egg shape variation caused by compression forces upon soft eggs inherently characterized by different two-dimensional outlines in coronal, sagittal and transversal view, conjointly explain why these findings went unnoticed until now. These and other aspects will be discussed further below.

### Bird skeletons in a lacustrine environment

The ornithuromorph specimens studied here, with long and slender pedal digits, straight pedal claws and other characteristics associated with aquatic environments,^13^ constitutes such a rare example of ornithuromorph dominance in its Early Cretaceous Jehol environment that a social behavior such as seen in a breeding colony or a migration stopover had inclusively been suggested.^13^

Species holotype IVPP V18958^13^ was not studied here but is conformant to a ventro-lateral view of the same nesting posture described for seven specimens in this study. 3D software analysis of CT scan data from all seven specimens favors their decomposition at the fossilization site and not carcass drifting. The opisthotonic posture observed in three of them could have developed postmortem, either on dry land or underwater, possibly initiated from a retroflexed neck position as seen in resting and nesting birds^9,16^.

Precedents of drowned bird colonies exist, inclusively from the Mesozoic^17^. The ornithuromorph specimens studied here are sometimes seen in the same fossil slab with other individuals from the same species (personal observation). Given the presence of eggs, the cause of their death, which remains to be determined, could include low energy flooding or another event that preceded it, such as sudden death, gas released from a nearby lake, toxic algae or other causes acting upon animals that were asleep, diseased, wasted by brooding, hampered by a temperature drop or a combination of these^18^.

### Novelty of soft eggs among birds

With the exception of a report on ground nesting of soft eggs by *Archaeopteryx*, a bird ancestor^1,9–12^, no eggs have been described for any other representative of bird evolution such as *Microraptor*, *Anchiornis*, *Aurornis*, or *Confuciusornis*, and no single association of an enantiornithe bird with its eggs had been discovered until this report, unless we reinterpret as unrecognized observations of soft eggs previous reports of fossilized egg follicles, which incidentally had already been disputed for their unlikelihood (see below)^19–22^.

### Bird soft eggs

The structures identified here as eggs were first observed in fossil slabs and subsequently extracted for further study.

They lack the signs of radial growth seen in sediment concretions.

Gastralia and seeds are sometimes found in the thoracoabdominal area of fossilized birds but do not generally have a smooth surface and certainly do not have a calcitic eggshell, embryos or fetuses with surface morphology preservation^19–22^.

In turn, fossilized fruits do not have an eggshell, tend to have a consistent 3D shape and their surface is not simultaneously smooth and deprived of indentations as that of an egg.^22^

They are not fossilized egg follicles either because the latter neither have a calcitic eggshell nor embryos or fetuses with surface morphology preservation (more on previous reports of egg follicles below)^19–22^.

These eggs were originally soft because they have consistent size and no signs of breakage but show diverse 2D outlines and 3D shapes. Shape variation did not result from discontinuities of the eggshell prior to fossilization as this would have resulted in loss and destruction of the egg content, making fetal surface morphology preservation unlikely. In contrast, a continuous and discrete outline is identifiable under high-resolution CT scanning, which delimits their periphery.

Some of the ornithuromorph eggs shown here seem to display, in their interior, the outline of a fetus with surface morphology preservation, similarly to what had been reported in *Archaeopteryx* soft eggs^9,11,12^.

Further to their close association with an adult bird, the avian origin of these eggs is suggested by what appear to be beaked fetal heads with surface morphology preservation and elongated fingers, also with surface morphology preservation, similar to the 3-finger anatomy of the adult animals they are associated with (Figure 2).

Following detection by CT scans at 3-10 μm resolution, the ornithuromorph eggshell was confirmed by calcite signature modes under RAMAN spectroscopy and observed as a continuous multi-layered structure under SEM coupled with EDS mapping. The latter technique allowed the detection of its most consistently preserved component—the outer calcitic layer—on the surface of a 3D preserved egg, in two separate quadrants of an intentionally broken egg and in an isolated eggshell.

From its outer to its inner surface, the ornithuromorph eggshell is composed of (1) a membranous non-calcified outer cuticle 2-5 μm or less thick, which is homologous to the chicken outer cuticle but where perforations surrounded by a ring or donut-shaped area were observed in its inner side, (2) an outer 20-70 μm calcitic layer that appears to include prismatic and globular calcite crystal sublayers, which is equivalent to the calcitic layer of the chicken eggshell, (3) a non-calcified middle layer that may have two sublayers, possibly equivalent to the inner and outer eggshell membranes of the chicken eggshell and, finally, (4) a 5-6 μm thick internal calcified layer, which is intriguing for not seeming to have, as a calcified structure, correspondence in extant bird eggs, but may be a modified version of the homogeneous coating of the inner membrane of today’s bird eggshells. The function of this thin calcified inner layer is unclear, but could have served as an initial source of calcium during embryonic or fetal development without weakening the thicker but still relatively thin calcitic outer layer. The latter may have served as source of calcium closer to the emergence of the chick^23^. The fact that this type of inner calcified layer has not been described in dinosaurs^24^ could be due to lack of preservation, could have been missed for being very thin, or simply did not exist in those animals. It would also have gone unnoticed here if it were not for the use of EDS mapping as it is otherwise almost indistinguishable from the middle layer under SEM.

The total thickness of the complete ornithuromorph eggshell appears to vary from approximately 50 to 130 μm.

Perforations were observed in the ornithuromorph eggshell that are consistent with the openings of pores. They have similar diameter but are more numerous than those of chicken eggs^25^.

Eggs with good preservation of the eggshell and fetal bone preservation in their interior would be ideal for study, but it is unclear if they will become available in the future. Additional ultrastructural studies^23,26^ of the ornithurmorph eggshell may help determine the trajectory of eggshell pores and how the donut- or ring-shaped structures observed in the inner side of the outer cuticle translate into its outer surface and if these match a similar arrangement in the outer calcitic layer or in any way extend into the interior of the eggshell. The relative displacement of ring-like structures in relation to their vicinity, perpendicularly to the eggshell surface, could account for eggshell deformability despite its calcitic content^9,12^. This type of mechanism would also apply to shell units recently described in dinosaur leathery eggs^24^.

Additional studies are necessary in well preserved enantiornithe eggshells, but the enantiornithe eggs, having variable 3D shapes, a size range similar to that of ornithuromorph eggs, and thin eggshells (approximately 100 μm or less; Figures 5B, 5C, and S61–S66, and data not shown), allow for a more plausible reinterpretation of previous reports of fossilized enantiornithe egg follicles in the thoracic region of adult birds as those of soft eggs (see below)^19–22^.

In brief, given their high 3D shape variability, size consistency, and thin eggshell, the ornithuromorph eggs were soft. Therefore, not only *Archaeopteryx*, an earlier bird ancestor^9–12^, but also ornithuromorphs reproduced via soft-shelled eggs.

### Clutch size

The exact number of eggs of the ornithuromorph clutch, for which several specimens were available for study, was probably in the tens, judging from eleven 3D-preserved globular structures and several additional impressions noted in NUS_A6_2016 (data not shown). It is not possible to estimate the clutch size of the enantiornithes shown in Supplementary Material because only one of their slabs was available.

### Exceptional fossil preservation

The surface morphology preservation of fossilized fetuses observed here is consistent with a high degree of preservation, inclusively 3D preservation, already previously noted among mature birds where these fossils were found^13^.

Hatchlings were searched for in fossils from this location but were not conclusively identified. The higher radiodensity of the eggs in the absence of fetal skeletal preservation, even in the case of the egg for which forefinger outlines were detected, suggests late ossification of the developing ornithuromorph organism. Alternatively to late ossification, lack of skeletal preservation might have resulted from dispersal of bone elements during a demineralization phase of the fossilization process. Calcium and other radiodense elements may have been adsorbed by matrix components of fetal soft tissues before re-mineralization occurred, which, together with other factors responsible for outer morphology preservation, would have resulted in fetal body surface demarcation. Research on exceptional fossil preservation and its mechanisms^27,28^ will gain from further investigation of these fossils.

### Ground nesting of soft eggs in a peri-aquatic environment

Eggs were found in association with several ornithuromorph specimens. The finding that the area that corresponds to the soft parts and immediate body surroundings of each of the seven complete specimens that were studied here can be differentiated from the surrounding environment using CT scan software settings of soft tissue is compatible with their *in loco* decomposition and fossilization.

The finding of extra eggshells in two ornithuromorph fossils suggests repeated usage of the same ground nesting site. Accordingly, examples exist of several adult birds in the same fossil slab, just a few decimeters apart from each other (personal observation; data not shown), suggesting colonial ground nesting. Furthermore, the close association of enantiornithe fossils with eggs, vegetation, and fish — one specimen’s left foot is dorsal to a fish skeleton, whereas its right foot is ventral to another — raises the possibility that fish were brought to the nest for feeding and were not the mere result of their joint sedimentation with sunken bird carcasses that had been floating at the bottom of a lake.

A similar discovery of colonial ground nesting in an aquatic ecosystem was reported for *Archaeopteryx* in the Jurassic Solnhofen marine environment^12^. In addition, there are also reports of dinosaur nests on tidal flats^29,30^ and periaquatic colonial nesting by pterosaurs^5^.

The thin eggshell of the ornithuromorph identified here would call for high humidity to avoid egg desiccation, as seen near shore. In turn, protection from sunlight would have been provided by the feathered and winged body of the adult animal. It is therefore probable that these birds nested near shore and that events led to their death and burial by lake waters.

### Reinterpretation of previous reports on fossilized egg follicles and other egg-related findings in ancient birds

This discovery calls for reinterpretation of several previous reports. These include similarly sized but differently shaped egg follicles located in the thoracic area of enantiornithes^19,20^, which are unlikely characteristics for egg follicles for their size consistency and body location, aside from their unlikely fossilization^21^. Instead, these findings are better explained by soft eggs.

The description of an un-laid enantiornithine egg with several eggshell layers^31^ is more likely to correspond to an egg atop extra eggshell remnants.

In turn, an egg-enclosed avian embryo from the Early Cretaceous of China that does not show an eggshell^32^ could represent a soft egg with no eggshell preservation or for which the presence of a thin, 100 μm or less, eggshell was not thoroughly investigated.

Paleo-environments such as the Yixian Formation in China, where numerous birds but no eggs have been found, need revisiting for the possibility that similar findings to the ones described here might have been missed in previous studies.

### Additional implications of soft eggs for birds and other animals

The ontogeny and phylogeny of birds will have to start taking into consideration, among other currently evaluated characteristics, reproduction via hard or soft-shelled eggs.

Descriptions of microvertebrates and small, including embryonic, dinosaur material in the absence of eggshells^33–37^ also need to be revisited for the possibility of soft eggs.

Some dinosaur lineages for which eggs were never found, such as tyrannosaurids, ornithomimosaurs, and ankylosaurs^38^, may have laid non-rigid eggs^9,39^ as reported for *Protoceratops* and *Mussaurus*^6^.

It is increasingly likely that an egg-associated Compsognathus from Solnhofen may have reproduced via soft eggs^9,12,40^.

### Implications in wing and flight evolution

The finding reported here of early birds nesting soft eggs on the ground adds to a similar report in *Archaeopteryx*^9,12^, reinforcing the possibility that elongated feathers may have developed primarily in association with nest protection on the ground and only secondarily with flight^9,12,41^, and favors the origin of flight from the ground up^12,42^.

### Alternative explanation to convergent evolution for the existence of soft and hard shelled eggs in birds

In this report, we just showed examples of early birds that reproduced via soft eggs, whereas all extant birds reproduce via hard-shelled eggs. The occurrence of eggshell type variation (soft or rigid) within different animal groups — geckos, turtles, pterosaurs, and dinosaurs — is commonly attributed to their independent and repeated evolution, also referred to as convergent evolution^6^. Given that crocodiles reproduce via hard-shelled eggs and precede birds in evolutionary terms, convergence would normally also be invoked here to explain the discovery of non-rigid eggs in the bird lineage.

However, a more parsimonious explanation in alternative to the repeated evolution of rigid eggshells among birds will be proposed here. As we embark on such an explanation, given the coexistence of hard and soft-shelled eggs in a variety of animals, a wider discussion is necessary. Moreover, some reptiles (namely, snake and lizard species) are capable of laying soft eggs while others are viviparous, for which reason a broader analysis of intra-group parity mode polymorphism (the capacity to use more than one reproduction mode in a given animal group) needs to be addressed, too.

The expression of oviparity and viviparity within the same animal group is actually ubiquitous. Far preceding amniotes, it is apparently as old as the first animals. Going backwards in evolutionary terms, it is found not only in non-amniotic vertebrates like certain fish^43^ and sharks^44^, but also in invertebrates like flies^45^ and thirps^46^, velvet worms^47^, flatworms^48^, and even sponges^49^. Groups of amniotes with oviparous and viviparous species include mammals, snakes, lizards, amphibians (frogs, salamanders, and caecilians)^50^, and possibly also marine reptiles^7,8^. Parity mode variation among squamates is such that convergence is commonly believed to be responsible for viviparity to have evolved at least 115 times in this animal group, the most among the approximately 150 times it would have evolved in vertebrates, including 9 in cartilaginous fish, 13 in bony fish, 8 in amphibians, and 1 in mammals^43, 46,48^.

What is more, from sponges to mammals, if variance between oviparity and viviparity is not seen within a given animal group, which will then be a strictly oviparous group, some of its animals will lay soft-shelled eggs while others lay hard-shelled eggs. The only known exception to this rule are crocodiles, which lay rigid eggs only.

The inherited capacity to switch between oviparity and viviparity is hypothesized here as a parsimonious alternative to the convergent evolution of one or the other parity modes. Such a parity switch is hypothesized to be present since the rise of animals and to have been inherited from a common ancestor to all animals, extending forward into and throughout the evolution of the amniotic egg (Figure 6).

**Figure 6.**
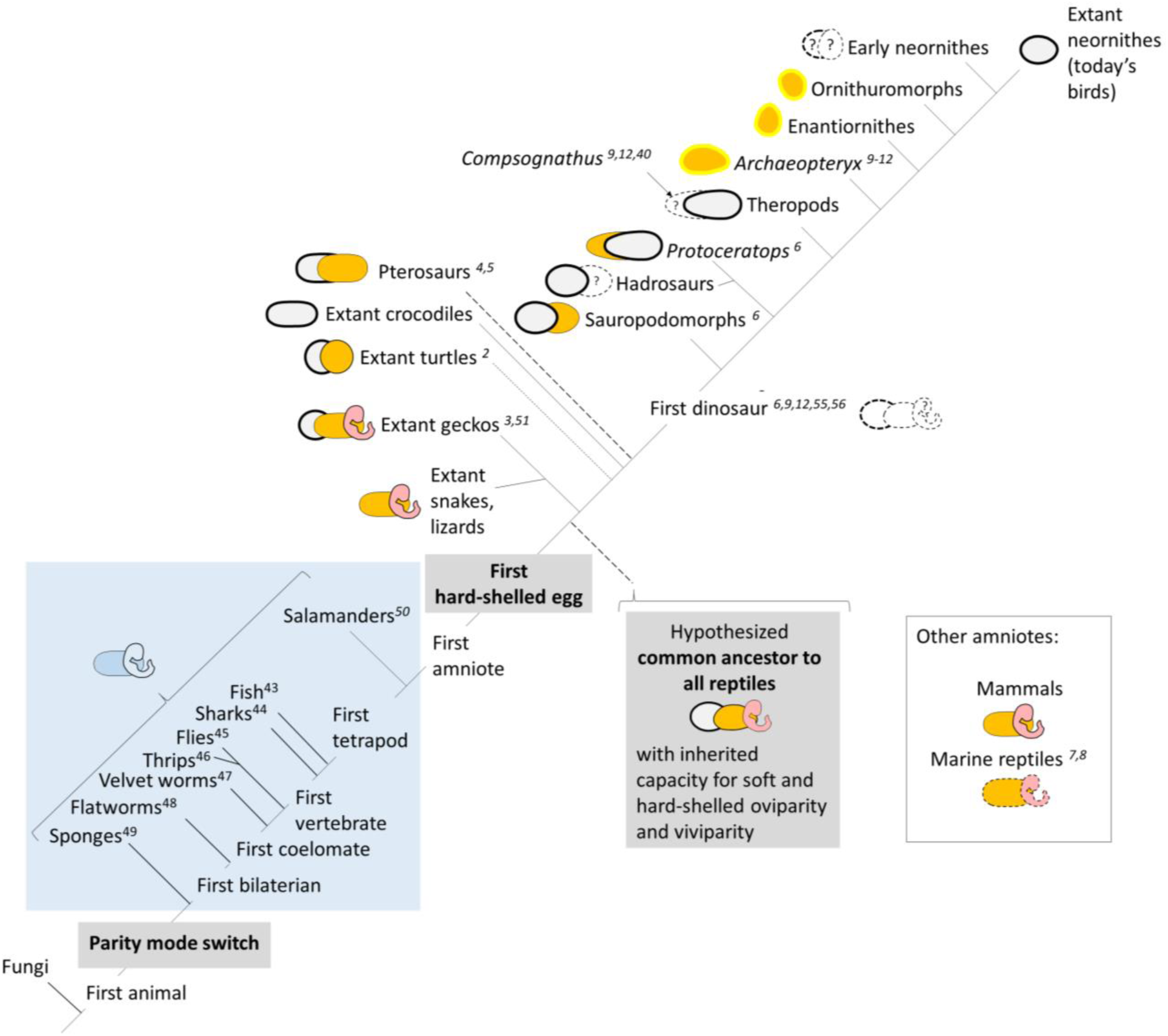
Parity mode switch hypothesis of non-convergent reproduction evolution in all animals. Viviparity is represented by an embryo, soft eggs are represented in orange, and hard-shelled eggs in light gray with a thick outline. The pre-amniotic stage is shown in blue background. Unverified possibilities are represented by dashed lines and what is unknown is represented by dashed outlines associated with a question mark. References are shown in superscript. Results for the enantiornithes and ornithuromorphs reported here and for *Archaeopteryx* reported separately are outlined in a thick yellow^9–12^. Dinosaur phylogeny as in Baron *et al*.^55^ The possibility that *Compsognathus* reproduced via soft eggs is suggested^9,12,40^. It remains to be determined if early neornithes reproduced via soft and/or hard-shelled eggs. The uncertain origin of pterosaurs is reflected by dashed lines. Mammals and marine reptiles are also encompassed by the parity mode switch hypothesis but will be analyzed separately (dashed outlines in marine reptiles’ reproduction strategies reflect reported results that will be revisited separately).

The parity switch may have conferred a selective advantage and may have evolved independently of the parity modes, as seen, for example, with the diversification of eggshell features in oviparous animals and the emergence of various degrees of newborn maturation in viviparous life forms.

Much later in evolution than the arrival of the parity switch between oviparity and viviparity, a single occurrence of *de novo* evolution of hard-shelled eggs in the common ancestor to reptiles and birds is assumed here to have taken place. After the appearance of hard eggshells in animals, the parity switch originally capable of alternatively engaging oviparity or viviparity would have evolved to extend its capacity to include oviparity modalities, enabling oviparous animals to switch between soft and hard-shelled eggs.

The expression of viviparity may have been lost before evolution of testudines or may have extended well into within the dinosaur lineage^50^, a not so remote possibility given that soft-shelled eggs were already reported in this animal group^6,9,12^. In this regard, the possibility that *Compsognathus*, a theropod, reproduced via soft eggs was proposed and is again suggested here^9,12,40^, and the possibility of viviparity among dinosaurs should gain renewed interest with this new parity switch hypothesis.

Consistent with the parity switch hypothesis put forward here, joint oviparity and viviparity were recently described within the same clutch of an individual extant female skink^51^. Throughout evolution and also in today’s animals, this trait may possibly have been initially expressed in the same pregnancy, within different seasons of the reproductive life of any given individual organism, or even among different members of a given species, be it within the same or in different generations of such species, and, over time, may have become more commonly expressed in a species-specific manner, or it was since early on associated with speciation mechanisms, as is certainly the case today for most land vertebrates, except for some squamates.

This hypothesis of co-evolution of a switch mechanism with retention of silenced alternatives is in conflict with the commonly accepted independent and repeated (i.e., convergent) evolution of parity modes and explains the occurrence of rare parity reversals that are actually predicted by phylogenetics^52,53^. These apparent reversals are proposed here to be simply the result of an alternation between preexisting capacities due to retention and recruitment, not because of new evolution after a true loss that had resulted in regression.

The parity mode switch is most likely associated with regulatory systems with ample effects in animal biology, such as immunity, repair, and blood vessel generation, which, beyond their requirement to assist in reproduction, have such importance to animal life forms that are obligatorily transmitted throughout evolution since the rise of animals, preceding even the development of organs, as denoted by the fact that sponges also display reproduction bimodality. This is in agreement with recent data on functional genetics of oviparity and viviparity^54^.

It remains to be determined if early neornithes reproduced via soft and/or hard-shelled eggs. Regardless, reproduction via soft eggs was present for longer than previously imagined in the evolution of birds, possibly until the extinction of enantiornithes at the end of the Cretaceous, and may also have persisted among pterosaur and dinosaur species until their extinction (Figure 6) ^4–7,55,56^.

### Candidate mechanisms involved in the ability to switch between soft and hard-shelled oviparity

Following formation of the inner eggshell membrane, which precedes eggshell calcification, candidate mechanisms for the determination of eggshell type include heterochrony^57–59^ and activation of alternative pathways that influence oviduct processes, with or without metabolic cues from the developing fertilized egg^9^.

### Pathway to the reproduction strategies of todays’ birds and reptiles

According to the hypothesis put forward here, reproduction in extant reptiles and birds are manifestations of inherited reproduction capacities that are now generally under speciation mechanisms. Unlike crocodiles and birds, which only lay rigid eggs, snakes and lizards would have preserved viviparity and soft egg parity, whereas the capacity to reproduce via soft or hard eggs would have persisted in turtles.

However, speciation mechanisms may not have irreversibly silenced the ancestral facultative parity feature in reptiles, as the already mentioned example of a skink with oviparity and viviparity within the same clutch may be manifesting^51^. If this is the case, other examples of individual, generational, or intra-group species variation in reproduction strategy within reptiles and manifestations, however unsuccessful, of the same mechanism in other animal groups, may be discovered in the future.

Despite occasional soft bird eggs associated with nutritional and environmental stress in today’s birds^60^, only bird species committed to reproduction via rigid eggs survived. One wonders, however, if an inherited capacity to reproduce via soft eggs is now irreversibly lost or could reappear in new bird species under appropriate conditions.

The hypothesis of non-convergent reproduction evolution was presented here to explain the existence of hard-shelled eggs in the descendants of a generally accepted common ancestor to reptiles, turtles, pterosaurs, marine reptiles, dinosaurs, and birds. The evolution of reproduction in marine reptiles and mammals was also encompassed by the parity mode switch mechanism, but the possible impact of this hypothesis in these animal groups will be discussed separately (manuscripts in preparation).

### Implications in the extinction of major bird groups and other animals

Today’s birds reproduce only via rigid eggs, which are believed to have a higher capacity to respond to environmental stress. Soft-shelled eggs may provide a new explanation for the extinction of major bird groups if eggshell structure was key to environmental adaptation and the capacity to commute between eggshell types was lost or conditions were not permissive to eggshell type variation within a few generations, as could have suddenly occurred at the end of the Cretaceous Period, when all enantiornithes underwent mass extinction^61^.

Eggshell type may also have contributed to the extinction of, among others, dinosaurs, pterosaurs, and marine reptiles at different moments, including at the end of the Cretaceous Period. However, in the case of these animal groups, because other reproduction strategies have also been reported for them^4–8^, additional factors must have been at play because all dinosaur, pterosaur, and marine reptile species perished, not just those that would have reproduced via soft eggshells. However, an abrupt decrease in their diversity, caused by the loss of soft egg-laying species in these animal groups, could have precipitated their mass group extinction.

## CONCLUSIONS

Extinct fossilized birds from the early Cretaceous — specimens of ornithuromorph *Iteravis huchzermeyeri* are shown here in nesting posture together with eggs of similar size but variable 3D shape, atop vegetation and extra eggshells.

The ornithuromorph eggs are 0.6 to 1.0 cm in their longer axis, comparable to today’s smallest hummingbird eggs and smaller than the 1.3 cm long eggs reported in *Archaeopteryx*. Among other examples suggestive of surface morphology preservation of egg-enclosed ornithuromorph fetuses with beaked heads, one is shown with what appear to be elongated wing fingers as seen in the adult animal.

The ornithuromorph soft eggshell is a continuous peripheral layer observable under CT scanning at 3 – 10 μm resolution, shows calcite signature modes under RAMAN spectroscopy and a multi-layered structure under SEM and EDS mapping comprised of a 2-5 μm outer cuticle, an outer calcitic layer, 20-70 μm thick, which is the layer most often preserved, a non-calcified middle layer, which appears to tend to be thinner than the outer layer (up to 50 μm thick) but is less frequently preserved, and a thin (5-6 μm) inner calcified layer. In total, it is 20-130 μm thick, depending on the degree of preservation of all its layers, a thickness somewhat comparable to the thinnest soft eggs from extant geckos^26^.

Nearshore ground nesting would provide the humidity required for soft egg incubation, explaining the intriguing abundance of bird fossils, including ornithuromorphs and enantiornithes, in lacustrine environments.

The novelty of non-rigid eggs in birds requires explanation because both crocodiles, whose ancestors antedated the animals that eventually evolved into bird precursors, and today’s birds reproduce exclusively via hard-shelled eggs.

All major animal groups, from sponges to mammals, with the apparent exception of crocodiles, either comprise species that are oviparous while others from the same group are viviparous or aggregate exclusively oviparous species, but some lay soft-shelled eggs while others lay hard-shelled eggs.

A novel parity mode switch hypothesis of non-convergent evolution of reproduction was formulated here as an alternative to the commonly accepted convergent evolution of viviparity and rigid eggshells. This hypothesis claims the existence, since the rise of animals, of an inherited ancestral parity mode switch between viviparity and oviparity, which was extended to also embrace hard-shelled oviparity after rigid eggshells appeared in evolution and, under circumstances and mechanisms to be identified, allows not only for retention but also recruitment of the silenced parity mode or eggshell type despite parallel diversification, via evolution, of viviparity and egg features in the various animal groups.

It is also proposed that stern restriction to a particular parity mode or eggshell type may condition survival of entire animal groups, especially during major extinction events. This may have been the case with enantiornithes and ornithuromorphs if these only reproduced via soft eggs.

## MATERIALS AND METHODS

### Fossils

The NUS_A6_2016, NUS_N7_2016, NUS_M1_2019, NUS_A3_2016, EHG_A1_2016, AX_A1_2016 and BRU_1 fossils are kept at the Ningcheng Administration for National Geopark, Ning Cheng, the EHG_J1_2016 and EHG_J2_2016 fossils are kept at the Sichuan Chongzhou Tianyan Museum, and the NUS_N1_2016 and NUS_N9_2016 fossils are kept at the Inner Mongolia Natural Museum. The specimens were brought to the respective museums, catalogued and made accessible for study after their discovery in 2010, except for BRU_1, which was discovered earlier, around 2001. The latter is currently on loan to the Royal Belgian Institute of Natural Sciences.

Eggshell samples from spherical fossilized eggs approximately 15 cm in size (Cairanoolithus) from the Campanian Upper Cretaceous period of Mèze, France, discovered in 2013-2014, were kindly provided by François Escuillié, Eldonia, France.

### Fossil search

A search for fossils of bird precursors that fossilized in a pose similar to that described for *Archaeaopteryx*^*9*^ led first to the identification of museum fossil NUS_A6_2016 from the locality of Sihedang, Liaoning, China. Other fossils of the same species were identified next as well as those from unspecified enantiornithe fossils.

MJG visited with co-authors the location of the Early Cretaceous Formation where the ornithuromorph fossils were found, in Sihedang, Lingyuan, Liaoning, and observed quarries where numerous fossils with similar sediment characteristics had been found.

### Species identification

The identification of the ornithuromorph bird was as described^13,62^ via association of a predentary bone, U-shaped furcula, expanded first phalanx of the major digit and short plow-shaped pygostyle with premaxillary corpus elongate and toothless, numerous teeth in the maxilla, rostrum that represents 50% of the skull’s length, ethmoid bone lining rostral to half of the orbit, tubercle on caudal margin of minor digit phalanx, pubes with dorsally expanded distal boot, and narrow ischium with concave ventral margin and weak dorsal process at midpoint.

### Extraction of samples

Macrophotographs, microphotographs, X-rays and 0.5 mm resolution CT scans were used for egg localization in the fossil slabs and also throughout egg extraction, which was performed under stereomicroscopy, between 2016 and 2019, at each museum, where some of the fossils are on permanent exhibition (such as EHG_J1_2016 and EHG_J2_2016) and others in their Collection Rooms.

Extracted eggs were individually documented under macro- and microphotography and labelled using (1) the name of the fossil followed by E or CS (depending on whether the egg was extracted from the main slab or the counterslab), (2) the minus sign if the egg was extracted from the back of a slab and (3) a number (originally initially assigned to corresponding radiodensities identified in X-rays).

Eight of eleven globular structures detected by surface and radiologic analysis of the NUS_A6_2016 fossil (NUS_A6_2016 E1, E2, E-2, E-11, CS1, CS-3, CS10 and CS11), five in the EHG_J1_2016 fossil, three in EHG_J2_2016, six in NUS_N1_2016, three in NUS_N7_2016 and five in NUS_N9_2016 were individually fossil extracted under stereomicroscopy, using X-ray and CT scan images as guidance. NUS_M1_2019 was left aside in its original condition for future studies.

### Photography

Photography of each fossil slab was done using a tripod in different angles and light conditions using the following cameras: a Canon 5D Mark II, a Canon EOS 5D Mark II or a Canon EOS 5D Mark IV. The first two were associated with an AF-S DX NIKKOR 18-105 mm f/3.5-5.6 GD VR lens. All were associated with either a Canon EF 24-105mm f/4 L IS USM lens, a Canon EW-88D 16-35mm ULTRASONIC lens, a Canon Macro Lens EF 100 mm f/2.8 USM, a Canon Macro Lens EF 100mm 1:2.8 L IS USM or a Canon Macro Lens EF 180 mm 1:3,5 L ULTRASONIC.

### Stereomicroscopy

Stereomicroscopy was done under a Nikon SMZ1270 stereomicroscope associated with a Photonic Optics LED control unit as a light source and mounted on a Motic horizontally articulated arm. The stereomicroscope was attached to a Nikon D5500 digital camera, which captured JPEG and RAW files under manual setting using a 20 second delay for picture shooting.

### X-rays and CT scans

X-rays were obtained of each slab from each fossil using several types of medical equipment, both film-based and digital, and CT scans of each slab from each fossil were initially obtained using several medical-degree CT scanners at 0.5 mm resolution that generated digital files in DICOM format.

The extracted ornithuromorph eggs were studied, from 2016 through 2020, using the following high-resolution CT scanners: a Mediso SPECT/CT Scanner at 73 μM resolution and a Mediso NanoScan SPECT/CT Scanner at 11 μm resolution (Molecular Imaging Department of Chung Gang Memorial Hospital, Taoyuan, Taiwan), a Milabs U-CT Scanner at 20 and 10 μm resolution (Yang Ming University, Taipei, Taiwan), a SkyScan 1076 from Bruker CT Scanner at 9 μm resolution (Animal Center at Chung Gang University, Taoyuan, Taiwan), a DELPet μCT 100 at 9 μm resolution (Taiwan Mouse Clinic, National Biotechnology Research Park, Taipei, Taiwan) and a SkyScan 1276 from Bruker at 3 μm resolution (Taiwan Mouse Clinic, National Biotechnology Research Park, Taipei, Taiwan).

The BRU_1 fossil was examined on a RX Solutions CT Scanner at 59,59 μm resolution (Royal Belgian Institute of Natural Sciences).

The egg cluster extracted from NUS_A3_2016 was studied under a Milabs U-CT Scanner at 10 μm resolution (Yang Ming University, Taipei, Taiwan). The egg extracted from unspecified enantiornithe fossil AX_A1_2016 was studied under a SkyScan 1276 from Bruker at 3 m resolution (Taiwan Mouse Clinic, National Biotechnology Research Park, Taipei, Taiwan).

### 3D Software analysis of CT scan data

CT scans in DICOM, TIFF, nii or BMP formats were analyzed with Pmod^®^, 3D Slicer^®^ version 4.6.2 r25516, CTVox^®^ and Vivoquant^®^. 2D and 3D images from CT scan software analysis were saved in BMP or JPEG format. CT scan data above 3 μm resolution was explored with an ASUSTek^®^ computer (GL502VS model, with Intel® Core i7-6700HQ, CPU@2.60 GHz, 32.0 Gb RAM, GPU NVIDIA Geforce GTX 1070 8 GB DDR5, and 64-bit Operating System on Windows^®^ 10) using different presets of volume rendering. Micro-CT scans at 3 μm resolution were analyzed in a computer with Windows 10 Pro, Intel^®^ Xenon^®^ CPU E5-2640 v4 @ 2.40GHz 2.39GHz (2 processors), NVIDIA Quadro M4000 graphic card, 64-bit Operating System, x64-based processor and 128G RAM.

### SEM and EDS

Extracted eggs were studied, from 2016 through 2019, under an EDAX Quanta 200 for Scanning Electronic Microscopy (SEM) and Electron Dispersive Spectroscopy (EDS) at the Royal Belgian Institute of Natural Sciences, in Brussels, Belgium, where some samples were gold plated using a BAL-TEC SCD 050 Sputter Coater (Royal Belgian Institute of Natural Sciences). The same samples were studied for SEM and EDS Mapping from October 2020 – 2023 at the Iberian Nanotechnology Institute in Braga, Portugal, in low-vacuum mode at 15-30 KV under a Quanta FEG 650 from Thermofisher. EDS mapping data was analyzed using AZtec® software.

### Image processing

Image processing was as follows: photographs with the Canon and Nikon cameras were taken in RAW and JPEG mode and later analyzed using Adobe Photoshop^®^. Whenever possible using RAW files, adjustments were made in the exposure and contrast levels in order to get the best shadows and highlights and saved into TIFF or JPEG format. All photographs were further studied and used in this manuscript in the latter formats after being copied into Microsoft PowerPoint^®^ without file compression and labelled in their lower right corner for identification purposes. Illustrations were also produced in PowerPoint^®^.

### RAMAN spectroscopy

Raman spectroscopy measurements were carried out at International Iberian Nanotechnology Laboratory, Braga, Portugal, at room temperature in a back scattering geometry on an alpha300 R confocal Raman microscope (WITec) using a 785 nm laser with laser P = 30 mW and 532 nm laser with laser P = 5 mW for excitation. The laser beam was focused on the sample using a 50x lens (Zeiss); spectra were collected with a 600 groove/mm grating using 30 acquisitions with 2 s acquisition time.

Raman spectra were acquired with the same conditions as those used for eggshell fragments from dinosaur eggs from Aix-en-Provence and Mèze, as well as chicken eggshell (positive controls). The chicken eggshell spectrum presents peaks at 1087, 713, 282, and 156 cm^-1^, which correspond to the characteristic vibrational modes of calcite, as expected^63^. Despite the lower signal/noise ratio in some spectra, it is clear that all Raman spectra shown (with the exception of the negative control) present the features associated to calcite, revealing similar composition to that of the chicken eggshell. The absence of the calcite mode at ≈ 713 cm^-1^ in the fossilized fragments is not surprising, since this mode is the less intense in the chicken eggshell. Typical Raman spectra without calcite modes were also acquired in each sample in other points.

By fitting each Raman spectrum, important parameters are obtained, such as the peak position and the corresponding full width half maximum (FWHM). Despite the lower intensity ratio between the calcite modes at ≈ 1091 and ≈ 283 cm^-1^ (I_1091_/I_283_) obtained for the fossil fragments, when compared with the chicken eggshell, the peak position and the full width half maximum of these calcites modes are approximately the same, revealing stable calcite.

Using the Spectral Imaging Mode of the alpha300 R, a spectrum at each pixel was acquired by performing a XY scan with a scan range, in the present study usually of 30 μm x 30 μm and 30 x 30 pixels, using a 50x objective with a 0.7 numerical aperture and an integration time of 1s per Raman spectrum. All Raman results/images presented in this paper were calculated from the acquired multi-spectrum file.

Raman images, i.e., Raman Maps can be extracted from the multi-spectrum Raman file by evaluating specific spectral features, by using the WITec Project Plus software tool for advanced microscopic data processing. In this way, Cluster Analysis algorithm can be performed, and the result is a certain number of clusters having similar spectra. Each cluster is represented by an average spectrum and a map showing the spatial distribution of these similar spectra in the analysed area.

By combining all the image clusters, the complete distribution of these clusters in the XY analysed area can be visualized and is given as one colour coded image, named Raman map. With the acquired Raman spectra in the analysed area it is also possible to create intensity maps of desired select Raman modes. In the present study the selected Raman modes for the intensity maps were the calcite Raman modes at ≈ 1091 cm^-1^ and ≈ 283 cm^-1.^

Each figure devoted to the Mapping is composed by the optical image identifying the analized area (marked in color), the corresponding Raman Map and Raman spectra basis (the pixel color in the Raman Map is related with the color of the Raman spectra basis).

## DATA AND MATERIALS AVAILABILITY

NUS_A6_2016, NUS_N7_2016, NUS_M1_2019, NUS_A3_2016 and EHG_A1_2016 are kept at the Ningcheng Administration for National Geopark (NANG), Ning Cheng, EHG_J1_2016 and EHG_J2_2016 at the Sichuan Chongzhou Tianyan Museum, NUS_N1_2016 and NUS_N9_2016 at the Inner Mongolia Natural Museum. The data that supports the findings of this study is available from the corresponding author upon reasonable request.

## ACKNOWLEDGMENTS

Many thanks to the staff at the Chinese Museums involved in this study for help and support. At ALERT, Pedro Sá for illustrations, Aurora Peixoto for manuscript review and the entire staff for support. At the Royal Belgian Institute of Natural Sciences (RBINS), Pascal Godefroit for important discussions, support and critical review of the manuscript, Julien Cillis for help with SEM and EDS, Thierry Leduc and Ulysse Lefèvre for support. At Eldonia, François and Isabelle Escuillié for dinosaur egg samples, support and expertise. At Yang Ming University, Animal Center at Chung Gang University and National Biotechnology Research Park, the staff for support. Alice Wong for support. At INL, Oliver Schraidt for assistance with low vacuum SEM and EDS mapping, helpful comments to the manuscript, and, together with Enrique Carbo and Paulo Ferreira, for discussions and help in coordinating the use of different techniques.

## AUTHOR CONTRIBUTIONS

MJG formulated the nesting hypothesis and designed the research plan, analyzed the fossils, extracted the structures for study from the fossils and analyzed them under macrophotography, stereomicroscopy, SEM, EDS and CT scanning, formulated the parity switch hypothesis of animal reproduction without convergence and wrote the manuscript. QS and RD participated in the search and identification of fossil specimens and reviewed the manuscript. MFC conducted the RAMAN experiments in collaboration with MJG under PA supervision. JW, XZ, BG, FM and XW jointly supervised the work performed in the different museums in China and reviewed the manuscript. Y-HC and T-CY participated in the high resolution micro-CT scanning studies and reviewed the manuscript.

## SUPPLEMENTARY MATERIAL

**Figure S1.**
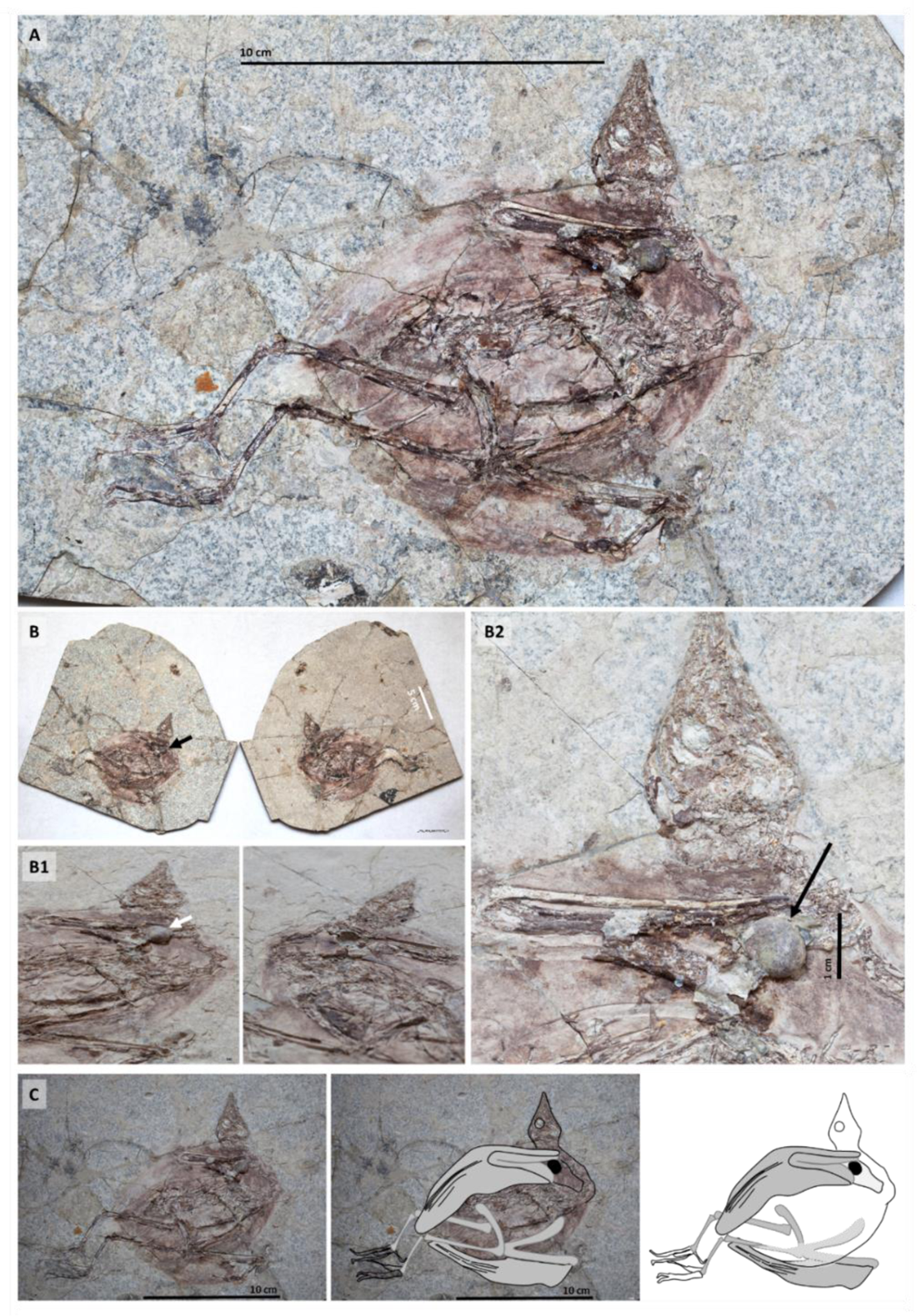
*I. huchzermeyeri* specimen NUS_A6_20216 and its surface exposed NUS_A6_20216 E1 egg. **A:** General view of the fossilized skeleton of NUS_A6_2016. **B:** The two slabs of NUS_A6_2016: main slab (left) and counterslab (right). **B1**: Region where the NUS_A6_2016 E1 egg is located in the main slab and its counterpart in the counterslab. **B2:** Zoom in on the NUS_A6_2016 E1 egg located against the right wing of the NUS_A6_2016 specimen**. C:** Interpretation of *I. huchzermeyeri* NUS_A6_2016 fossil in accordance with a nesting hypothesis. The black globular structure against the right wing is egg NUS_A6_2016 E1.

**Figure S2.**
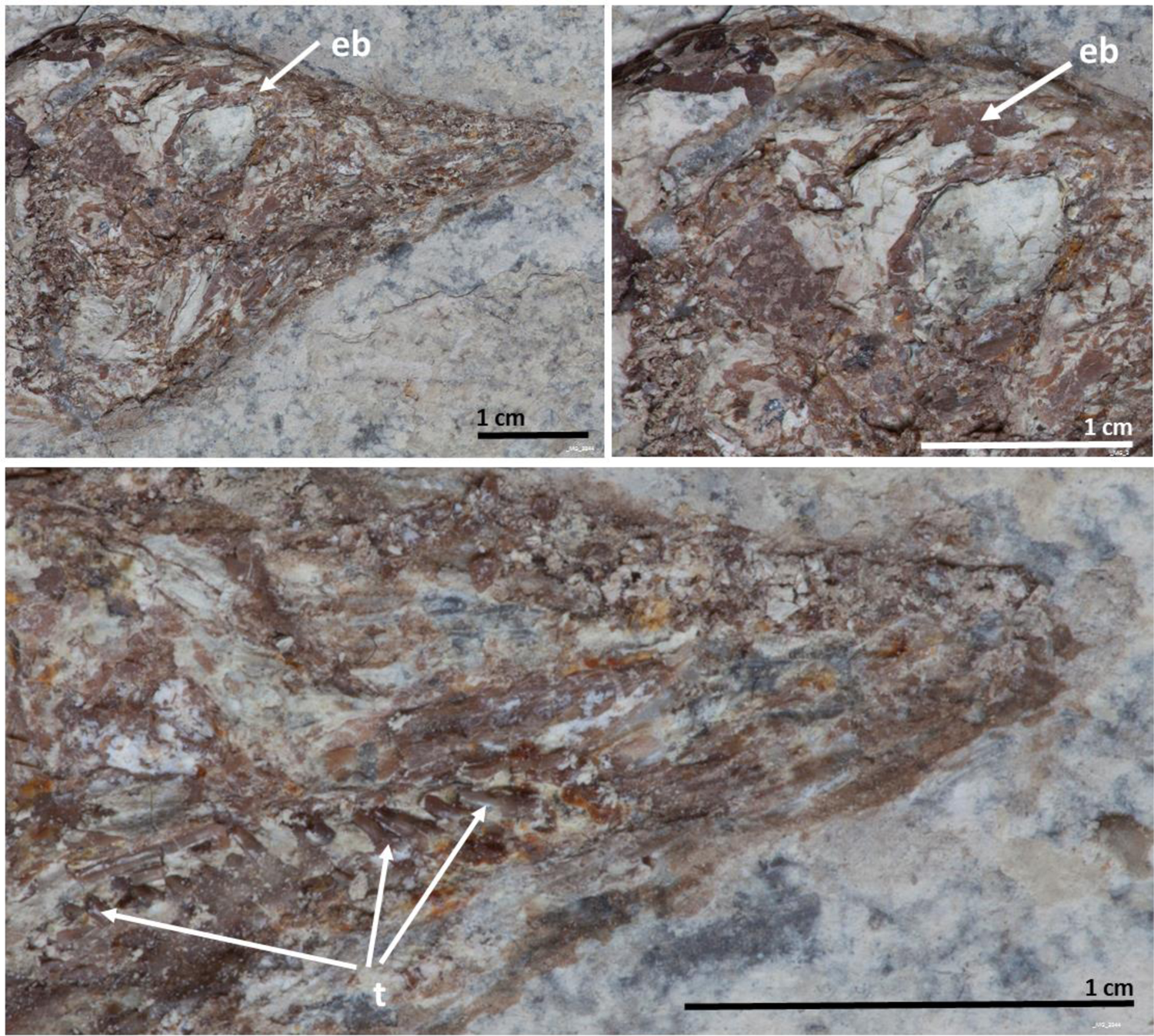
Aspects of the head of the adult skeleton of NUS_A6_2016 (main slab). eb – ethmoid bone; t – teeth.

**Figure S3.**
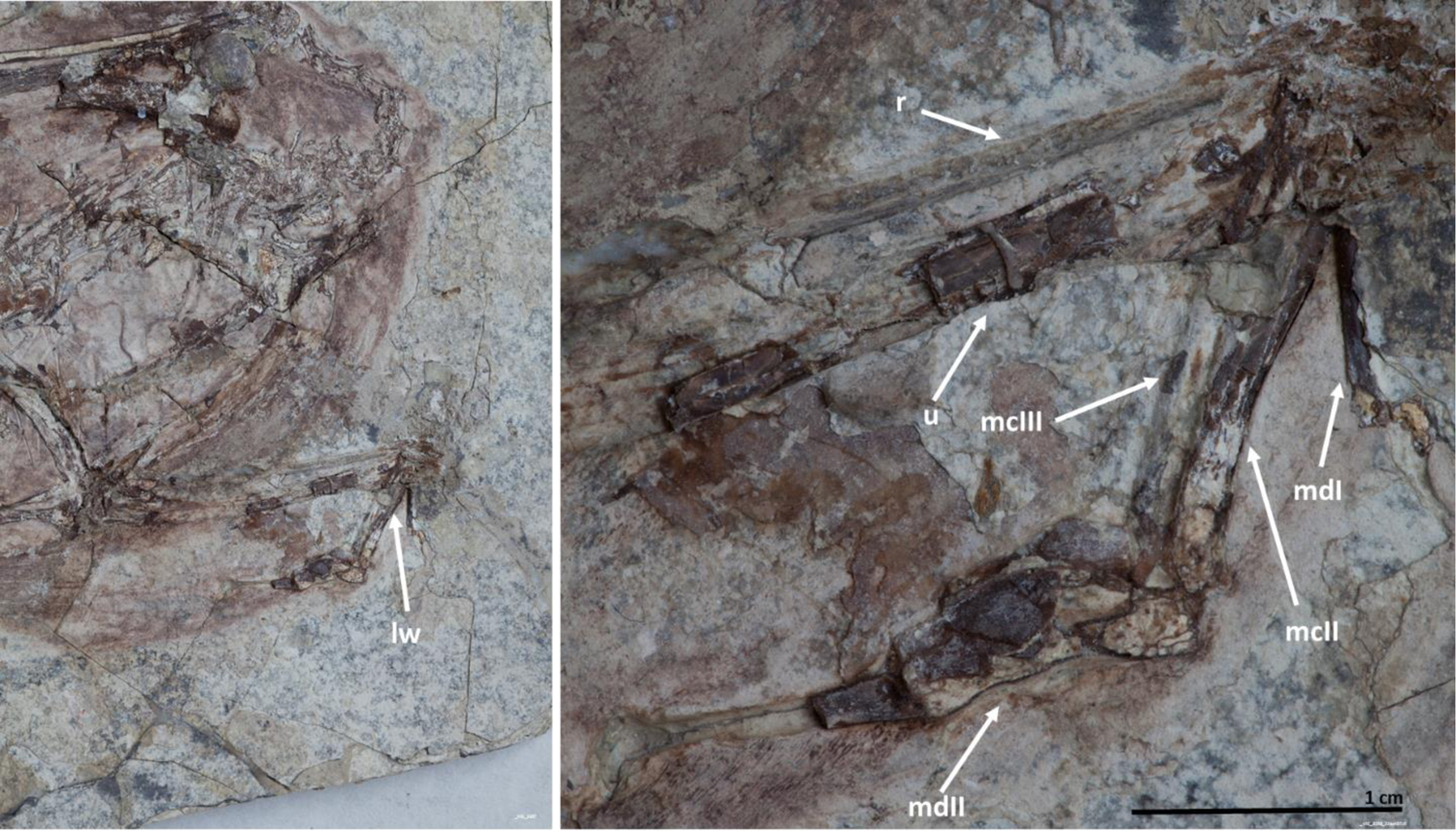
Details of the left wing (lw) skeleton of NUS_A6_2016 (main slab). mcII – major metacarpal; mcIII – minor metacarpal; mdI – manual digit I; mdII - manual digit II; r – radius; u – ulna.

**Figure S4.**
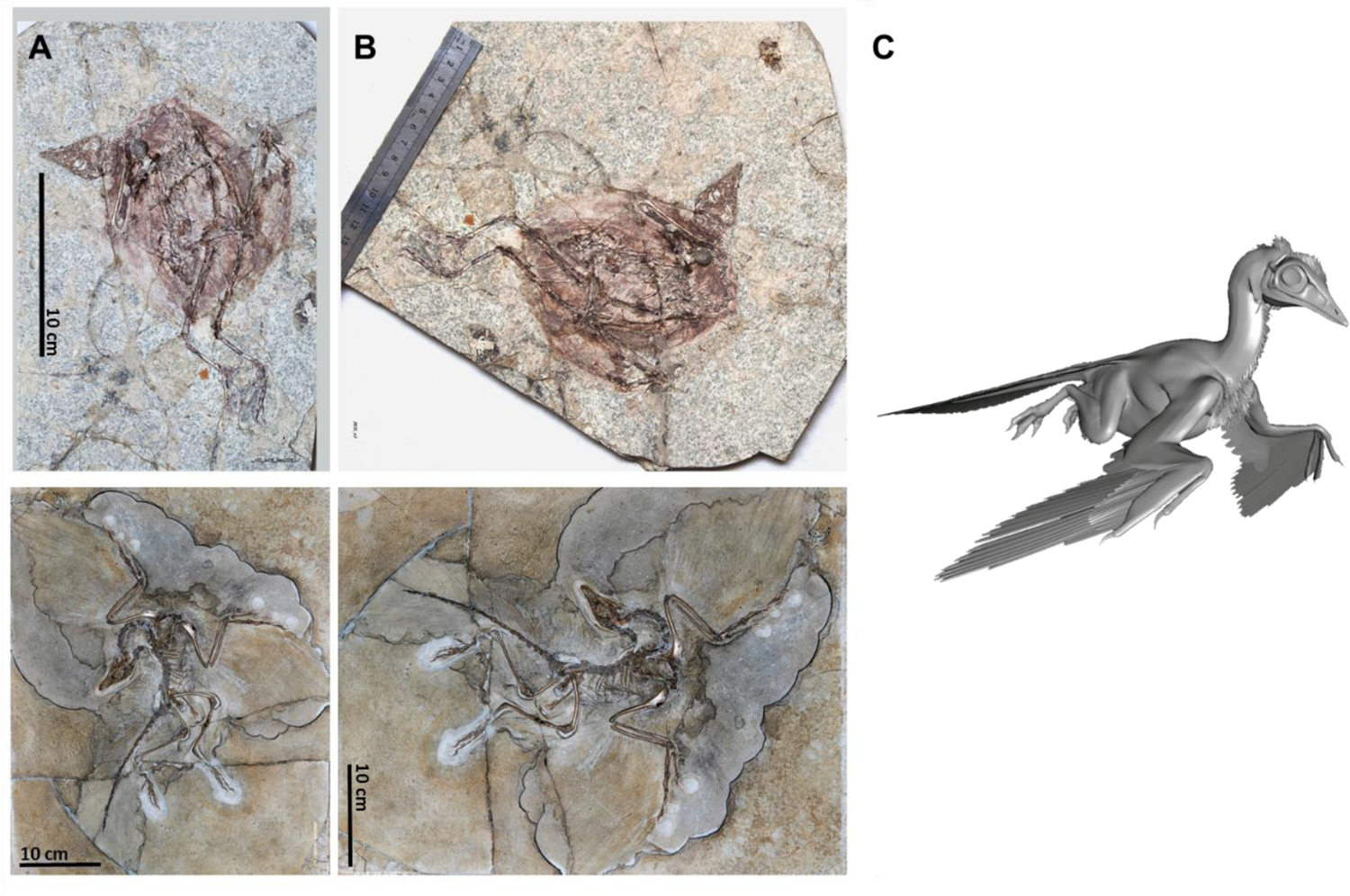
Specimen NUS_A6_2016 of ornithuromorph *I. huchzermeyeri* fossilized in a pose similar to that of the Berlin specimen of *Archaeopteryx*. NUS_A6_2016 (**A** top and **B** top) is shown in similar orientations to those of the Berlin specimen of *Archaeopteryx* in its usual display position (**A**, bottom) and in accordance with the nesting hypothesis (**B**, bottom). A software 3D model of a nesting posture that could have resulted in these fossilized poses is shown in **C**.

**Figure S5.**
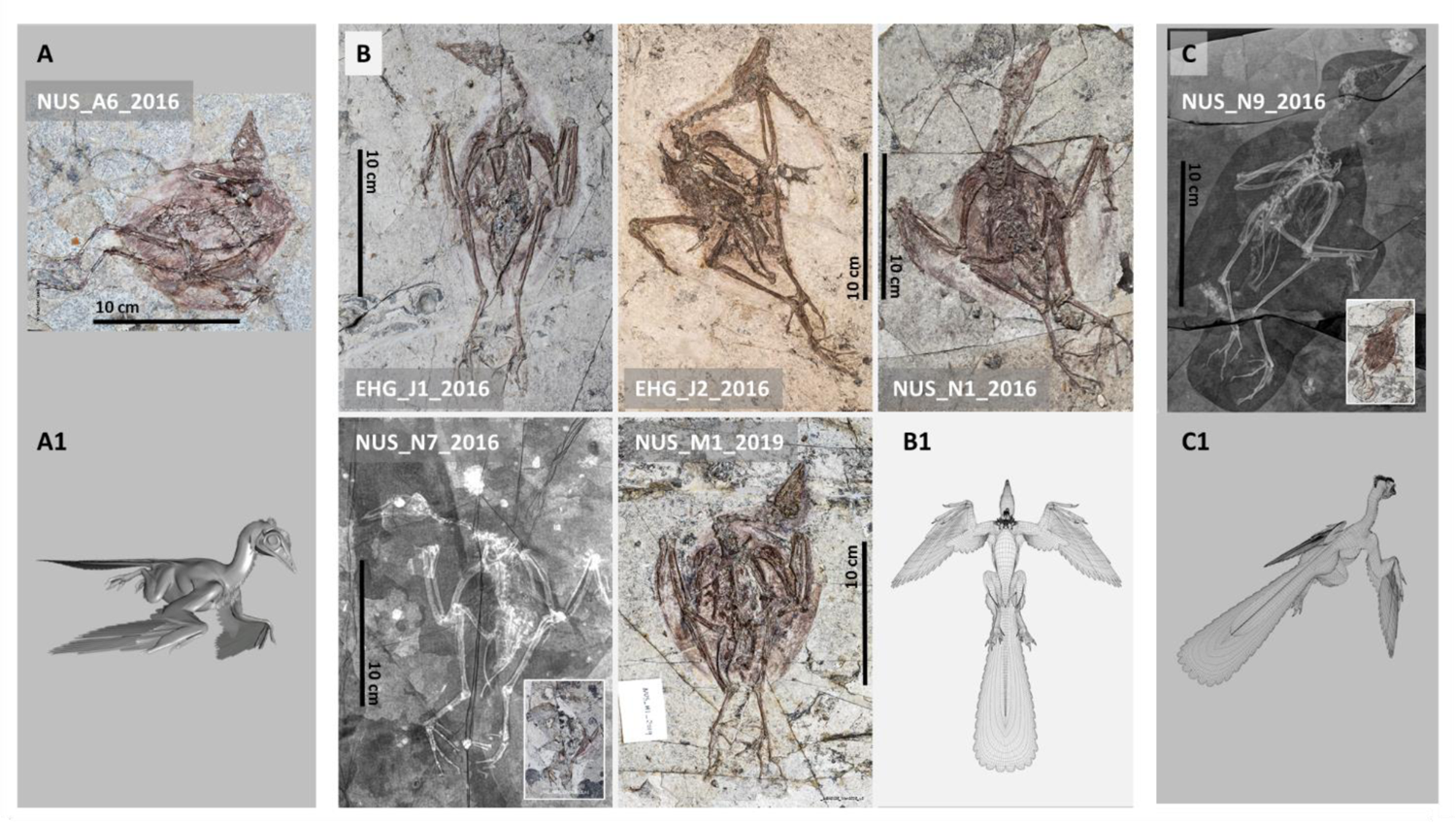
Seven specimens of ornithuromorph *I. huchzermeyeri* as different views of the same fossilized pose. NUS_A6_2016 (**A**) corresponds to a ventro-lateral view (illustrated in **A1**), the five specimens in the center (**B**) correspond to ventral views (illustrated in **B1**) and the specimen on the right to a dorso-lateral view (illustrated in **C**) of the same pose nesting posture.

**Figure S6.**
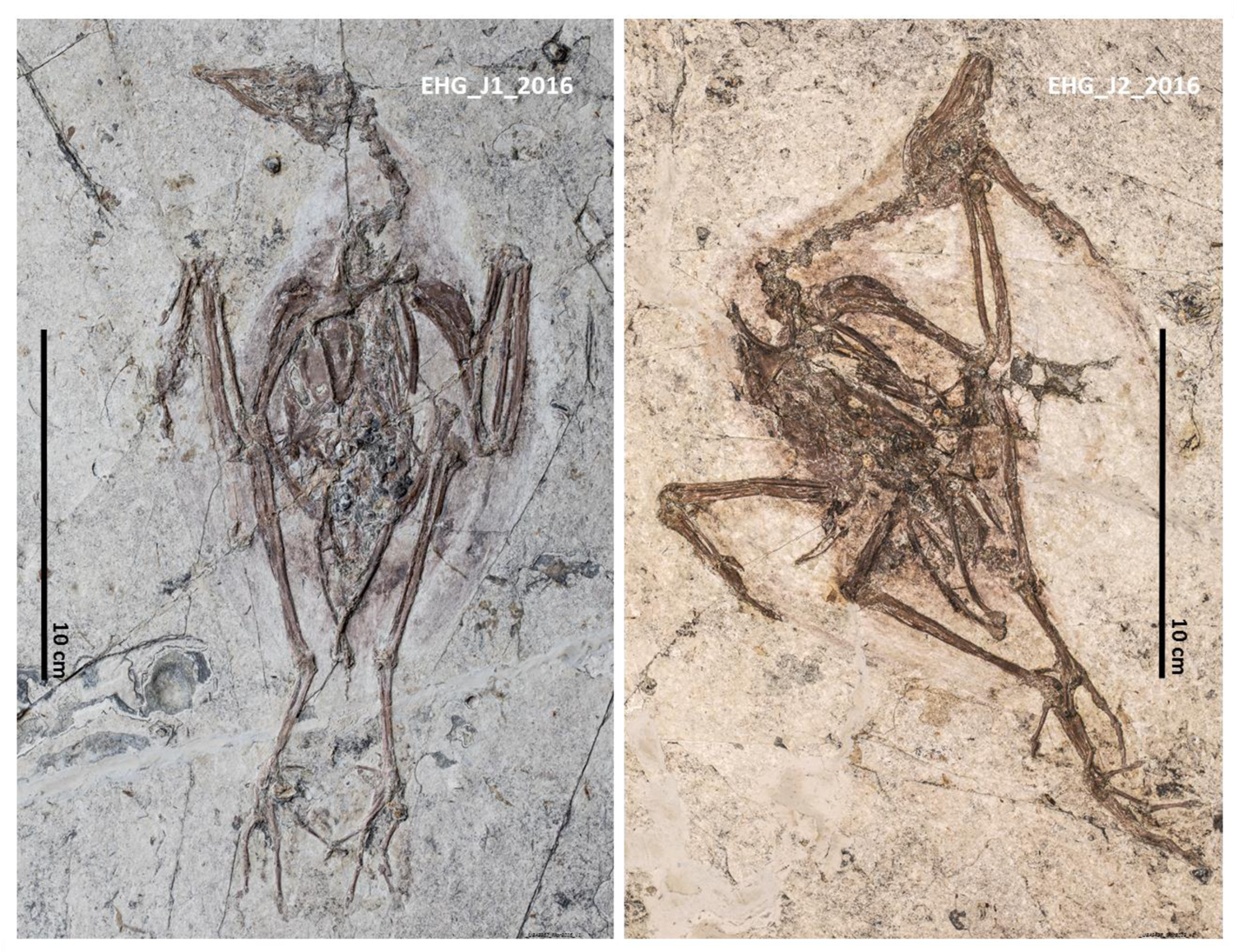
EHG_J1_2016 (left) and EHG_J2_2016 (right) as ventral views of birds fossilized in a nesting posture.

**Figure S7.**
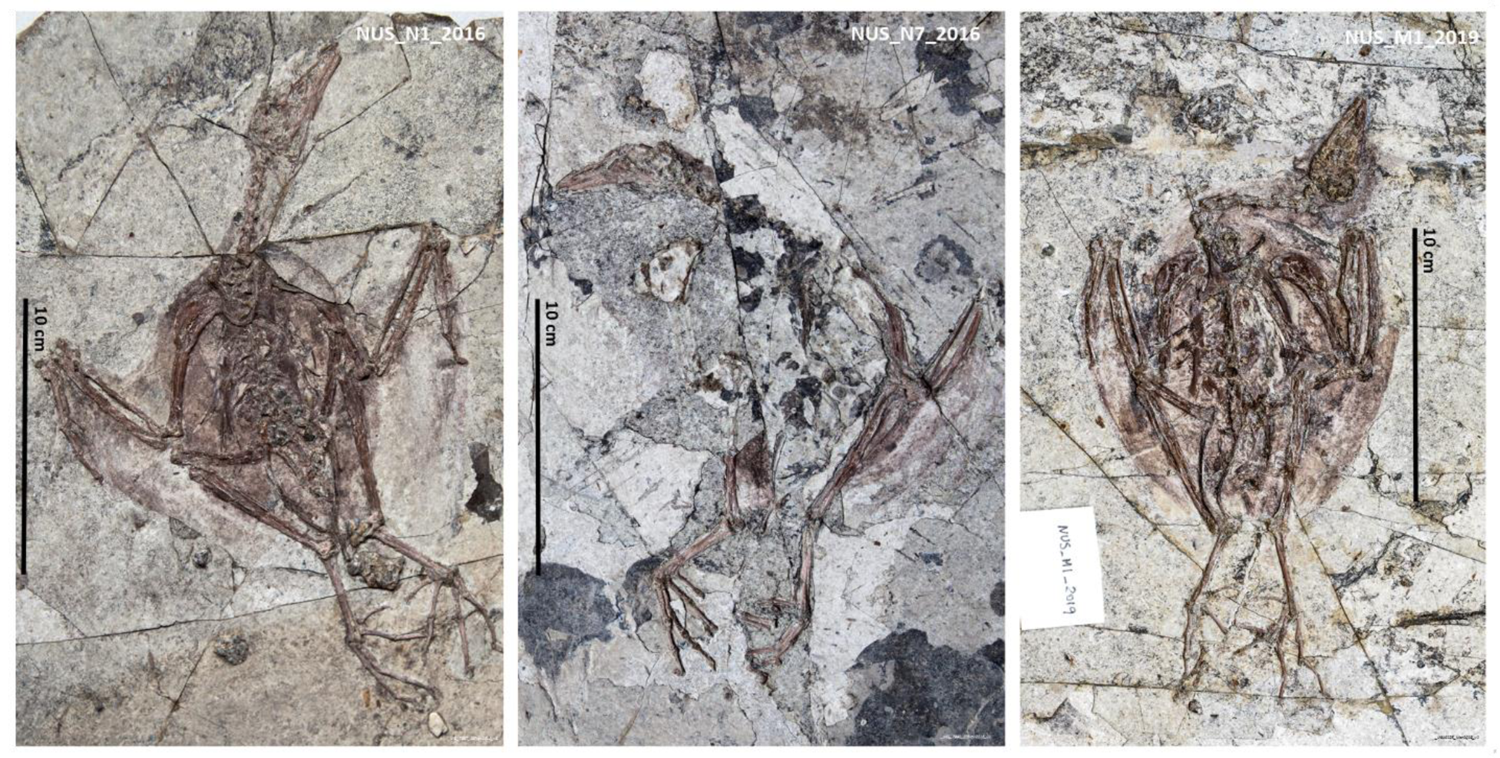
NUS_N1_2016 (left), NUS_N7_2016 (center) and NUS_M1_2019 (right) as ventral views of birds fossilized in a nesting posture.

**Figure S8.**
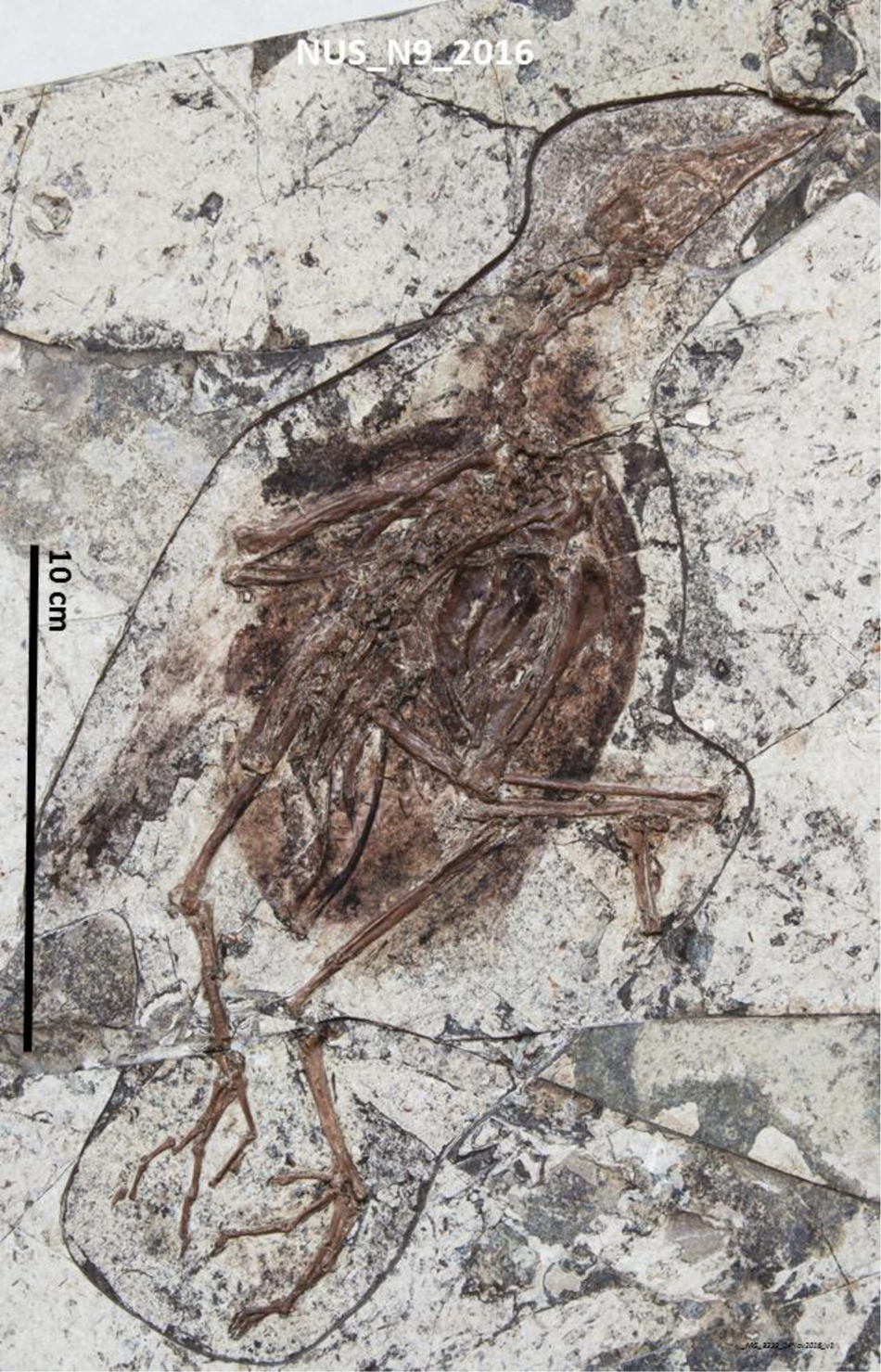
NUS_N9_2016 as a dorsolateral view of a bird fossilized in nesting posture.

**Figure S9.**
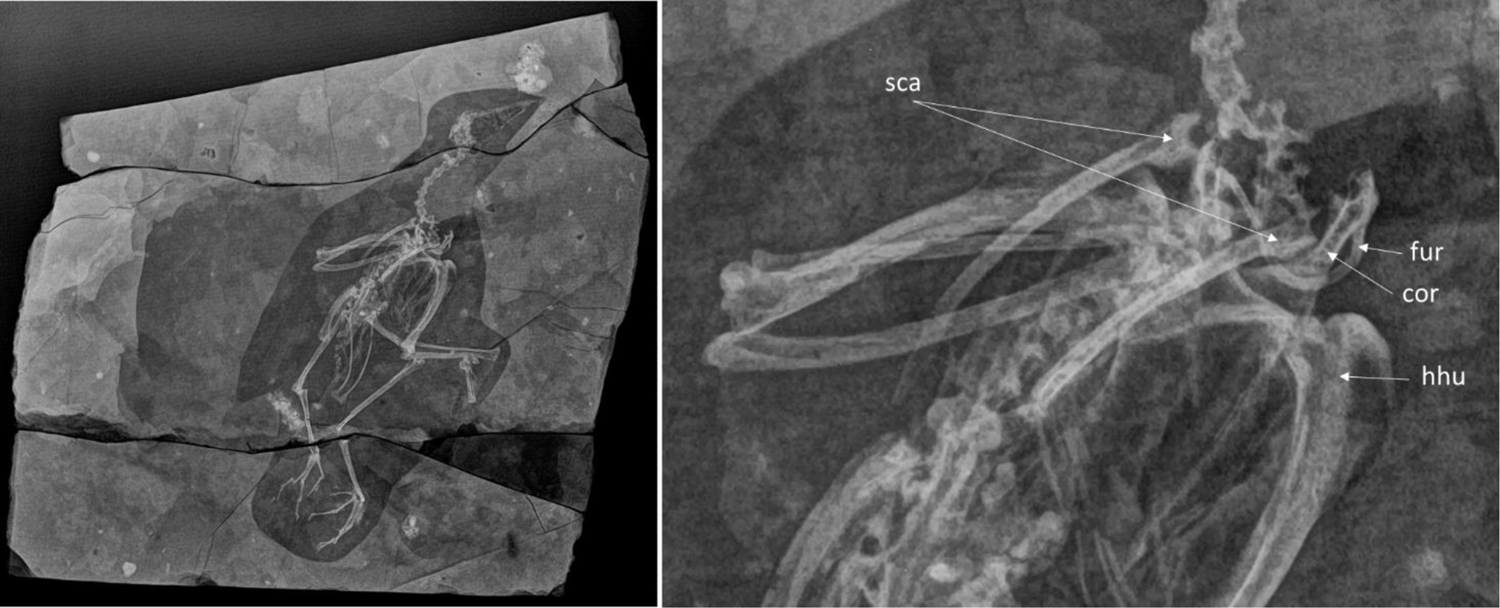
X-ray images of NUS_N9_2016 to put into evidence its interpretation as a dorsolateral view of the nesting position and details of its shoulder anatomy. cor-coracoid; fur – furcula; hhu – head of the humerus; sca – scapulae.

**Figure S10.**
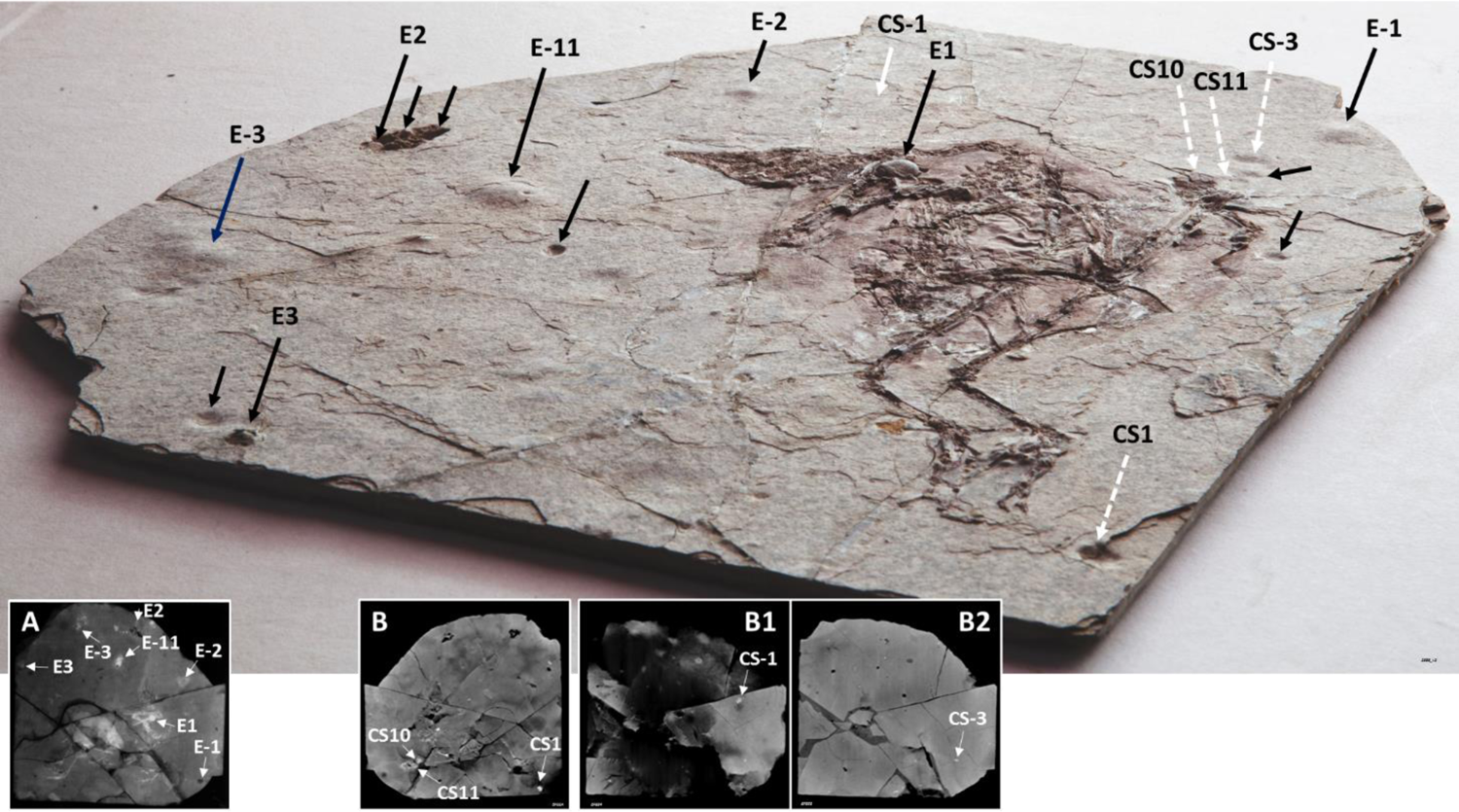
The main slab of *I. huchzermeyeri* specimen NUS_A6_2016 contains other globular structures with similar characteristics to NUS_A6_2016 E1. Oblique view of the main slab with indication of the location of convexities (long black full arrows), concavities that match convexities in the counterslab (white dashed arrows; see also Figures S11 and S12), in addition to the projection onto the main slab of the location of CS-1 identified in the counterslab (full white arrow; see also Figures S11 and S12), and their corresponding images in coronal sections of CT scans of the main slab (**A**) and counterslab (**B – B2**). Additional concavities are indicated by short black arrows.

**Figure S11.**
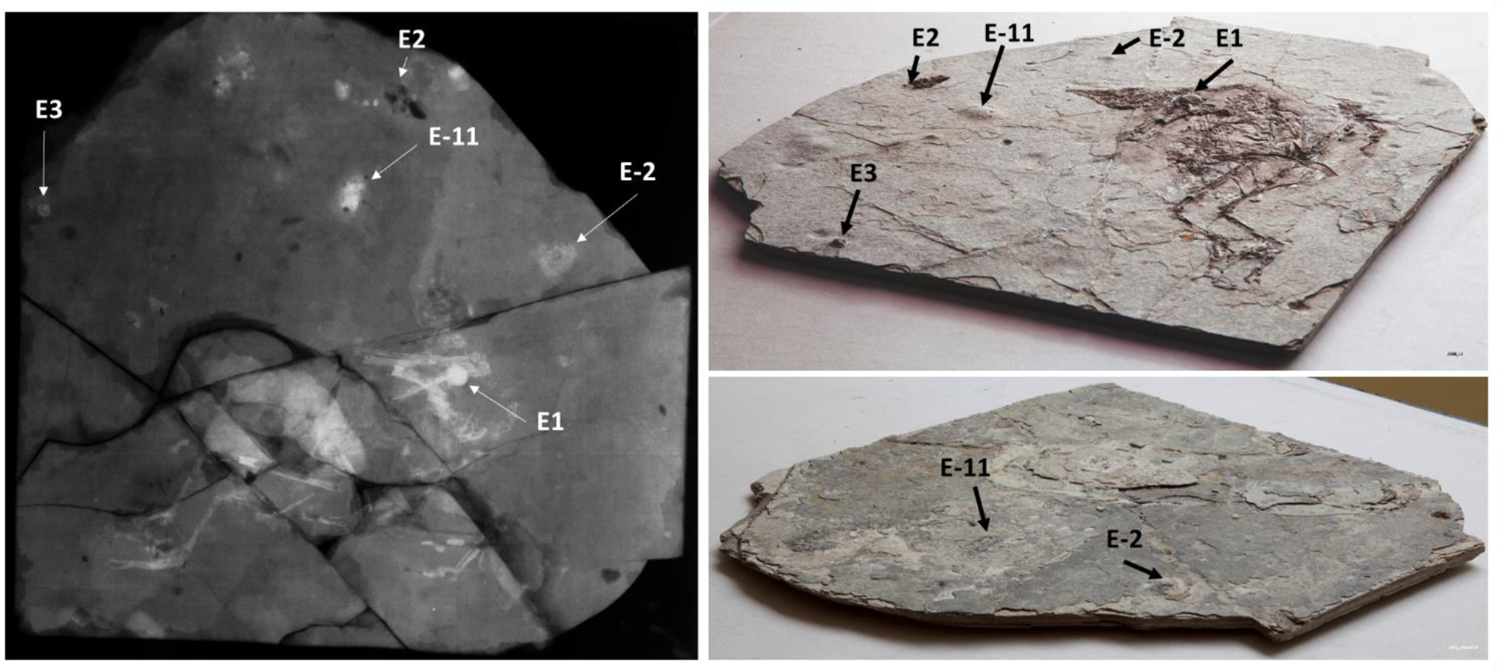
Location of five studied structures from the main slab of the NUS_A6_2016 fossil. These are shown under a coronal CT scan view at 0.5 mm resolution (left) and oblique views of the front and back surfaces of the same slab captured under macrophotography (top and bottom right, respectively). Convexities (full arrows) and concavities (dashed arrows) of similar size are also shown.

**Figure S12.**
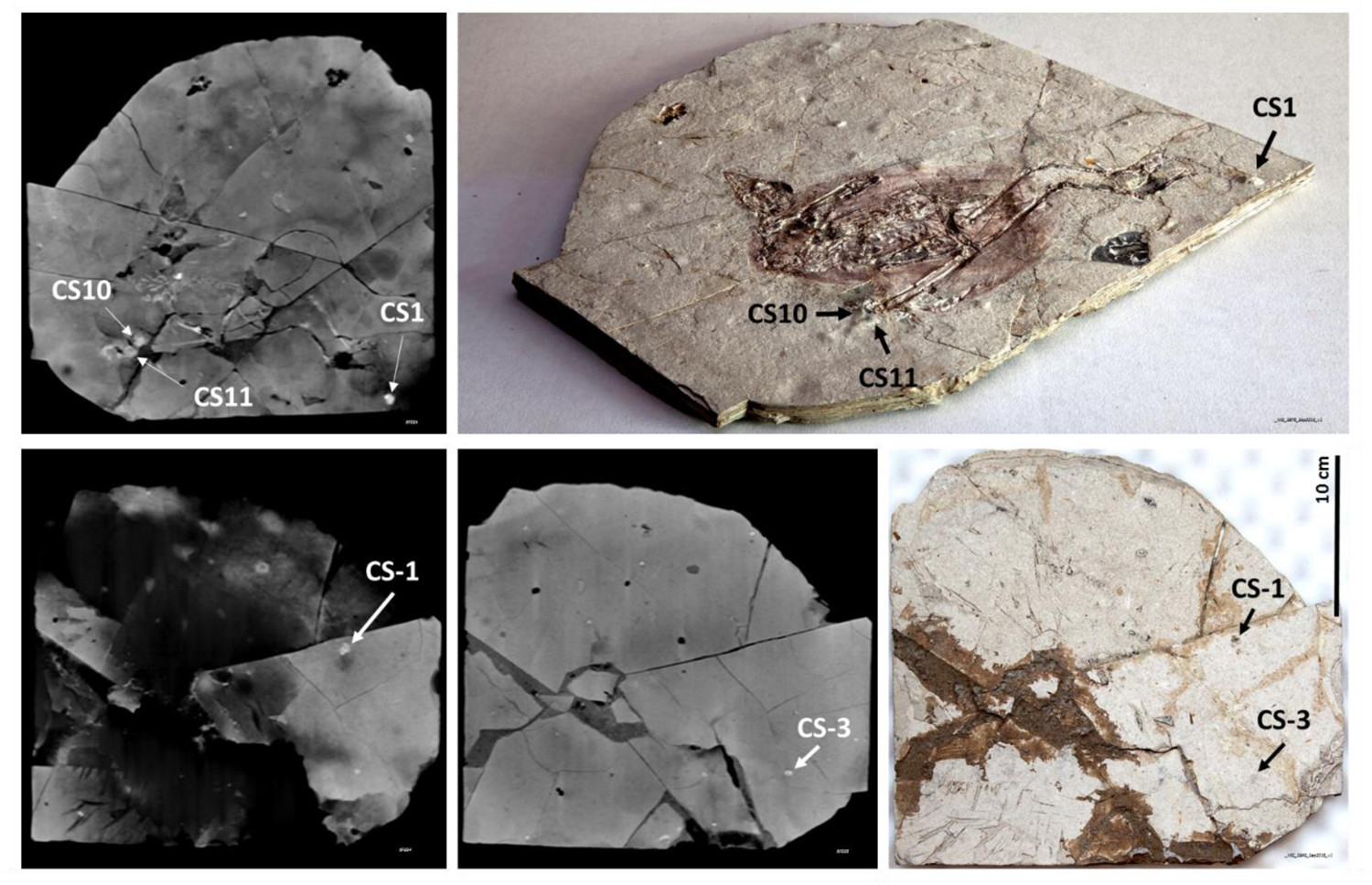
Location of five studied structures from the counterslab of the NUS_A6_2016 fossil. These are shown under coronal CT scan views at 0.5 mm resolution (top left in front to back orientation; bottom left and center in back to front orientation), an oblique view of the front surface of the same slab (top right), and a top view of the back surface of the same slab (bottom right).

**Figure S13.**
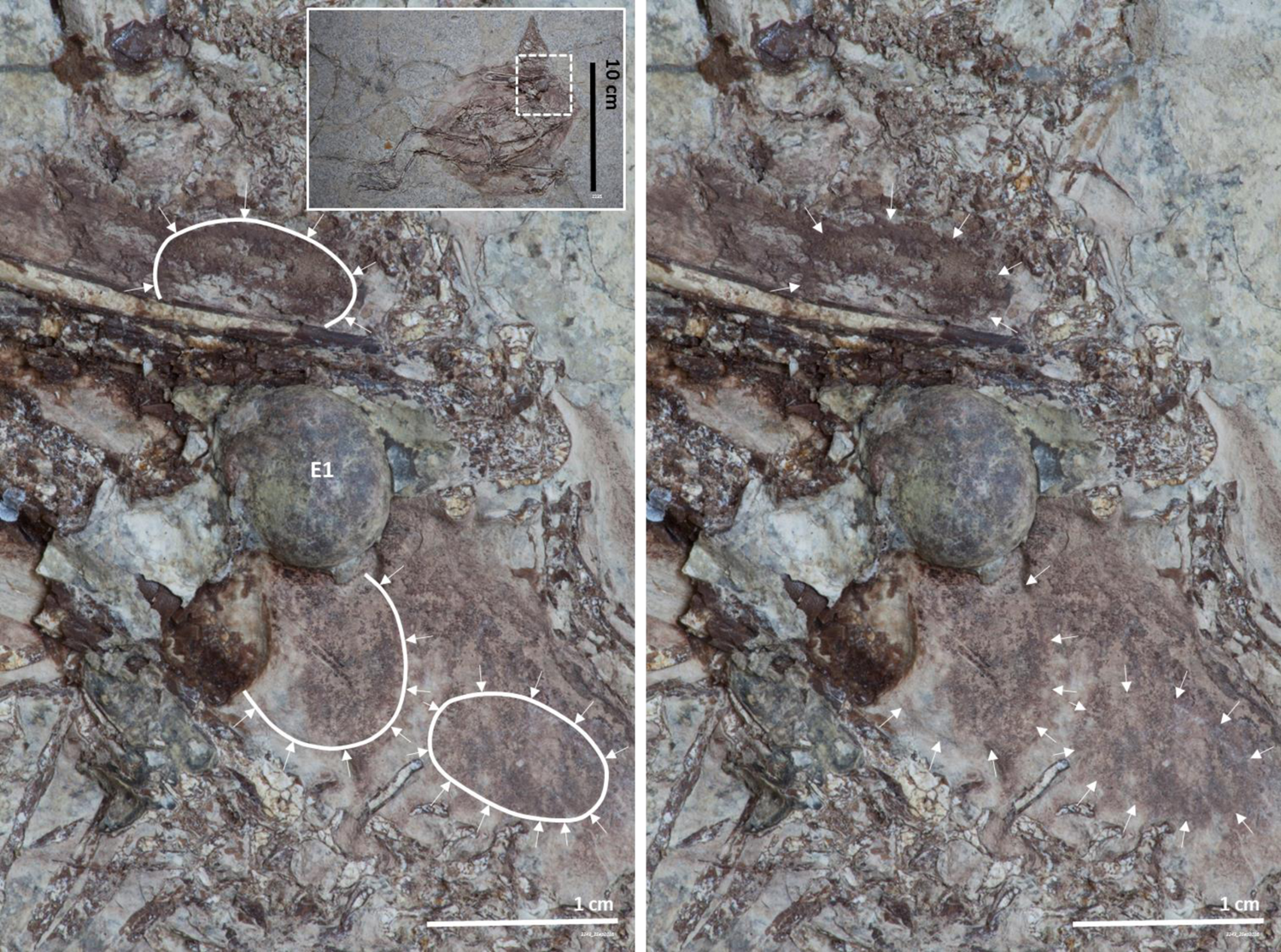
Impressions in the vicinity of the NUS_A6_2016 E1 egg of structures with 2D outlines that would have had a 3D structure that is similar to that of NUS_A6_2016 E1. These may have fallen off when the two slabs of this museum fossil were separated.

**Figure S14.**
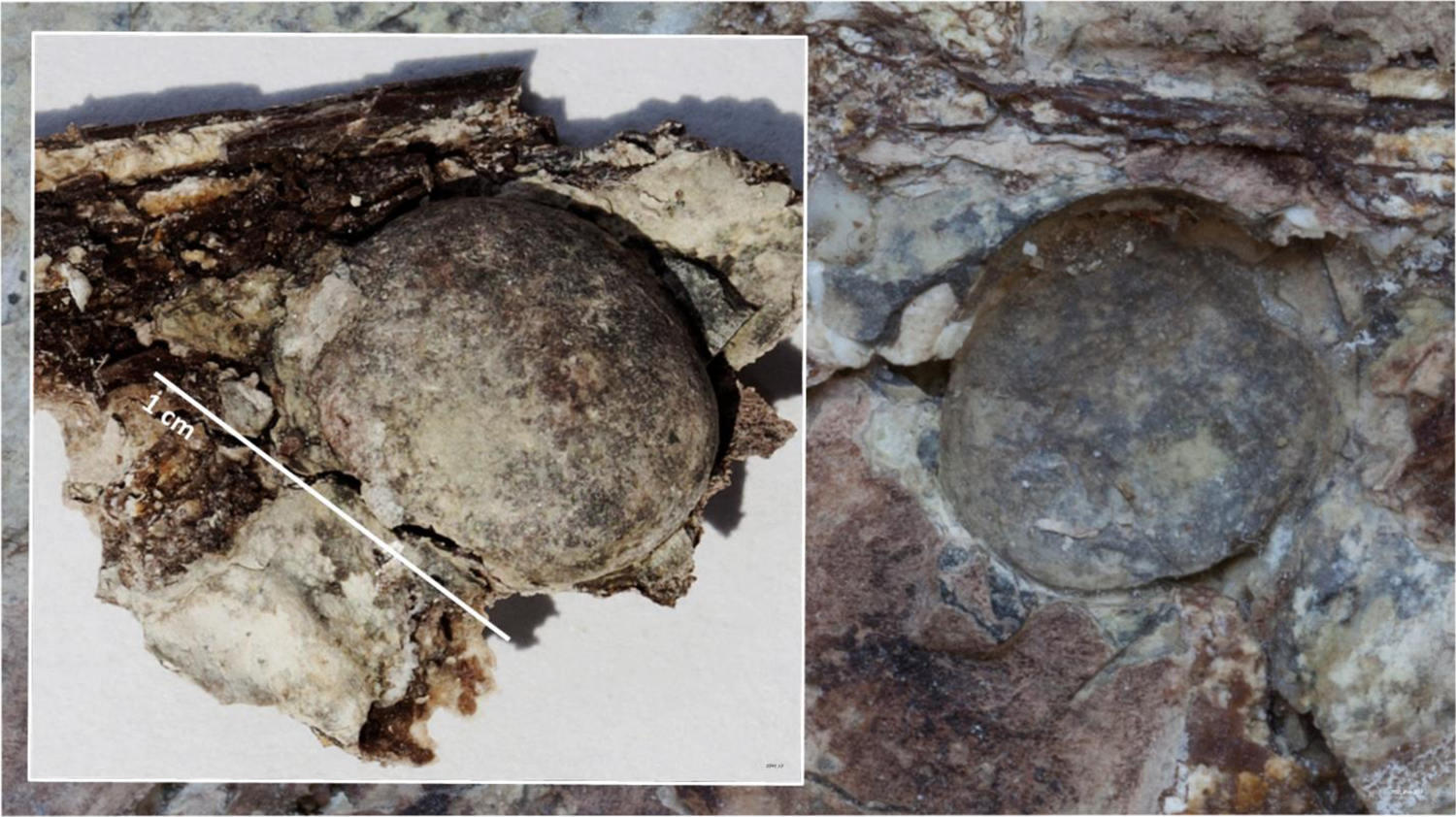
Macrophotography of NUS_A6_2016 E1 after it was extracted from the fossil’s main slab (left) and its 3D impression on the counterslab (right).

**Figure S15.**
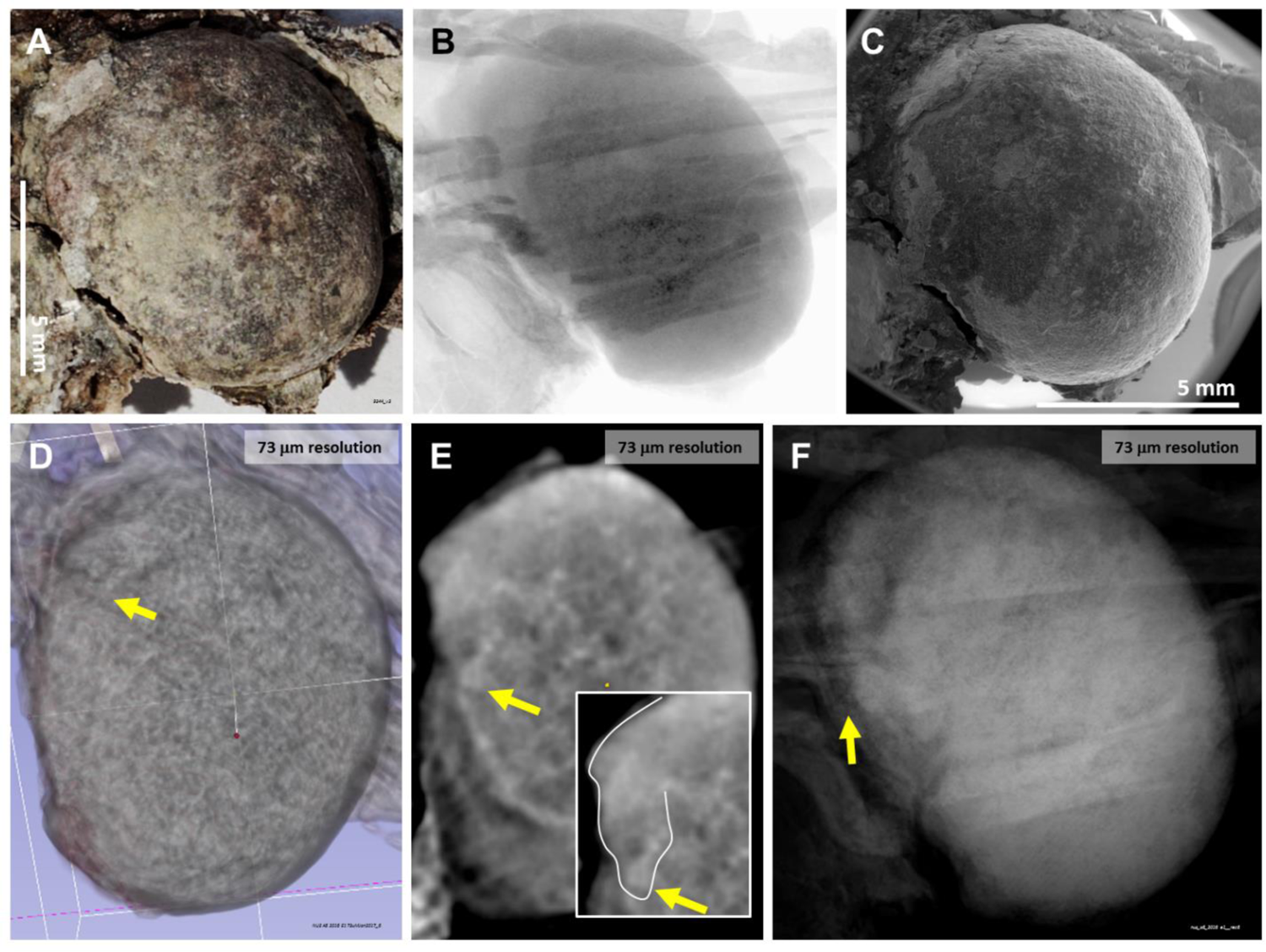
NUS_A6_2016 E1 egg. This egg is shown here under macrophotography (**A**), X-ray (**B**), SEM (**C**) and CT-scanning at 73 μm resolution (**D-F**). The yellow arrow points at the area (outlined in E) investigated further at higher resolutions for the possibility of representing the outer morphology of a fetal head (see also Figures 2, S29 and S30).

**Figure S16.**
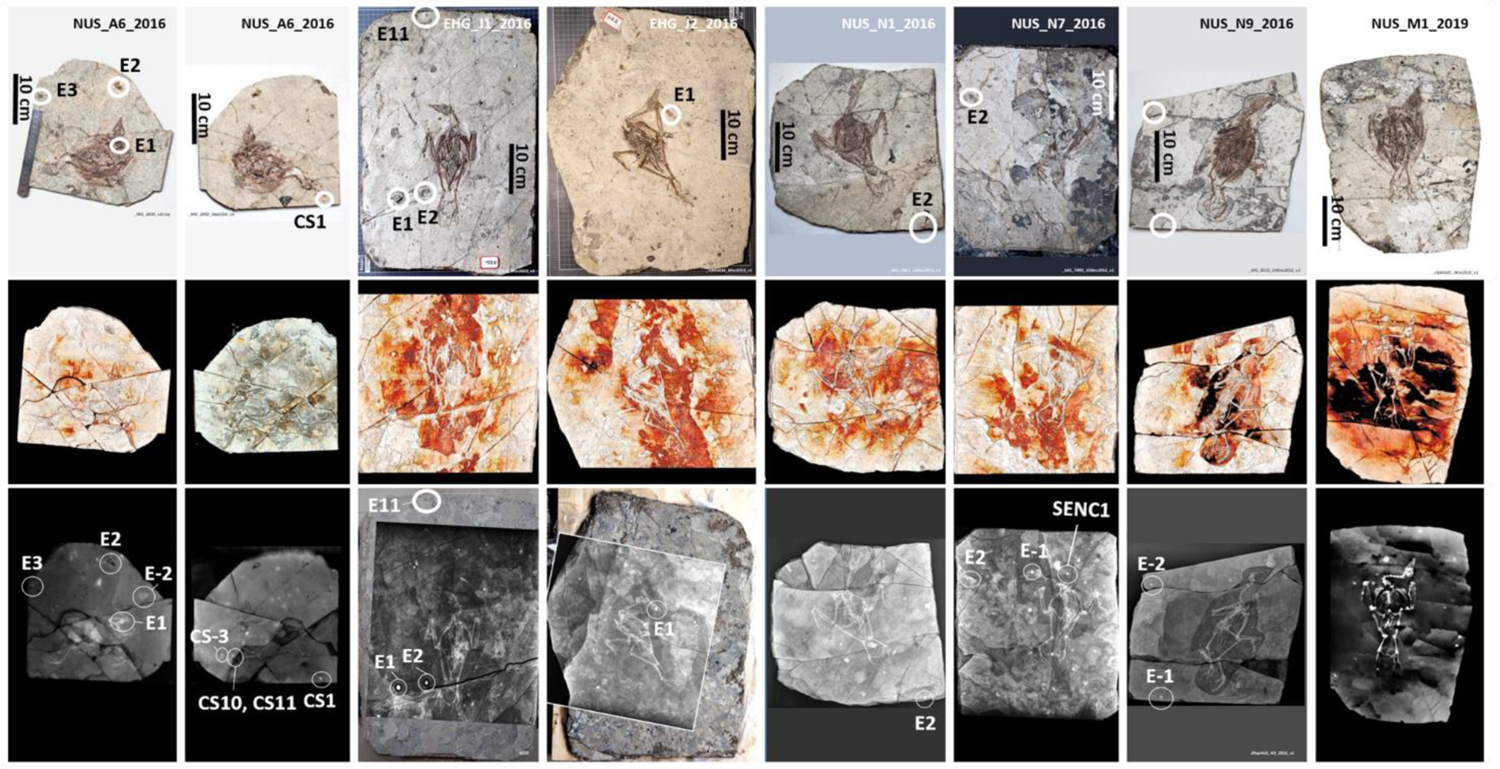
Seven specimens of ornithuromorph *I. huchzermeyeri* have a heterogeneous fossilized environment and 1 cm globular structures around their skeletons. Specimen NUS_A6_2016 is represented by both the main slab and the counterslab (first two columns from the left), whereas all other specimens are represented by the main slab only. Under 3D software analysis of CT scan data, each fossil shows heterogeneous radiodensity in the area that immediately surrounds the fossilized skeleton (middle), suggesting *in loco* body decomposition and fossilization. Globular radiodensities approximately 1 cm in size are shown in association with each skeleton (bottom).

**Figure S17.**
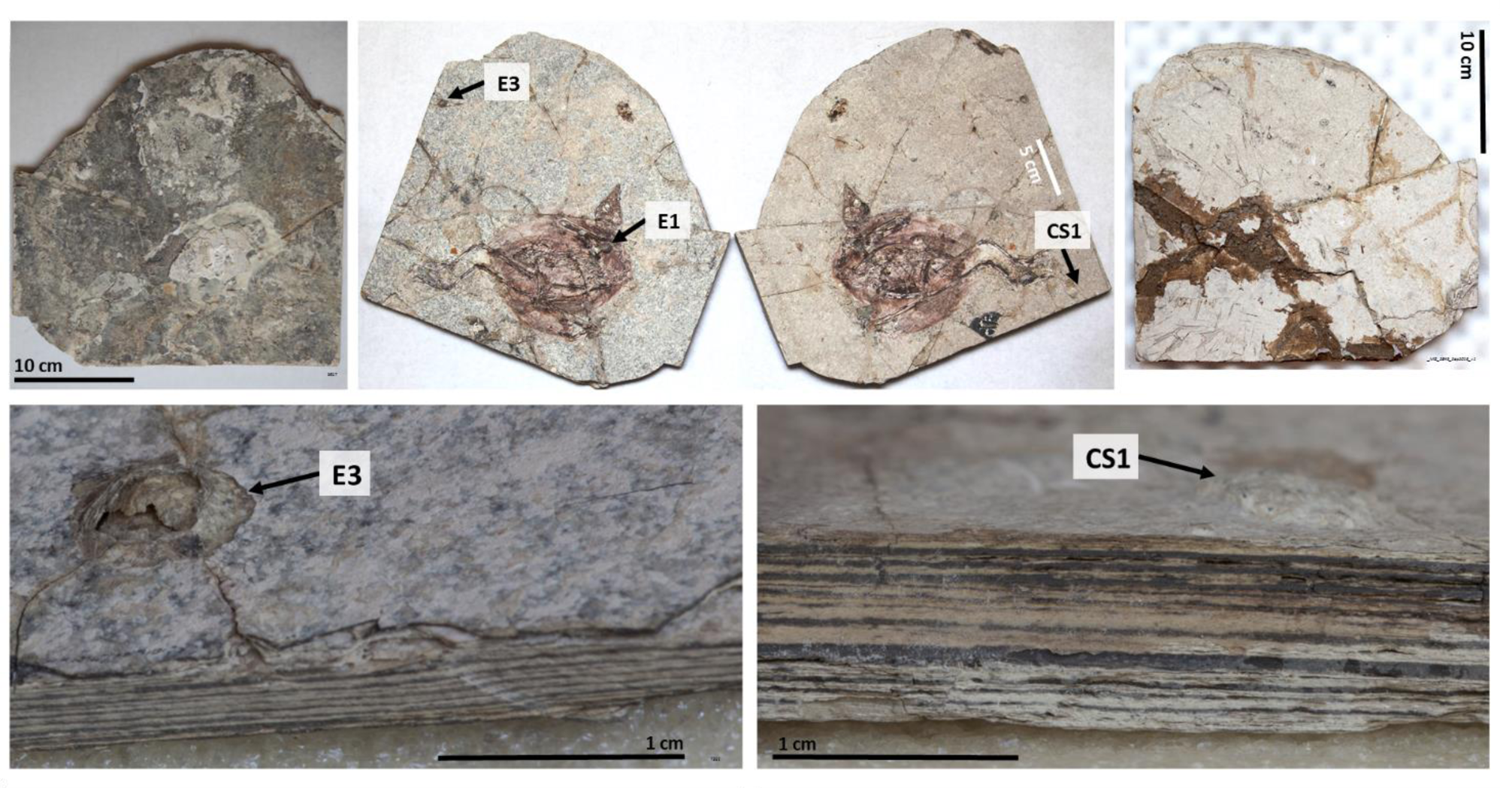
NUS_A6_2016 E3 and NUS_A6_2016 CS1 are two eggs associated with the NUS_A6_2016 specimen fossil. Like NUS_A6_2016 E1, NUS_A6_2016 E3 and NUS_A6_2016 CS1 are examples of radiodense structures associated with the NUS_A6_2016 specimen that also measure approximately 1 cm but are not as surface exposed and 3D preserved as the former. The back and front surfaces of the main slab are shown above on the left. The front and back surfaces of the counterslab are shown above on the right. Close-up lateral views of NUS_A6_2016 E3 and NUS_A6_2016 CS1 are shown at the bottom.

**Figure S18.**
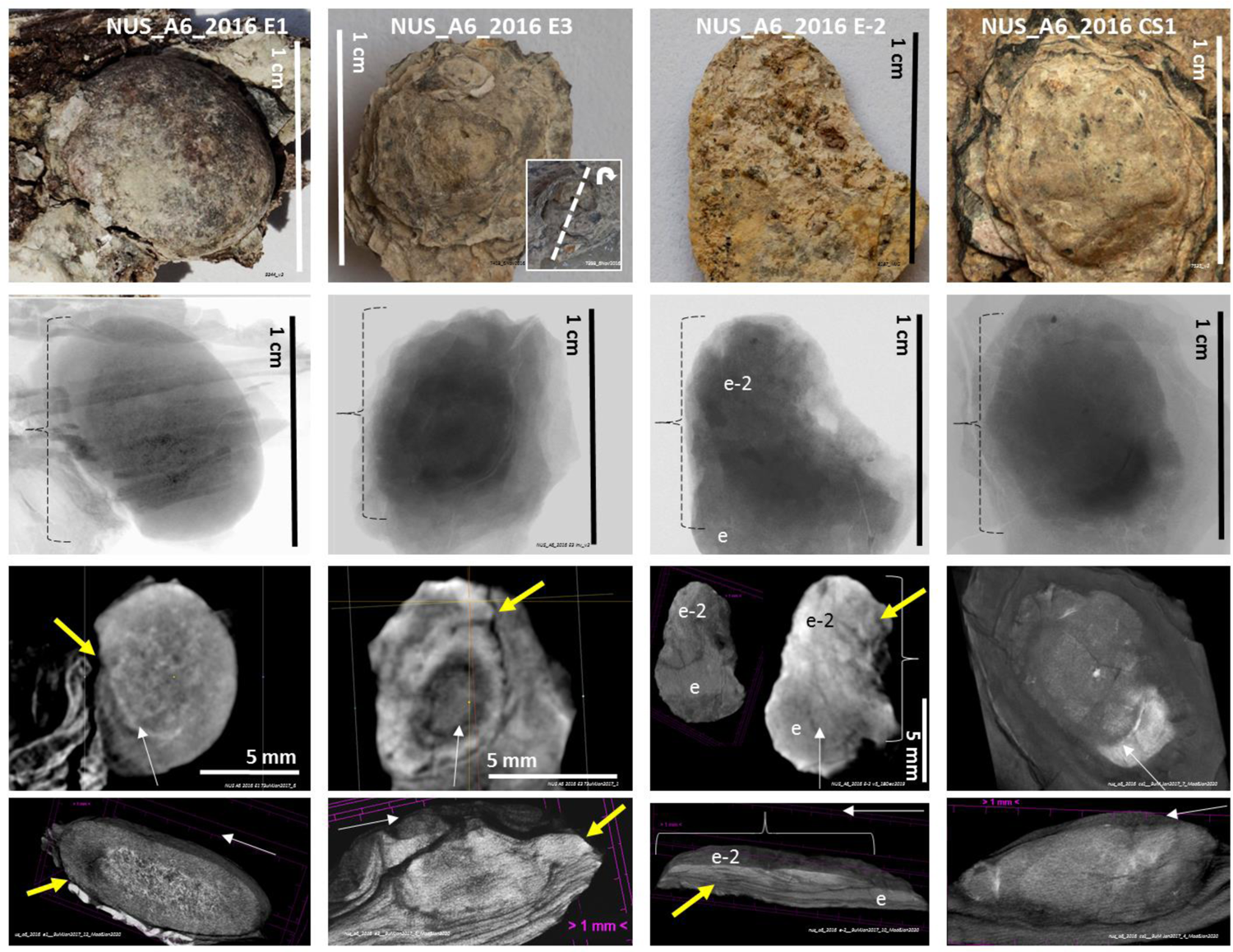
Four NUS_A6_2016 eggs. Four eggs extracted from the NUS_A6_2016 fossil under macrophotography (top), high resolution X-ray (captured in the chamber of CT scanners) and CT scanning in coronal and sagittal views at 73 μm (for NUS_A6_2016 E1 and NUS_A6_2016 E3) and 9 μm resolution (for NUS_A6_2016 E-2 and NUS_A6_2016 CS1), except in the case of NUS_A6_2016 E3 for a which a virtual sagittal cut on a 3D view is shown instead of a 2D sagittal view. Where applicable, yellow arrows point at the presumed head location of the fossilized fetus with surface morphology preservation. Two eggs (e-2 and e) may be overlapping in NUS_A6_2016 E-2. White arrows are used for cross-orientation of different CT scan views of the same sample.

**Figure S19.**
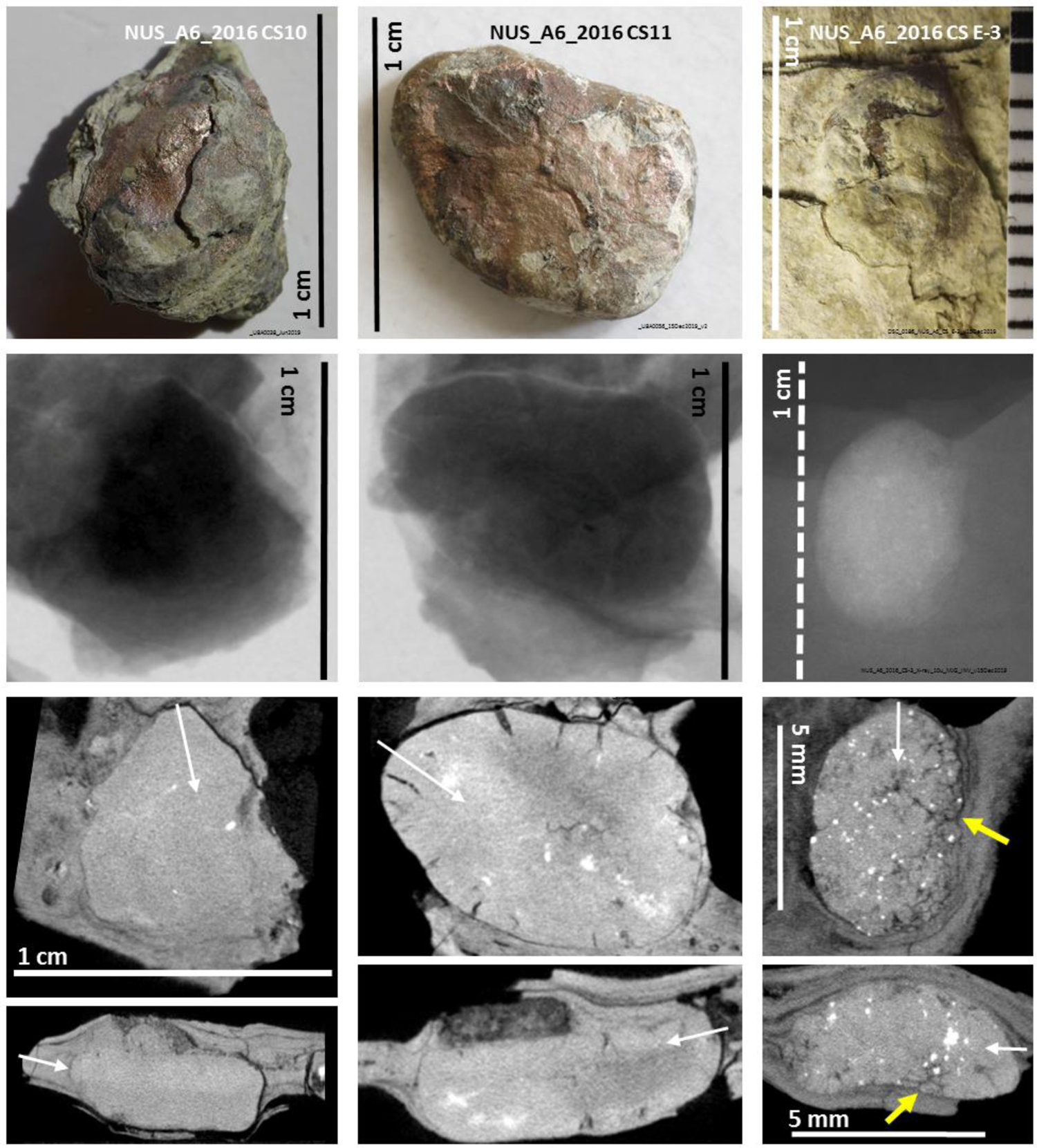
Three additional NUS_A6_2016 eggs. Three other eggs extracted from the NUS_A6_2016 fossil under macrophotography (top), high resolution X-ray imaging (captured in the chamber of CT scanners; at 73 μm for NUS_A6_2016 CS10 and CS11 and at 10 μm for NUS_A6_2016 CS E-3) and coronal and sagittal views of micro-CT scans. A yellow arrow points at the presumed head location of the fossilized fetus with surface morphology preservation of NUS_A6_2016 CS E-3 (see also Figure 2). White arrows are used for cross-orientation of different CT scan views of the same sample.

**Figure S20.**
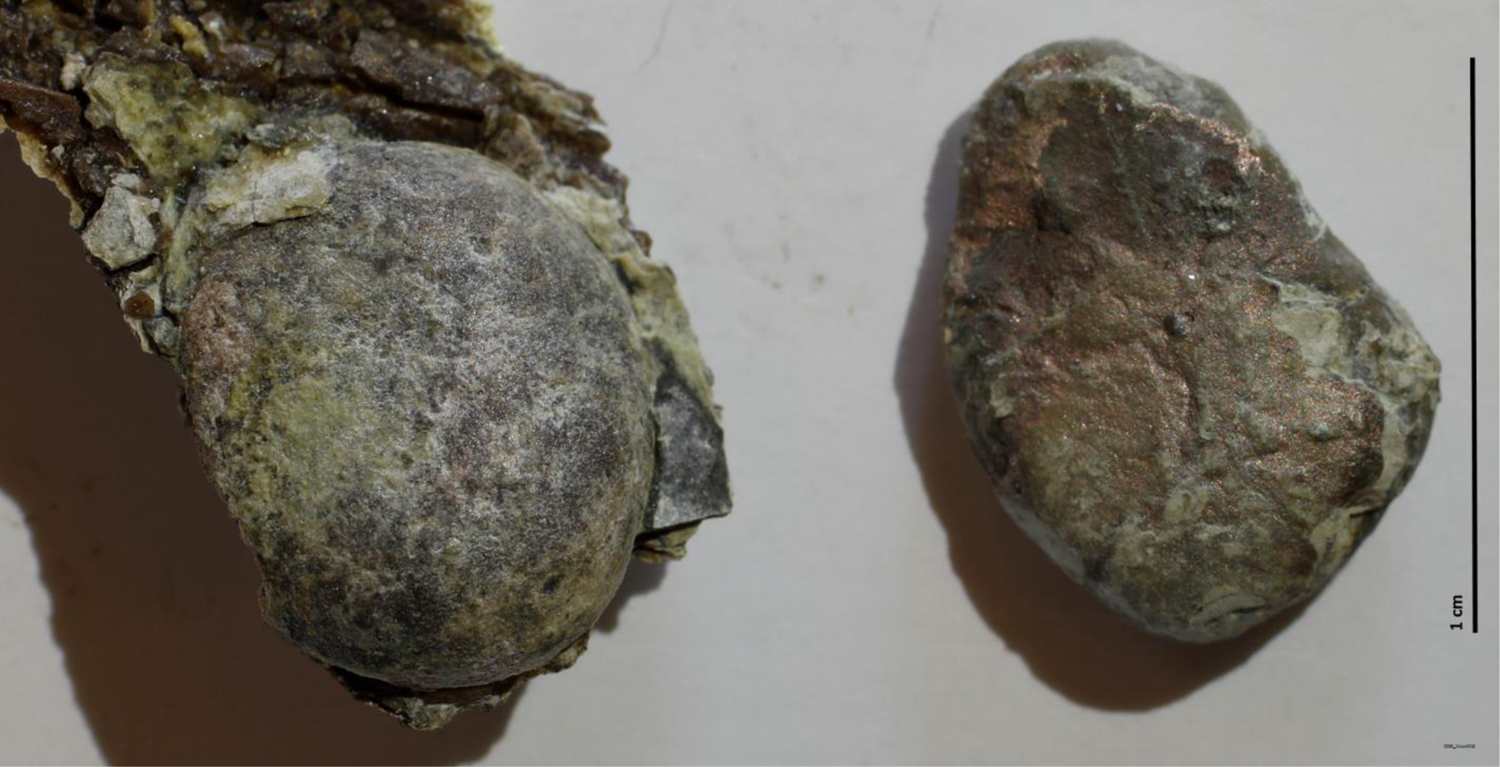
Two eggs extracted from the NUS_A6_2016 fossil. NUS_A6_2016 E1 (left) was extracted from the main slab and NUS_A6_2016 CS11 (right) was derived from the counterslab.

**Figure S21.**
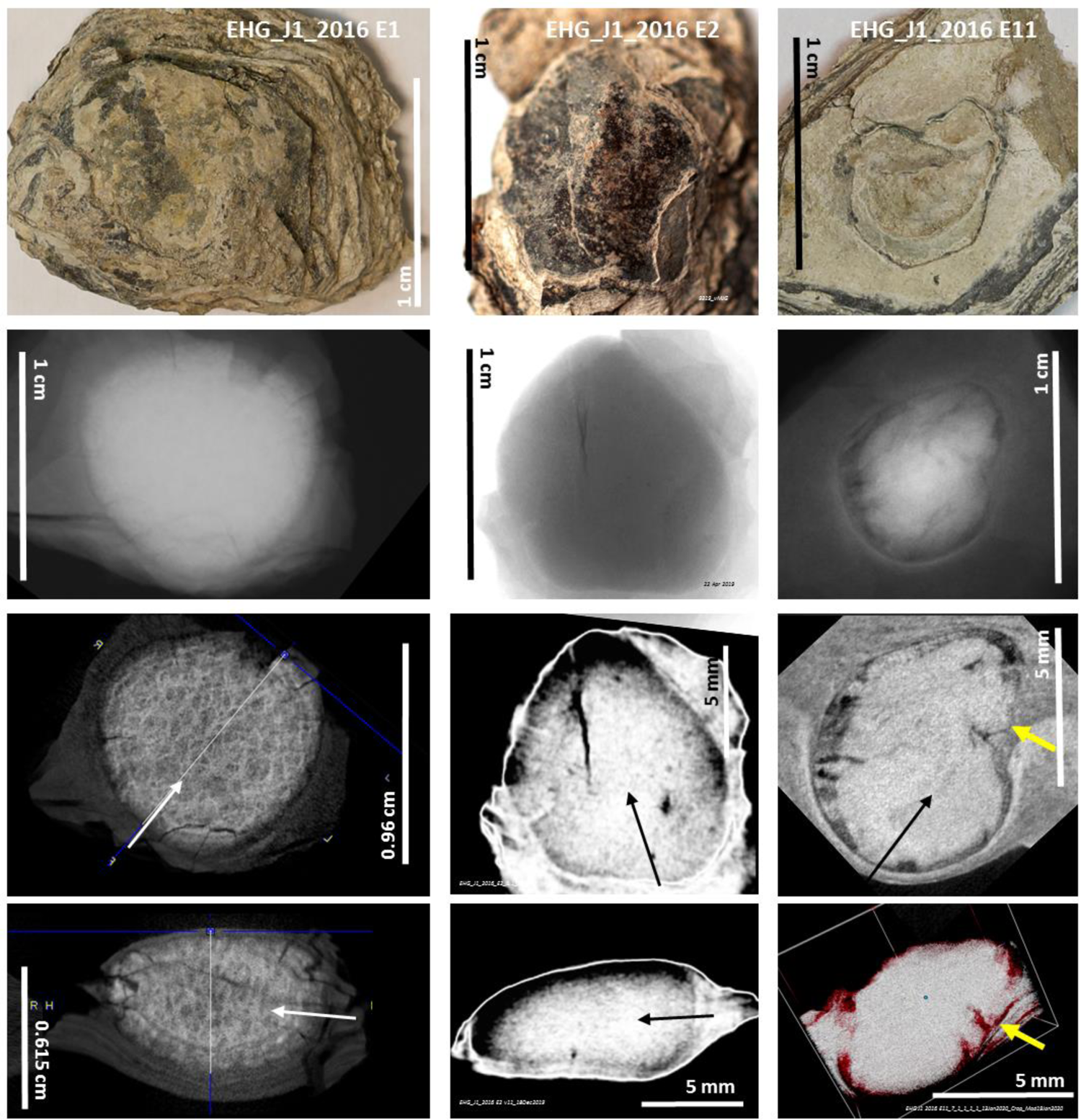
Three eggs extracted from the EHG_J1_2016 fossil. Under macrophotography (top), high resolution X-ray imaging (captured in the chamber of CT scanners; at 73 μm resolution for EHG_J1_2016 E1 and EHG_J1_2016 E2, and at 10 μm for EHG_J1_2016 E11) and micro-CT scanning with coronal and sagittal views (at 73 μm resolution for EHG_J1_2016 E1 and EHG_J1_2016 E2, and at 9 μm for EHG_J1_2016 E11), except for EHG_J1_2016 E11 for which a sagittal view of a 3D image is shown instead of a 2D sagittal view. White and black arrows are used for cross-orientation of different CT scan views of the same sample. A yellow arrow points at the presumed head location of the fossilized fetus with surface morphology preservation of EHG_J1_2016 E11.

**Figure S22.**
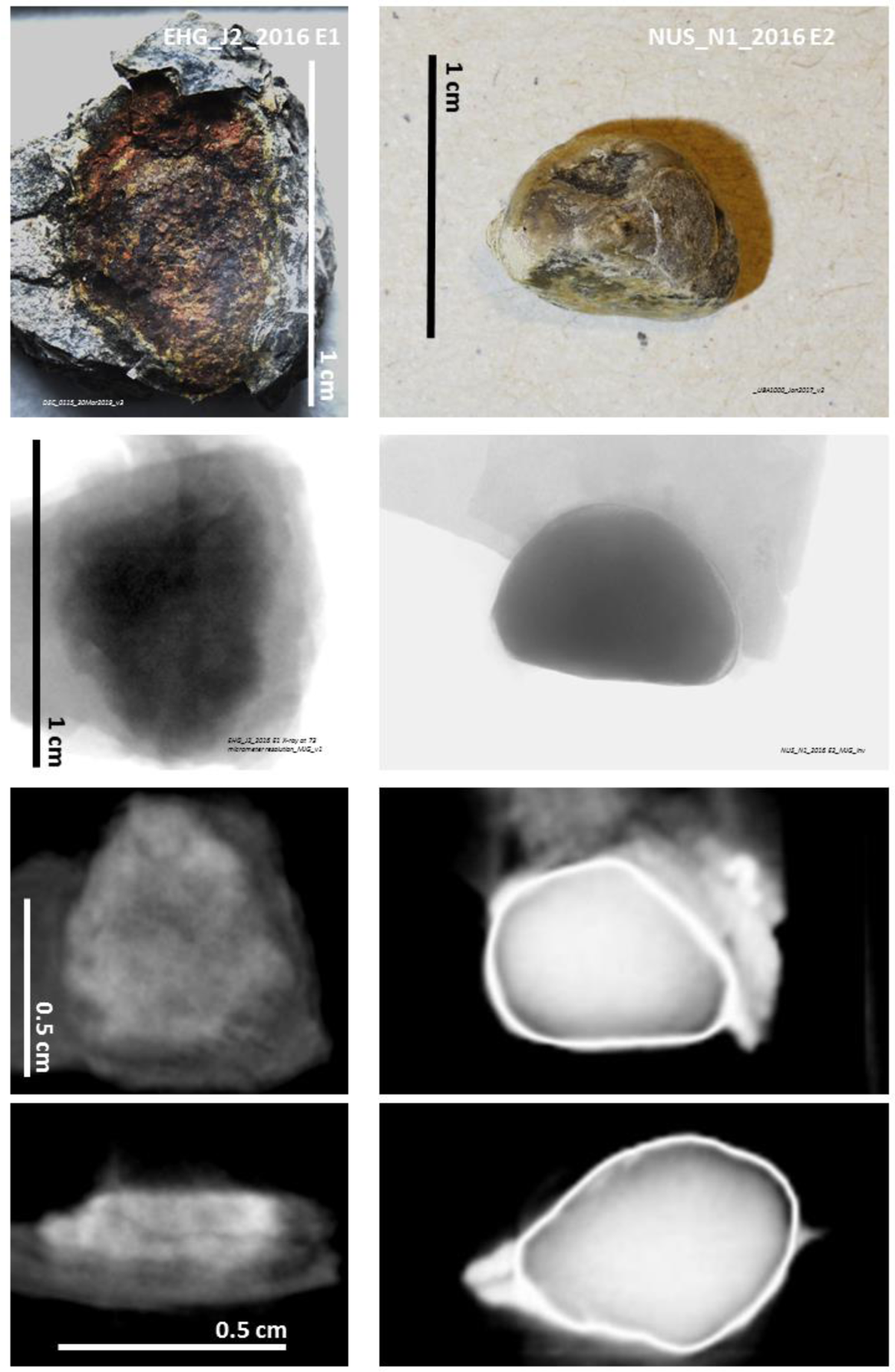
Eggs from the EHG_J2_2016 and NUS_N1_2016 fossils. One egg extracted from the EHG_J2_2016 fossil (top left) and another from the NUS_N1_2016 fossil under macrophotography (top right), high resolution X-ray imaging (captured in the chamber of a CT scanner at 73 μm resolution) and 2D CT scan views at 73 μm resolution.

**Figure S23.**
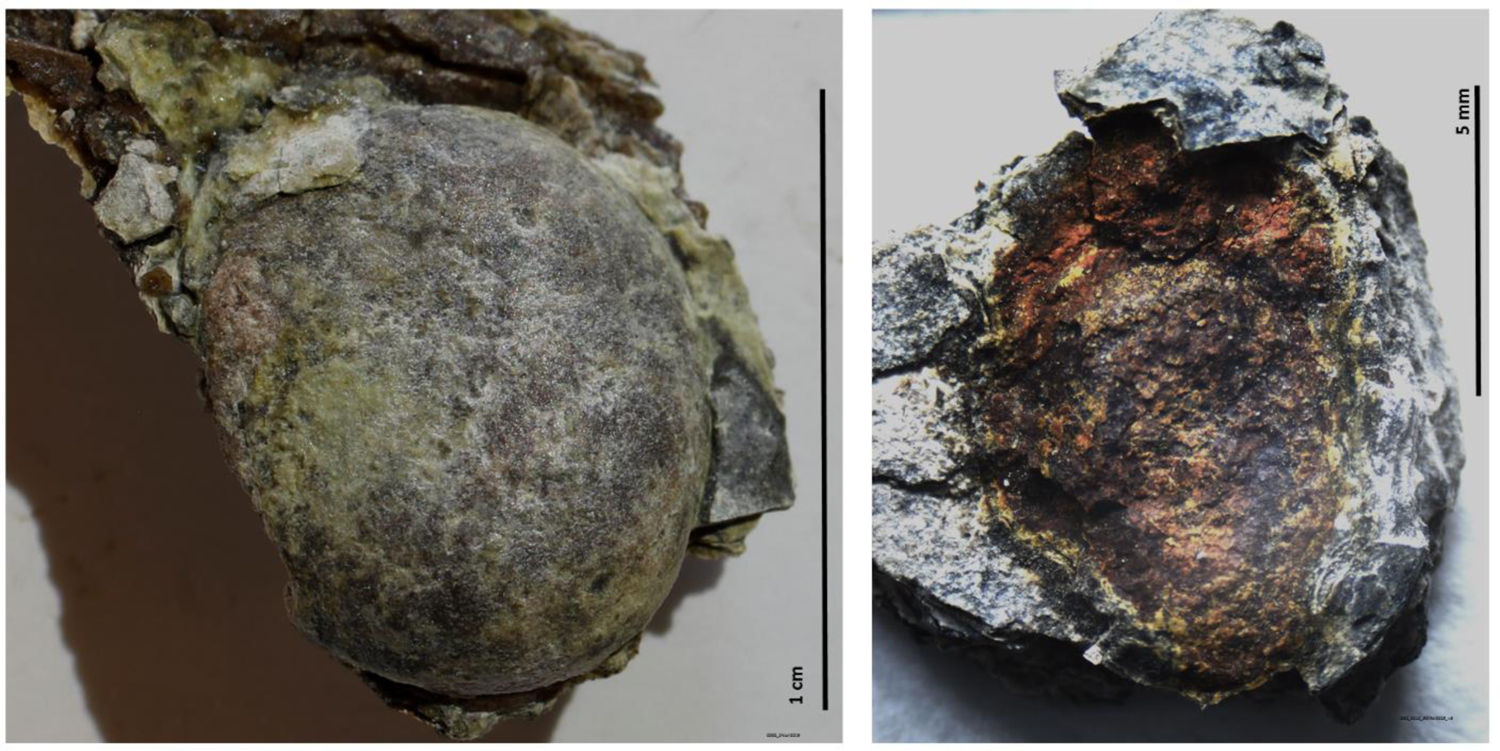
Two eggs extracted from the fossil of NUS_A6_2016 specimen. NUS_A6_2016 E1 (left) derived from the NUS_A6_2016 specimen and EHG_J2_2016 E1 (right) derived from the EHG_J2_2016 specimen. The latter broke apart during preparation.

**Figure S24.**
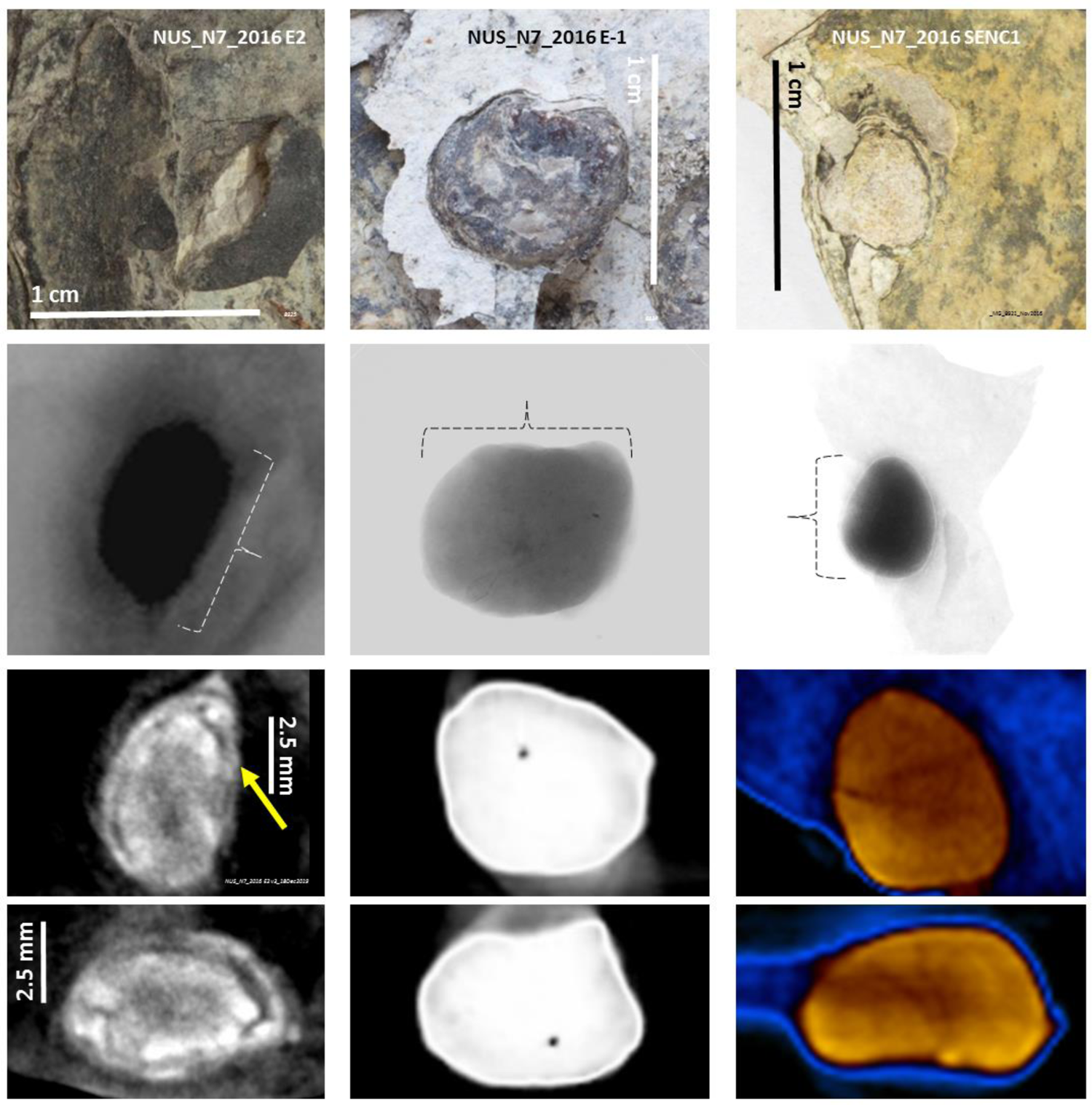
Three eggs extracted from the NUS_N7_2016 fossil. Macrophotography (top), high resolution X-ray imaging (captured in the chamber of a CT scanner at 73 μm resolution) and 2D views of CT scans at 73 μm resolution. It remains to be demonstrated if the yellow arrow points at the beaked head of a fetus with surface morphology preservation in NUS_N7_2016 E2.

**Figure S25.**
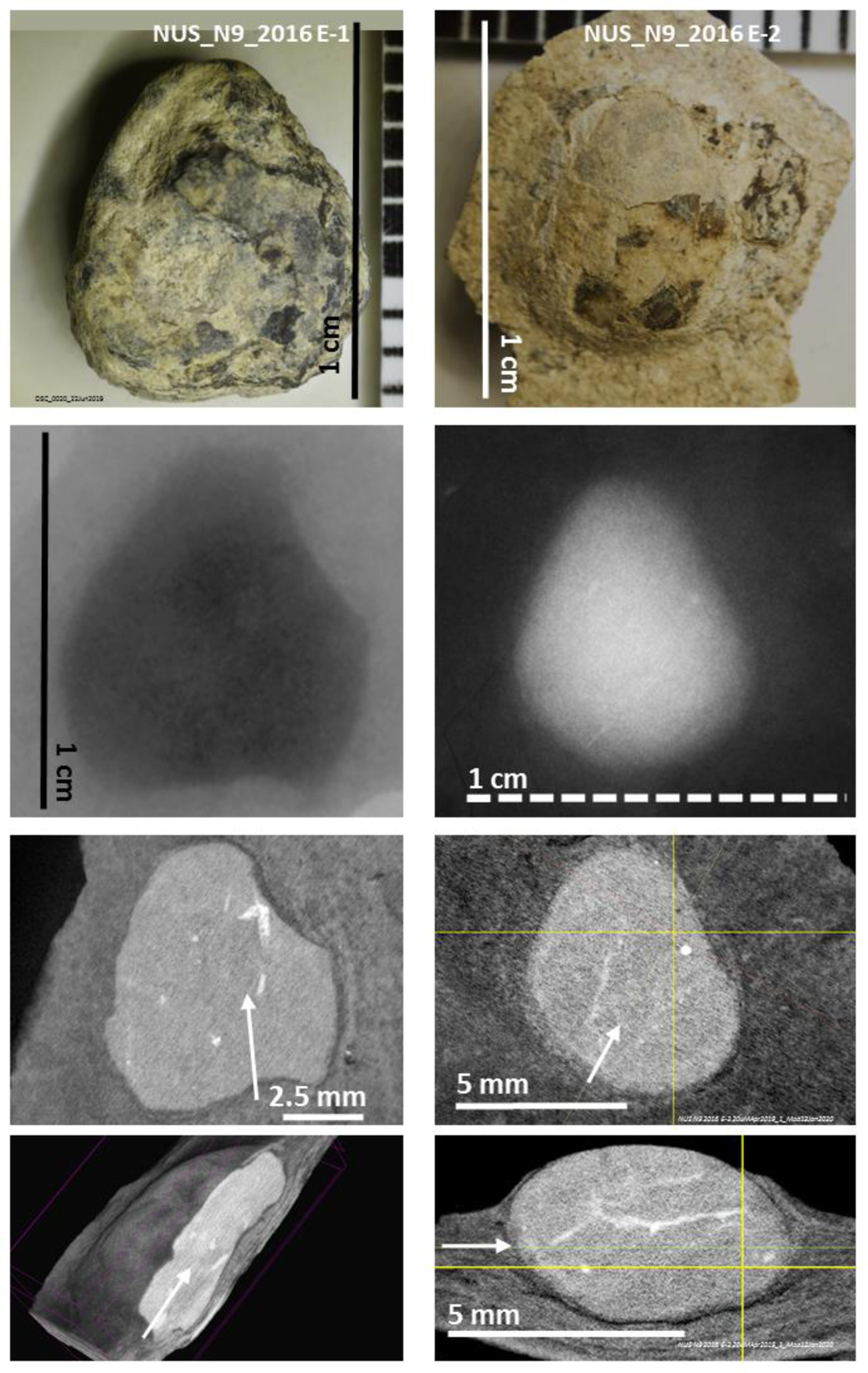
Two eggs extracted from the NUS_N9_2016 fossil. Macrophotography (top), high resolution X-ray imaging (captured in the chamber of a CT scanner) and 2D views of CT scans at 20 μm resolution, except for NUS_N9_2016 E-1 for which a virtual sagittal cut of a 3D view is shown instead of a 2D sagittal view (bottom left). White arrows are used for cross-orientation of different CT scan images of the same sample.

**Figure S26.**
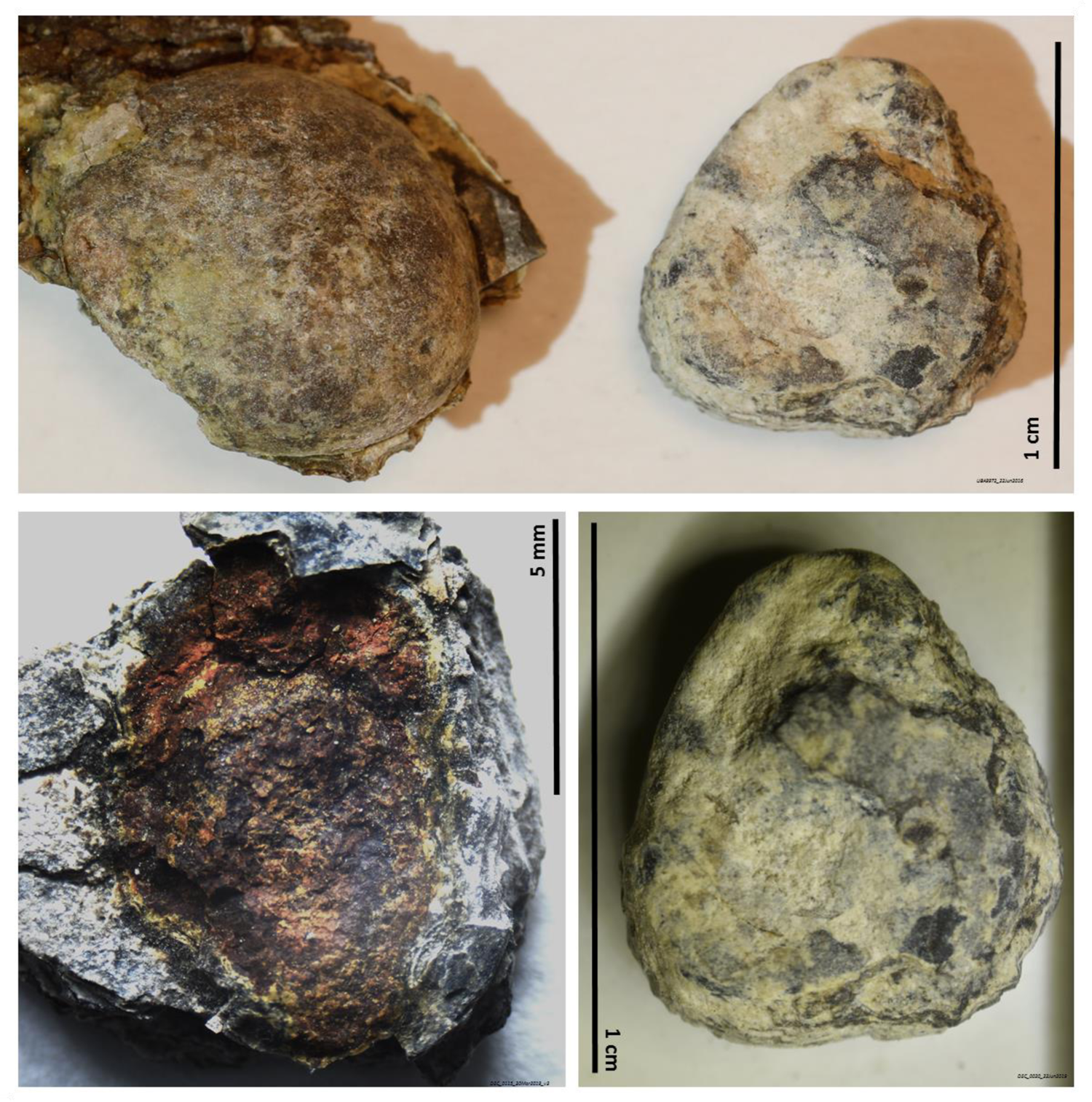
NUS_N9_2016 E-1 egg next to two other eggs derived from the fossils of two other specimens. NUS_A6_2016 E1 derived from the fossil of specimen NUS_A6_2016 is show on top left. NUS_N9_2016 E-1 derived from the fossil of specimen NUS_N9_2016 is shown on the top right and bottom right. EHG_J2_2016 E1 derived from the fossil of specimen EHG_J2_2016 is shown bottom left.

**Figure S27.**
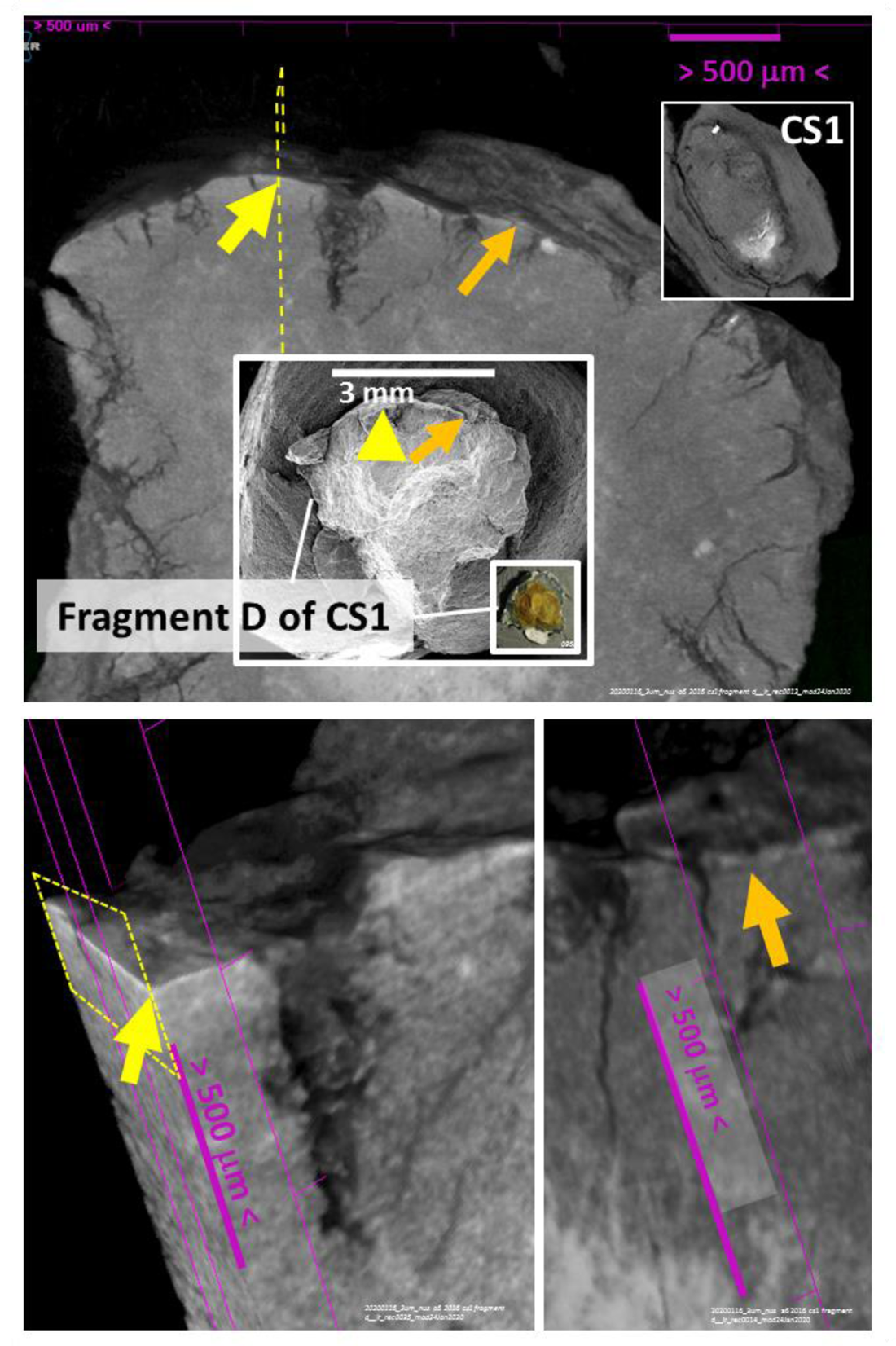
Eggshell in fragment D of the NUS_A6_2016 CS1 egg. 3D views of CT scans at 3 μm resolution. The arrows indicate two different eggshell positions of fragment D of the CS1 egg (the entire egg before breakage is shown in the inset before breakage). A virtual cut is shown on the bottom left at the position indicated by the yellow arrow to expose the eggshell as a continuous surface layer.

**Figure S28.**
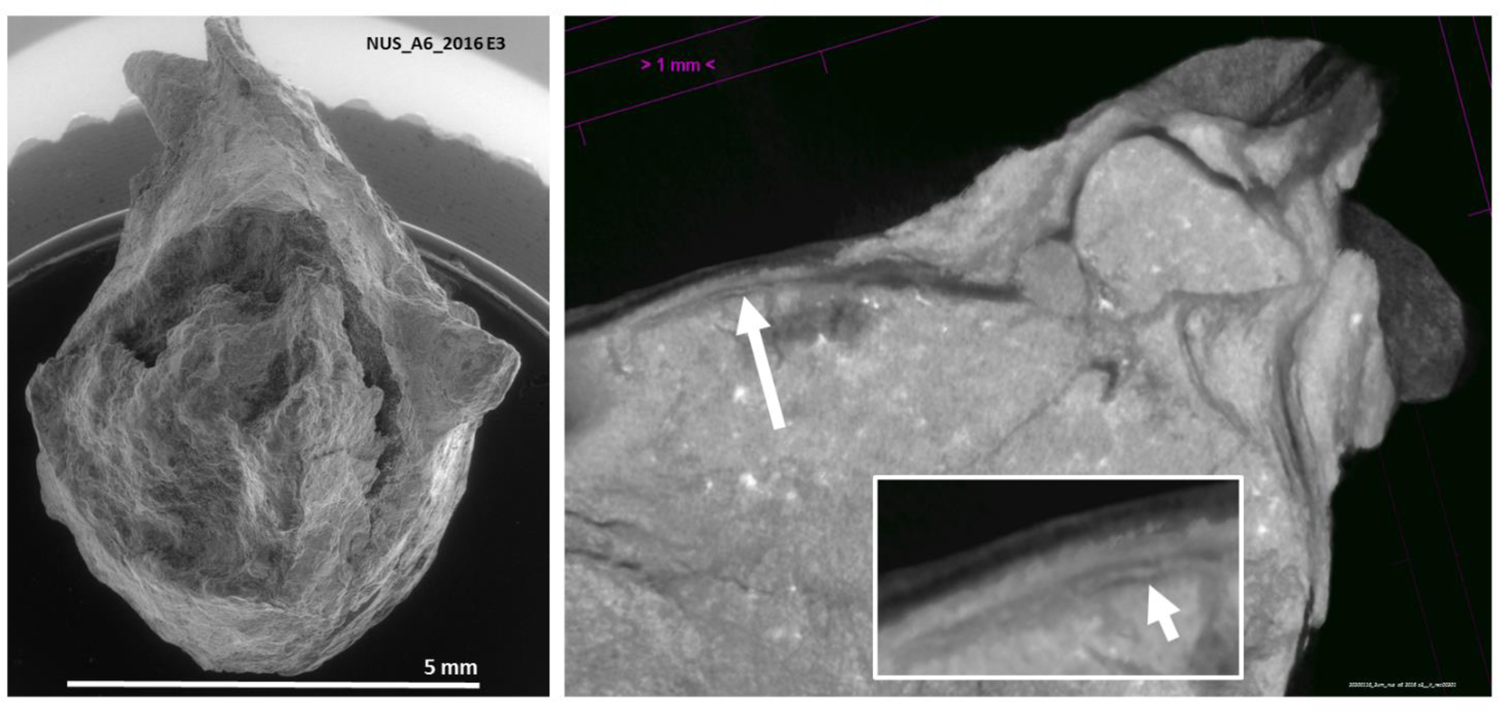
Eggshell in the NUS_A6_2016 E3 egg. 3D views of CT scans at 3 μm resolution. The arrow on the virtual cut shown on the right indicates an eggshell position magnified in the inset, which is not apparent on the egg surface.

**Figure S29.**
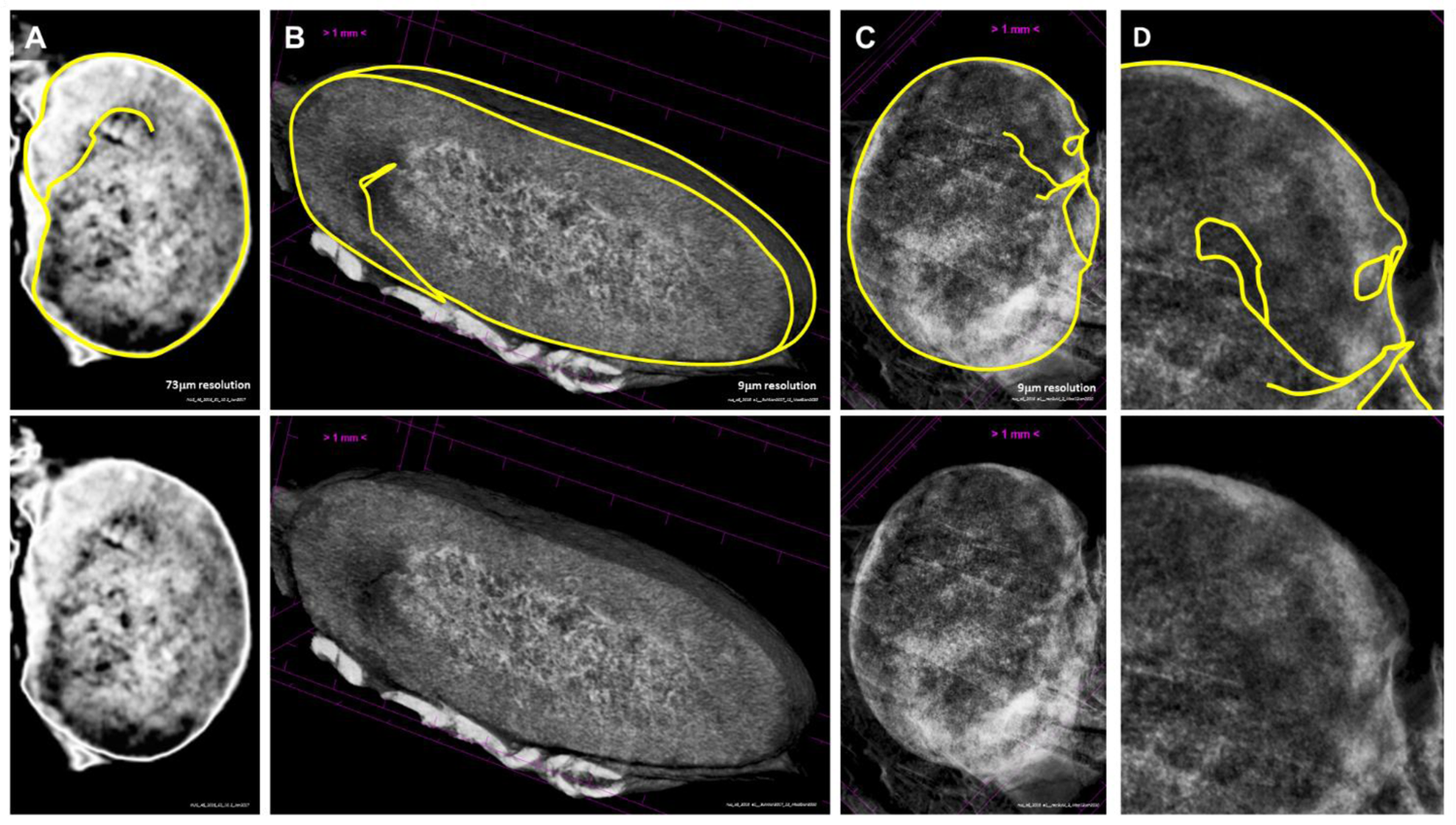
Detection via CT scanning of the preservation of the outer surface morphology of a fetus that resides within the NUS_A6_2016 E1 egg. The NUS_A6_2016 E1 egg is shown under CT scan analysis in coronal view at 73 μm resolution (**A**), together with a sagittal cut of a 3D view at 3 μm resolution (**B**) and a coronal view also at 3 μm resolution (**C**), magnified in **D** to put into evidence the contour of the head of the NUS_A6_2016 E1 fetus, which is outlined in yellow in A-D.

**Figure S30.**
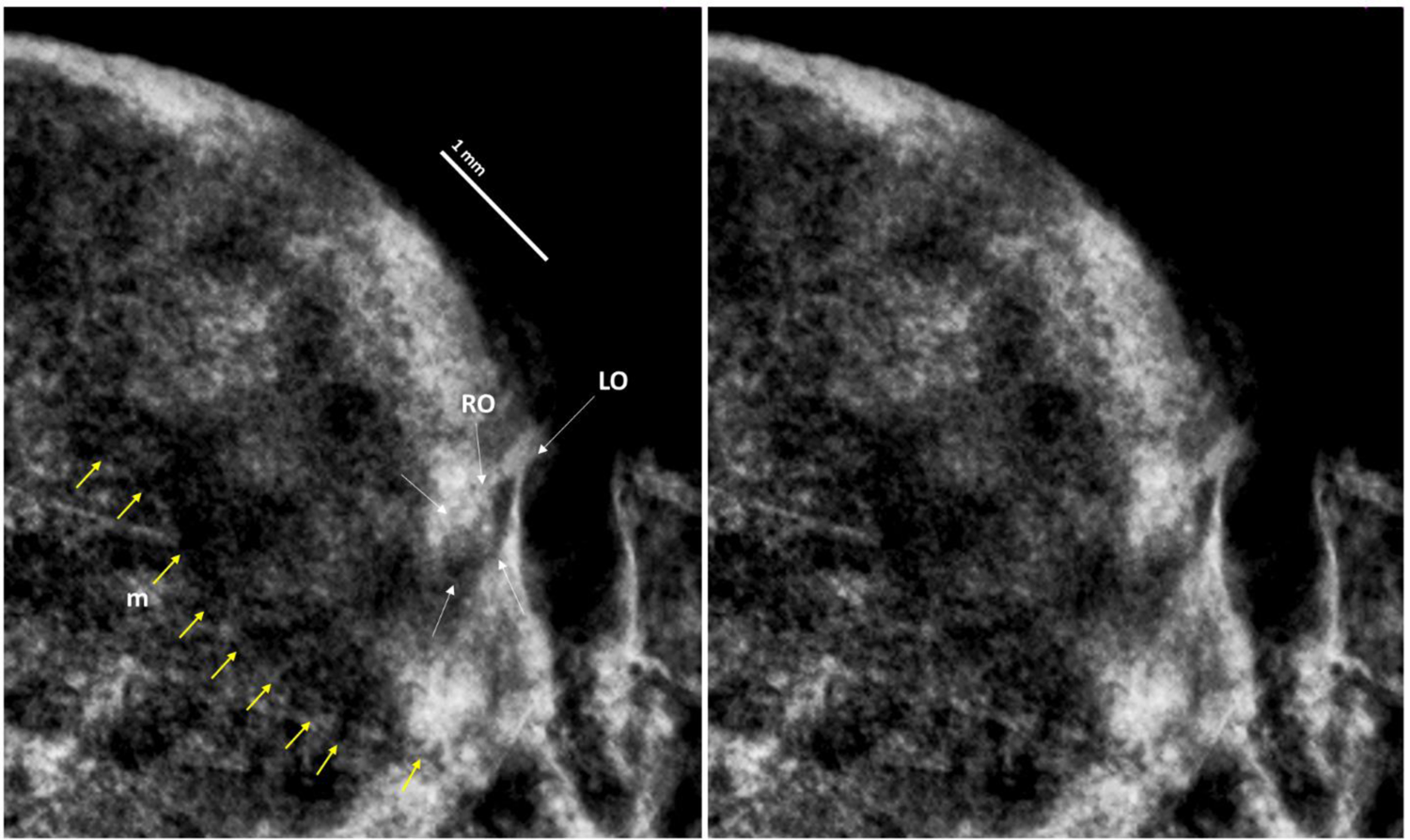
Surface morphology preservation of the head of the fetus enclosed within NUS_A6_2016 E1. Thin yellow arrows point at the lower margin of the mandible and thin white arrows at the right orbit (RO). The left orbit (LO) is also indicated. 3 μm resolution CT scan.

**Figure S31.**
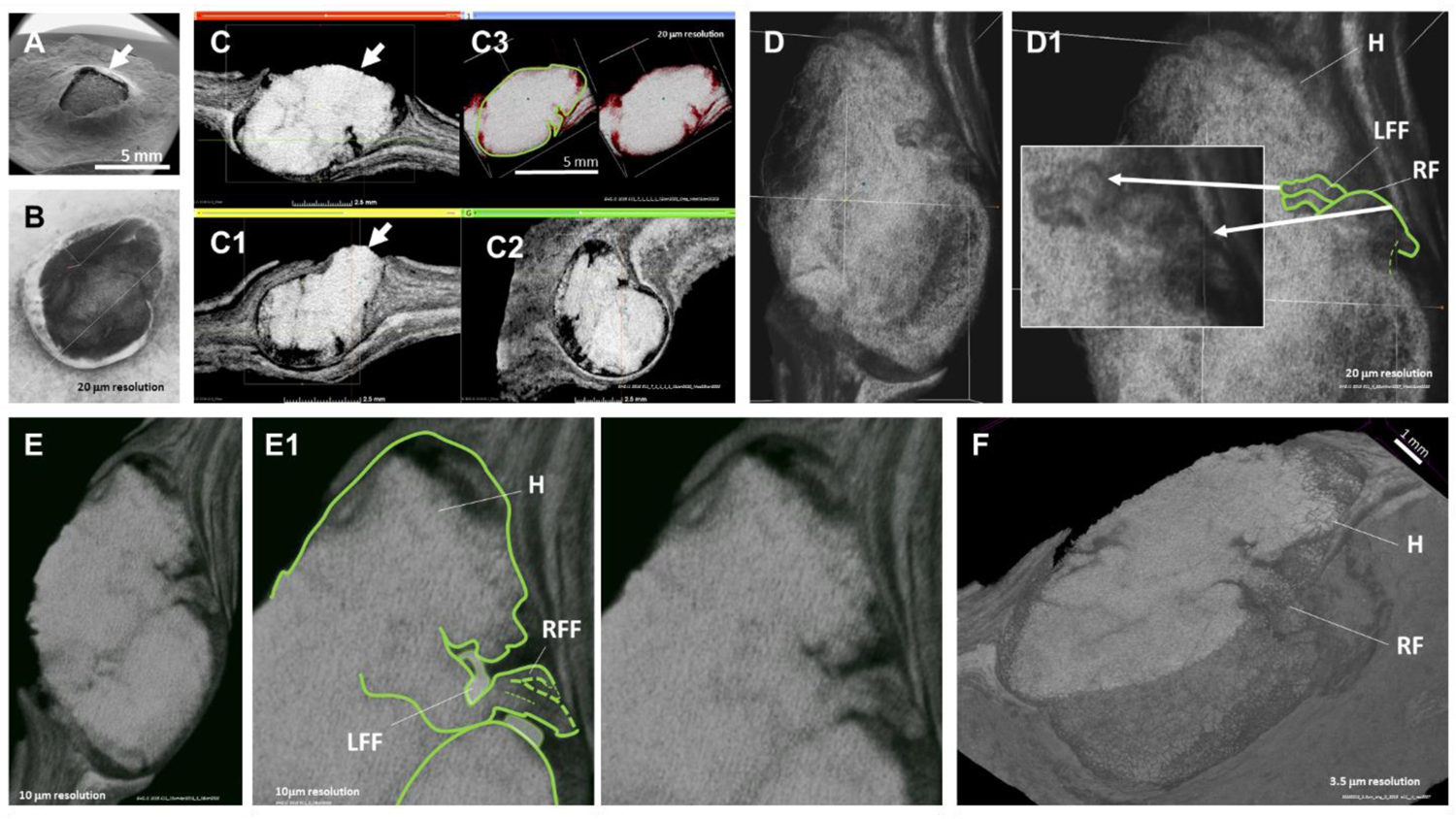
EHG_J1_2016 E11 egg and its enclosed fetus with surface morphology preservation. This egg was identified embedded in its fossil (**A**) with surface erosion (arrow). At 20 μm resolution a space can be identified between the surface of the fetus and the egg periphery (**B**). 2D (**C**, **C1** and **C2**) and 3D (**C3**) views of the egg enclosed fetus are shown under CT scans at 20 μm resolution (the fetus is outlined in green in C3). A 3D image at the same resolution is shown in **D** and enlarged in **D1** to show the outline of two left forelimb fingers. Another 3D image of the fetus is shown under a CT scan at 10 μm resolution (**E**) and enlarged in **E1**, showing the head and the putative outline of its left and right forelimbs. A virtual section of a 3D image of the fetus is shown in **F** under CT scanning at 3 μm resolution. H – Head; LFF – Left forelimb finger; RF – Right forelimb; RFF – Right forelimb finger.

**Figure S32.**
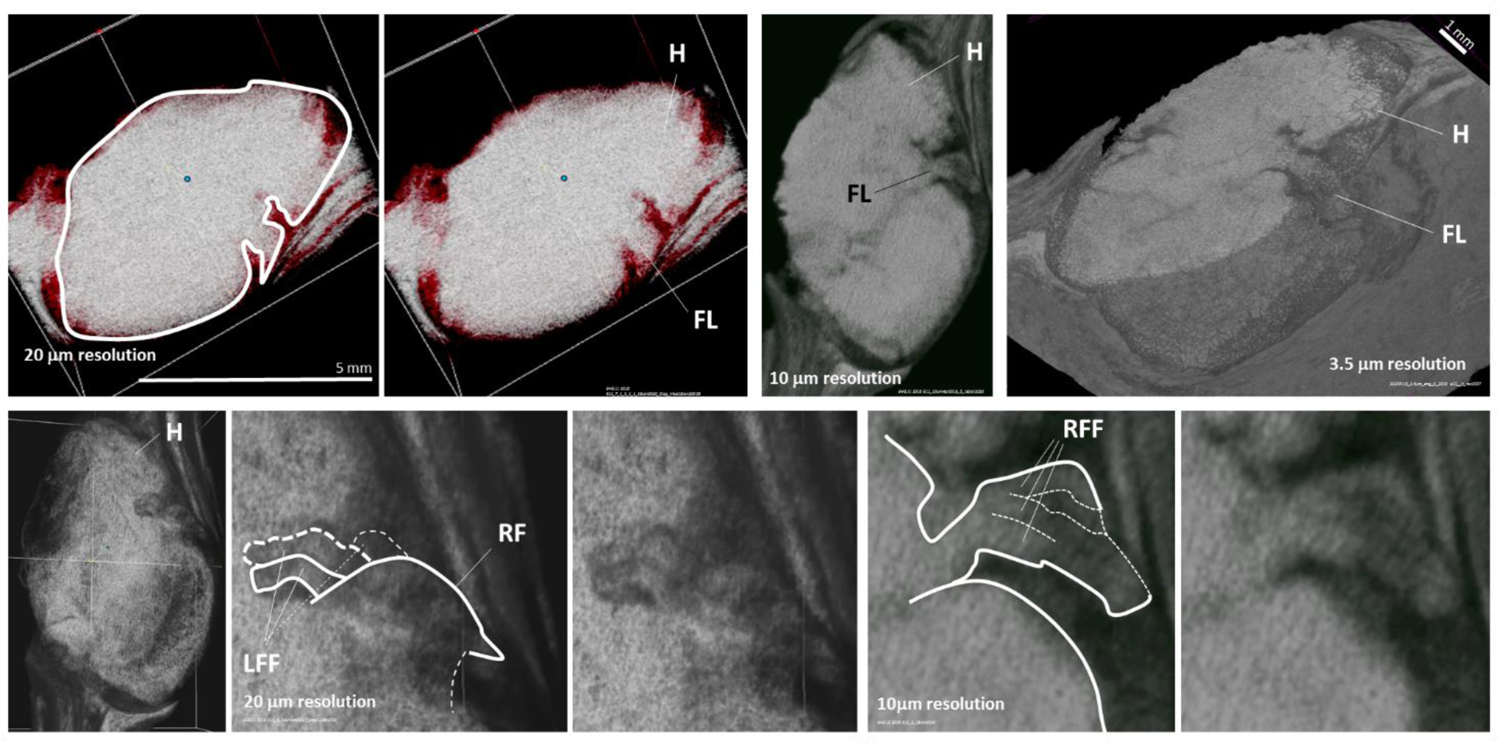
Close look at the fetus enclosed within EHG_J1_2016 E11. CT scan resolutions are indicated. H – Head; FL – Forelimb; LFF – Left forelimb fingers; RF – Right forelimb; RFF – Right forelimb fingers.

**Figure S33.**
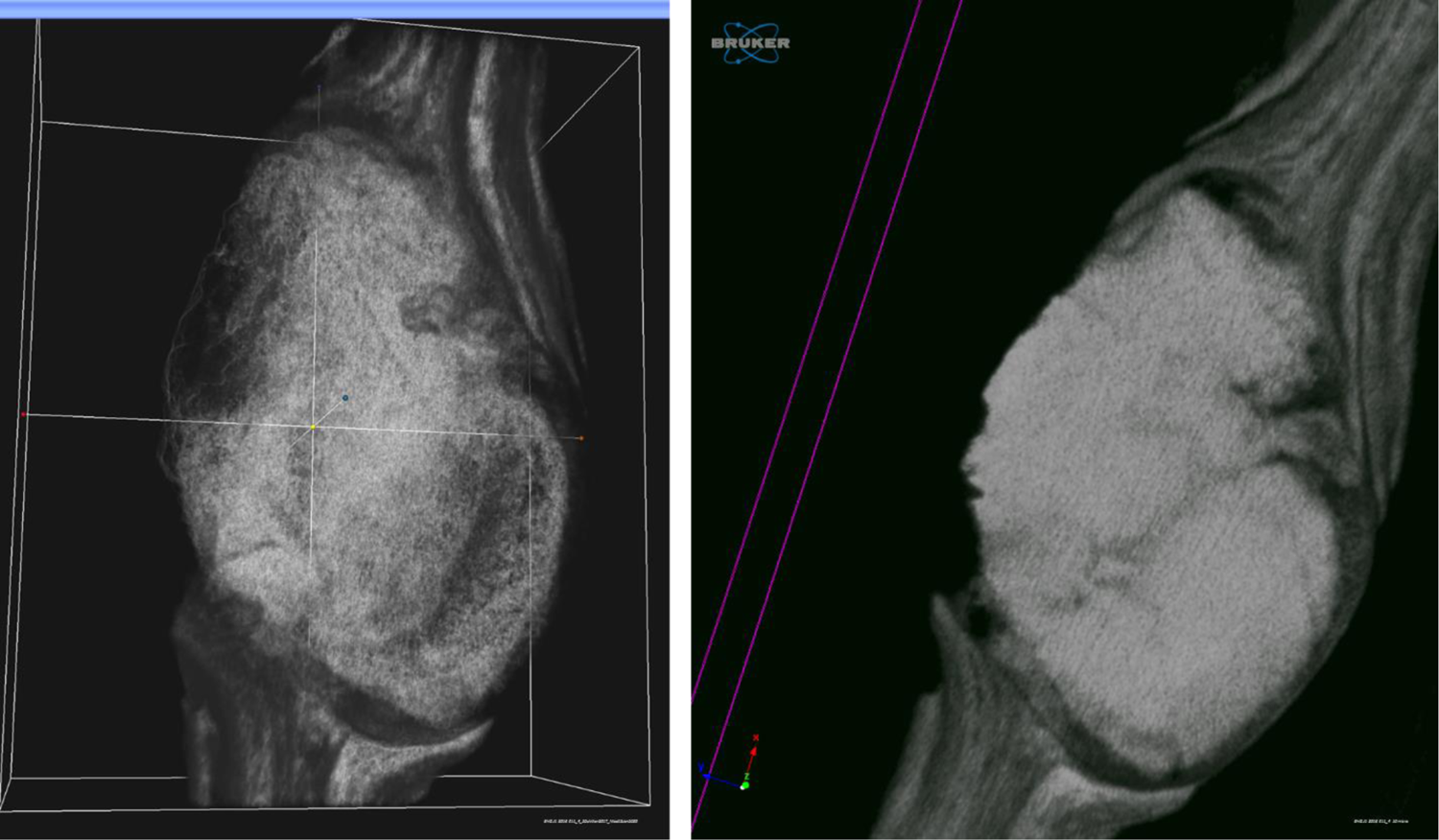
3D micro-CT scan views of the fetus enclosed within EHG_J1_2016 E11. Images are shown at 20 μm (left) and 10 μm resolution (right).

**Figure S34.**
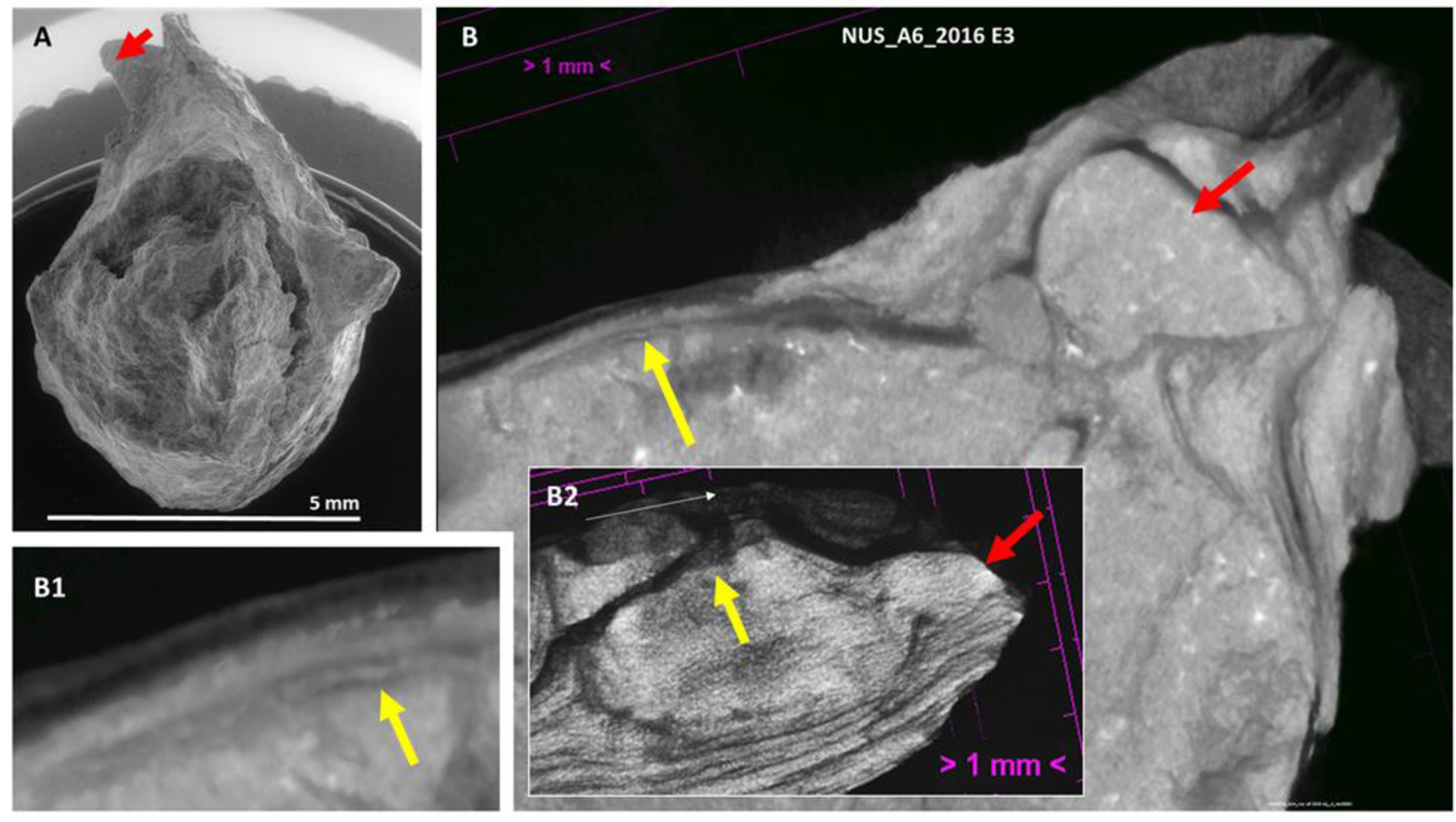
NUS_A6_2016 E3 egg. The egg is shown under SEM (**A**) and virtual 3D CT cuts performed at 3 μm resolution in **B**, **B1** and **B2**. The yellow arrow points at the eggshell, which lies beneath the exposed surface. The red arrow points at what might be the head of the hatching hatchling with surface morphology preservation (see also Figures S35-S37).

**Figure S35.**
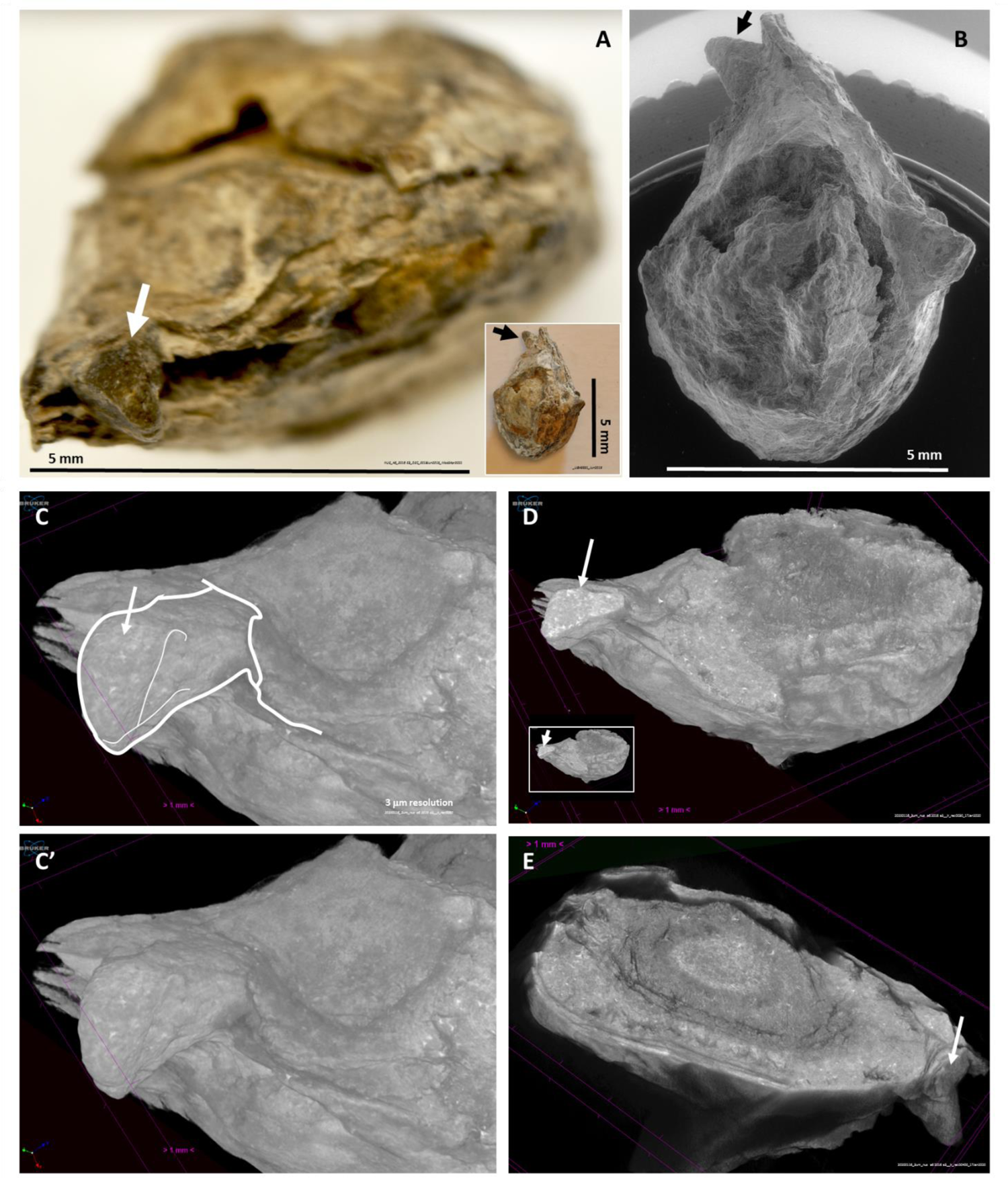
NUS_A6_2016 E3 egg. This egg is shown under macrophotography (**A**) and SEM (**B**). The arrows indicate what may correspond to the protruding head of the hatching hatchling (see also Figs S34 and S36). The putative head is shown in a CT scan at 3 μm resolution (**C**). Virtual coronal cuts at the same resolution are shown in **D** and **E** (see also Figs S41 and S42).

**Figure S36.**
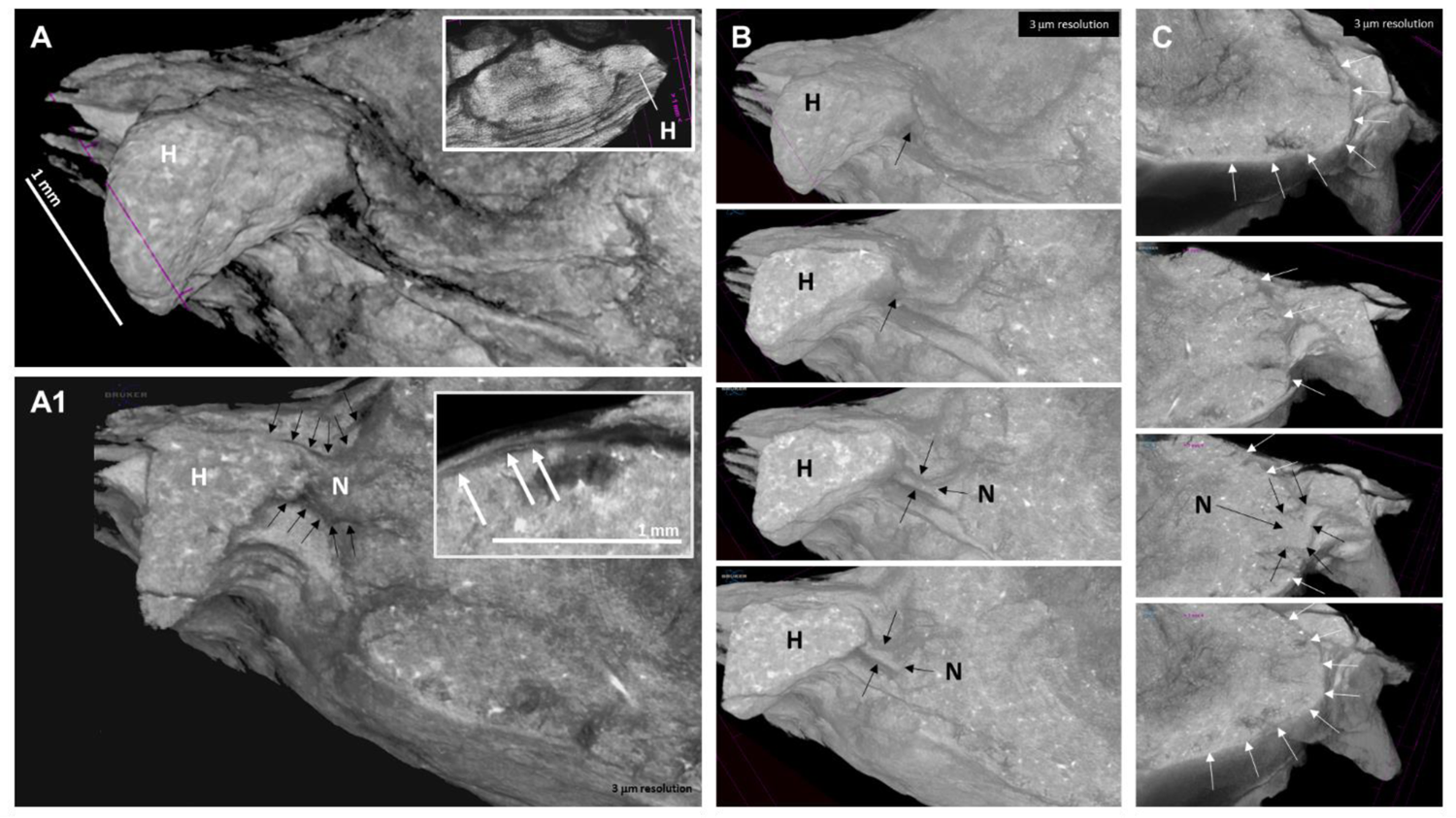
NUS_A6_2016 E3 under 3D CT scan views at 3 μm resolution. The area that may correspond to the protruding head (H) of a hatching animal is indicated in A in 3D and under a virtual cut in A1. Additional virtual cuts, performed from top to bottom, are shown in B and C to show that a segment compatible with that of a neck (N) with surface morphology preservation can be observed. Note that both the top and the bottom image in C show the eggshell as a continuous layer, but not the sections in the center, where the putative neck is visible as an extruding structure that interrupts the continuity of the eggshell in the plane of the virtual cut.

**Figure S37.**
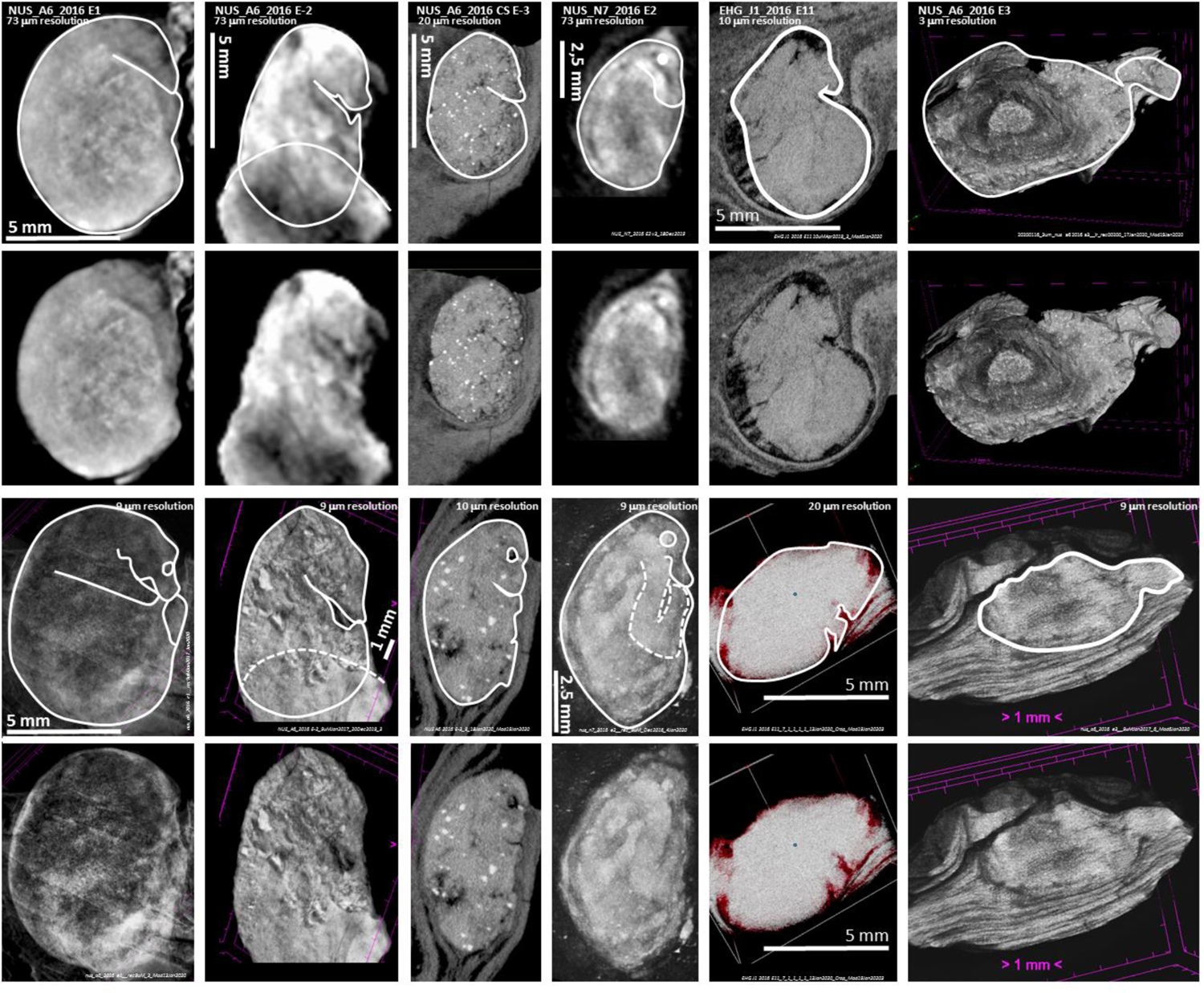
Six eggs with enclosed fetuses characterized by surface morphology preservation. Eggs are shown in coronal CT view (top two rows), except for NUS_A6_2016 E3 (far right), which is shown in a virtual 3D sagittal view, and under 3D CT scan view (bottom two rows). Fetuses with surface morphology preservation are outlined (NUS_A6_2016 E3, in the far right, may have fossilized at the hatching stage; see also Figures S34-S36).

**Figure S38.**
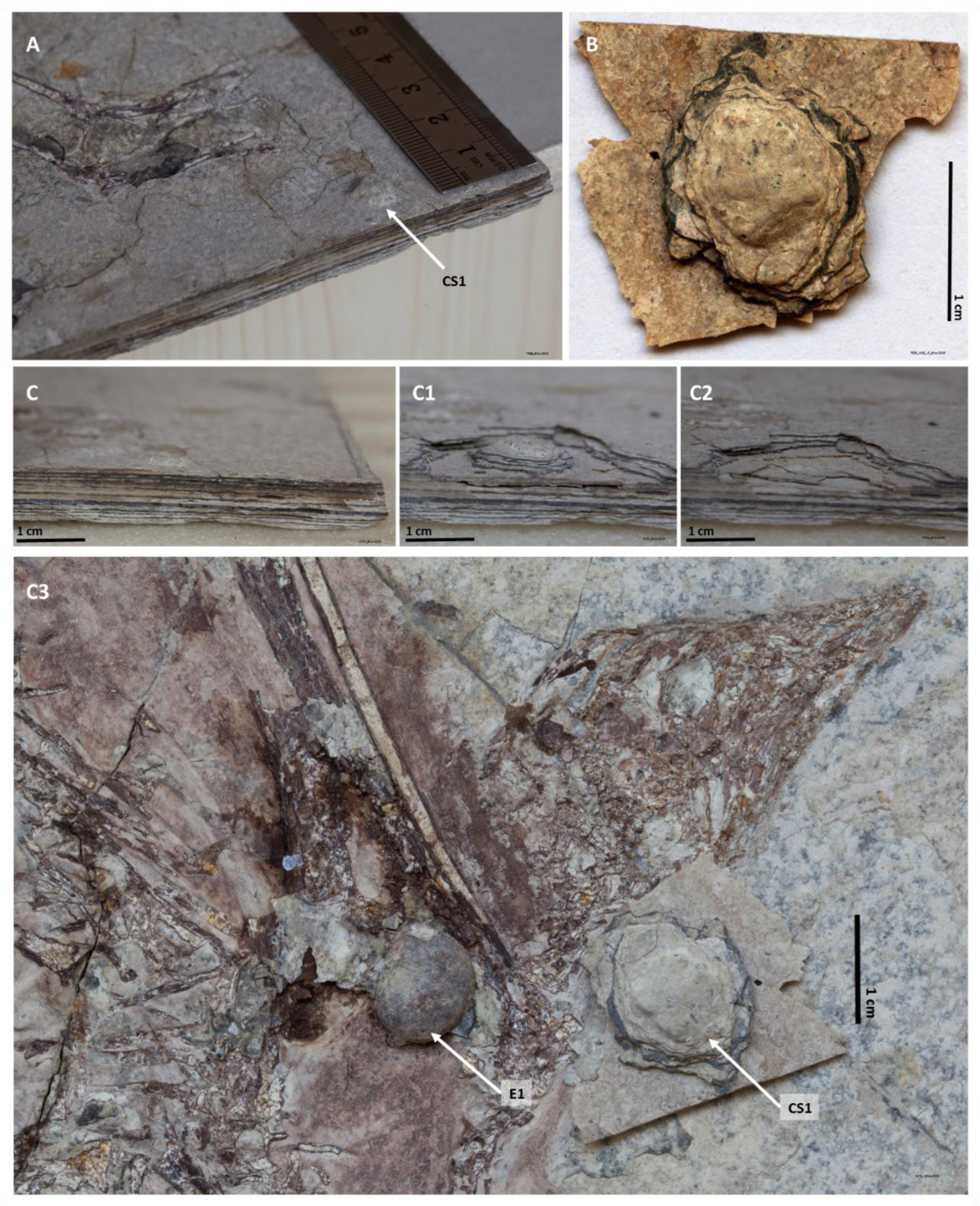
The NUS_A6_2016 CS1 egg. The NUS_A6_2016 CS1 egg is shown in its original location in the counterslab (**A**), after its extraction from the slab (**B**) and a lateral view of the margin of the same slab before, during and after extraction of this egg (**C**, **C1** and **C2**, respectively). For size and appearance comparison, the NUS_A6_2016 CS1 egg extracted from the counterslab is also shown placed to the right of the NUS_A6_2016 E1 egg on the surface of the main slab of the same fossil (**C3**).

**Figure S39.**
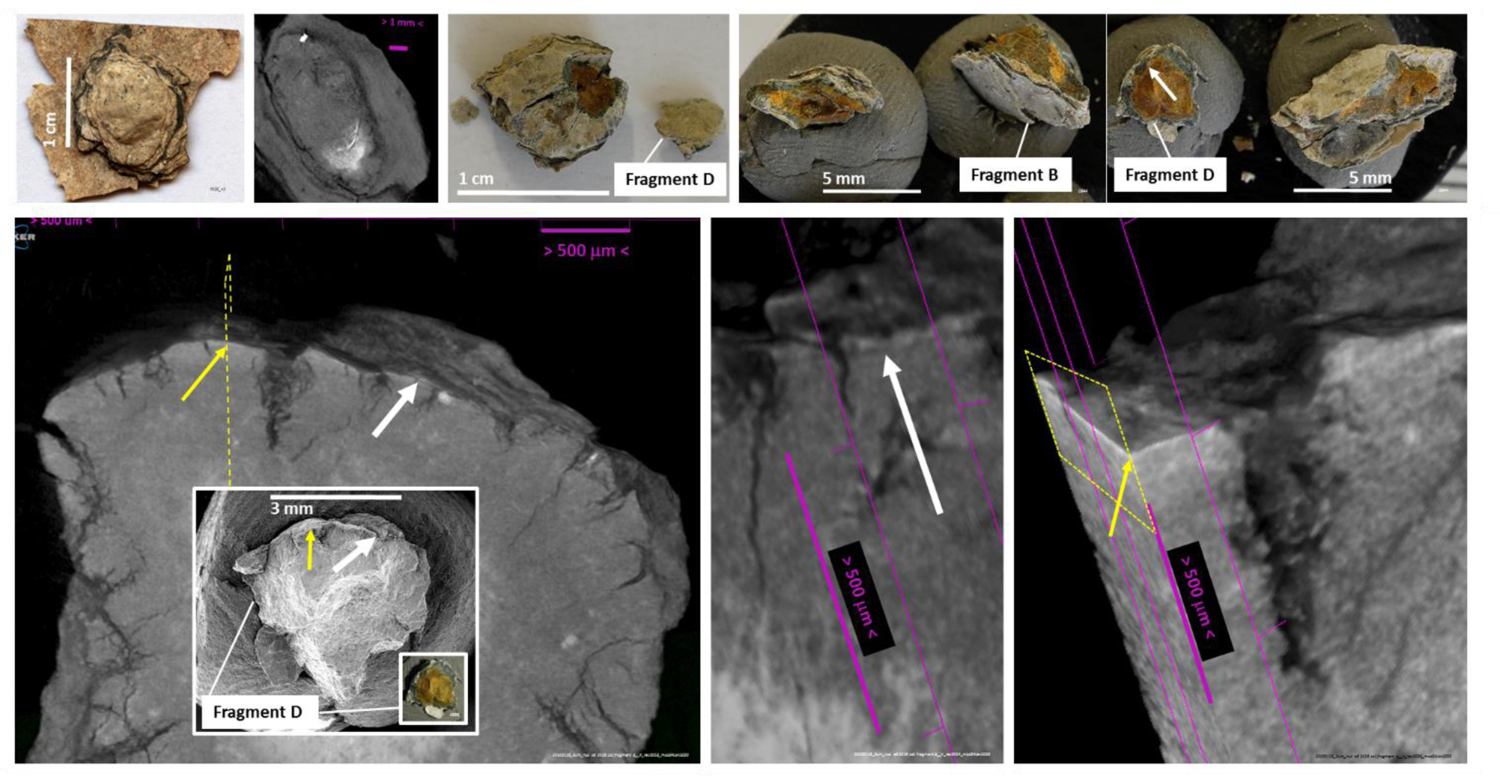
Detection of the eggshell in Fragment D of NUS_A6_2016 CS1. This egg is shown under macrophotography after its extraction from the NUS_A6_2016 fossil (top far left) and under CT scan analysis (top left). It was broken for eggshell exposure in radial view (top center left, top center right and top right). Fragment B and Fragment D are indicated. The area of Fragment D where the eggshell was identified via SEM and EDS is indicated (white arrow on the top right image). Virtual cuts of 3D views of Fragment D and identification of the eggshell in two locations of this egg fragment are indicated by white and yellow arrows (bottom). Close-ups of the areas of the eggshell indicated by white and yellow arrows are provided on the bottom right (see also Figure S40). CT scans at 3 μm resolution.

**Figure S40.**
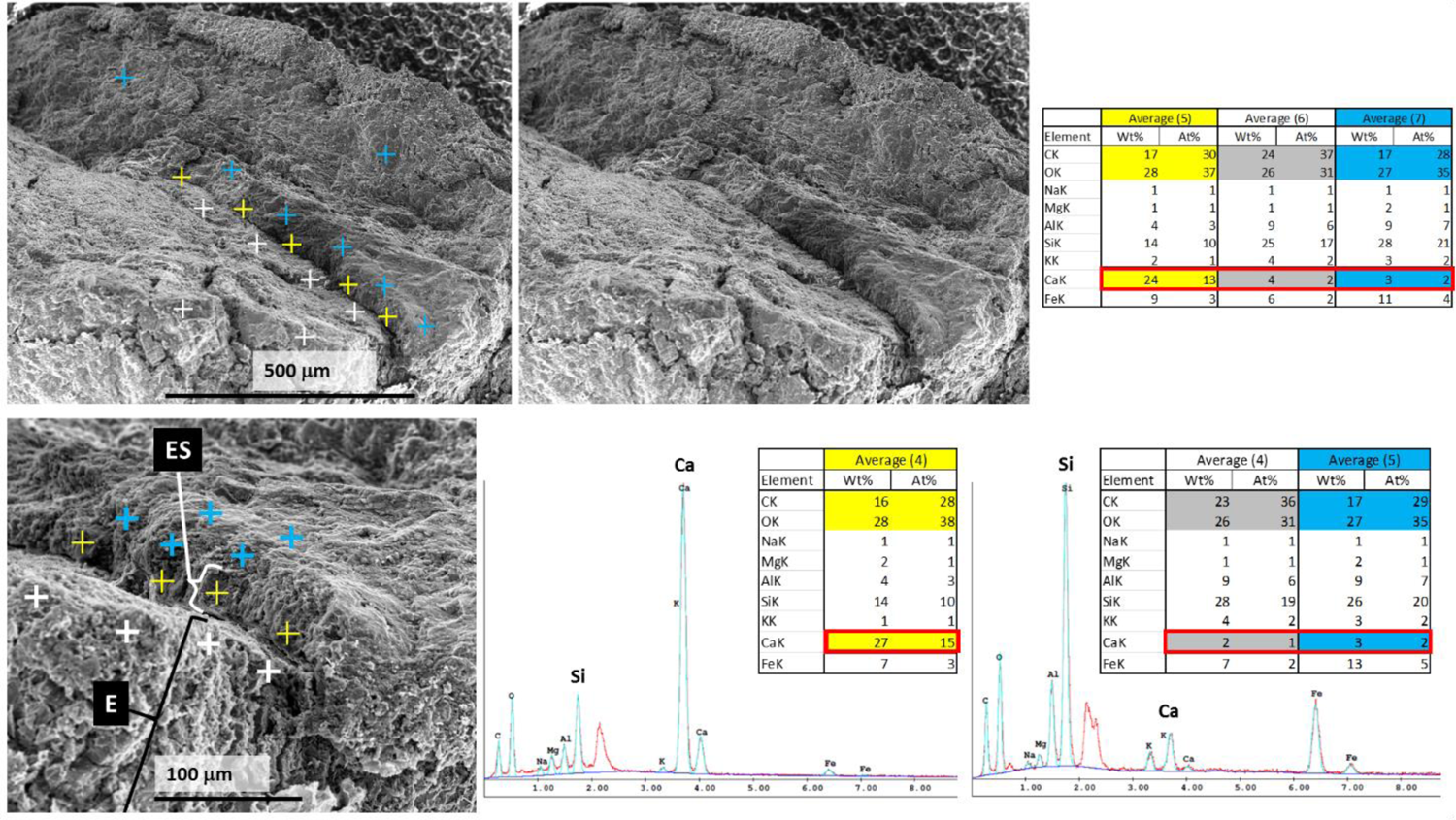
Ornithuromorph eggshell under EDS analysis. **Top**: SEM and EDS analysis of the eggshell detected in Fragment D of the NUS_A6_2016 CS1 egg (same area shown in Figure S39) with average values displayed in table format for points collected above the eggshell (blue crosses), at the eggshell (yellow crosses) and inside the egg, below the eggshell (white crosses). **Bottom**: analysis at higher magnification with average values displayed in table format together with a graphic view, for profile comparison, of one the yellow points (bottom center) and one non-eggshell point (bottom right). E – Egg interior; ES – Eggshell.

**Figure S41.**
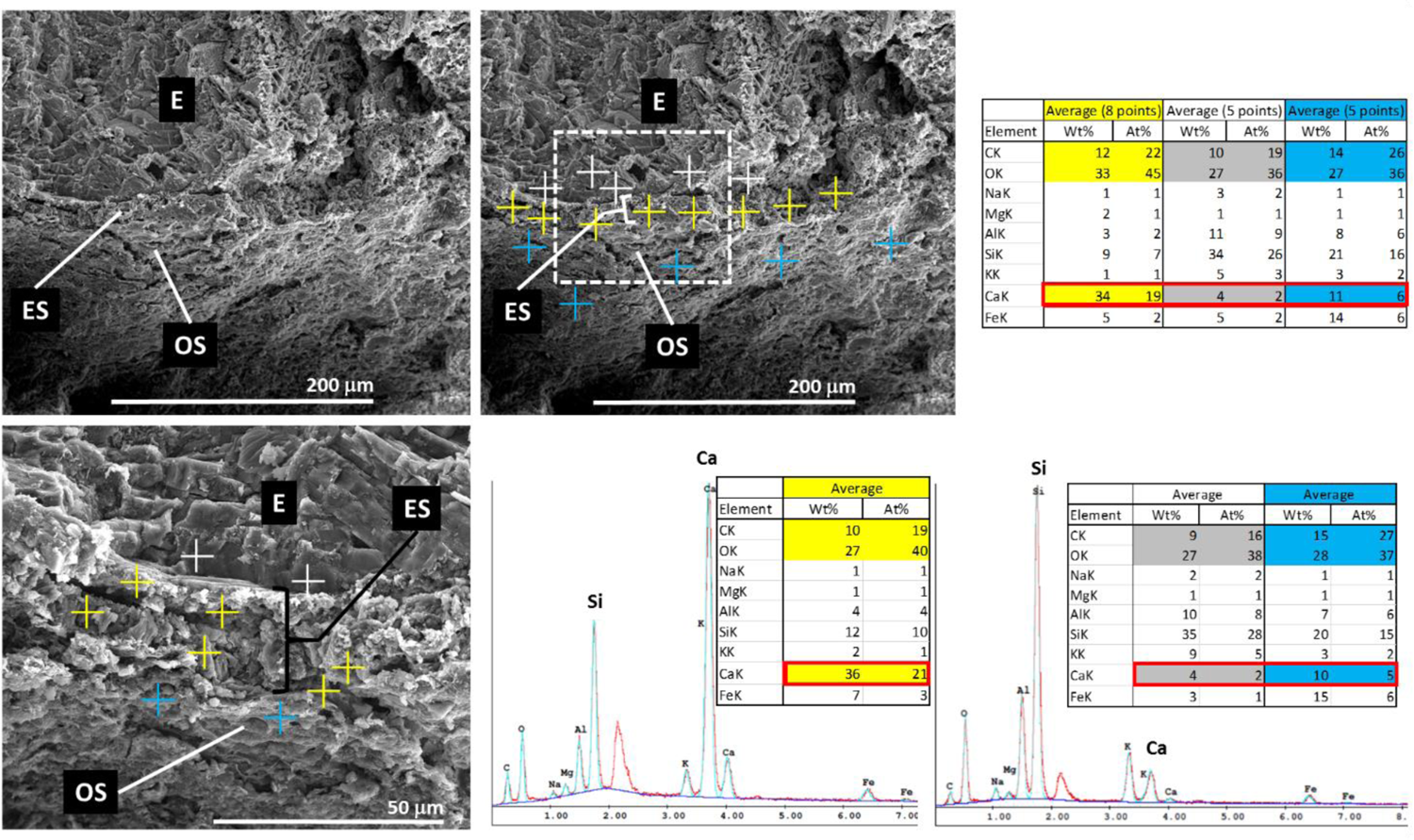
Ornithuromorph eggshell under EDS analysis. **Top**: Radial view of the eggshell in Fragment B of the NUS_A6_2016 CS1 egg. This area was submitted to EDS analysis at the indicated points (top center). Average values collected from the indicated points in the outer surface (blue crosses), within the eggshell thickness (yellow crosses) and from inside the egg (white crosses) are displayed in table format on the top right highlighting carbon, oxygen and calcium readings (the latter with red outline). EDS analysis was also performed at higher magnification in the same eggshell area (bottom). Average values from the points represented in the bottom left image are colored as on top. EDS readings from this magnified area are shown, to permit profile comparison, in graphic view for one the yellow eggshell points next to a table view for such points (bottom) and also in graph view for a blue point located outside the eggshell thickness next to a table view of such points (column with blue cells) and points located inside the egg (column with grey cells). E – Egg interior; ES – Eggshell; OS – Outer surface of the egg.

**Figure S42.**
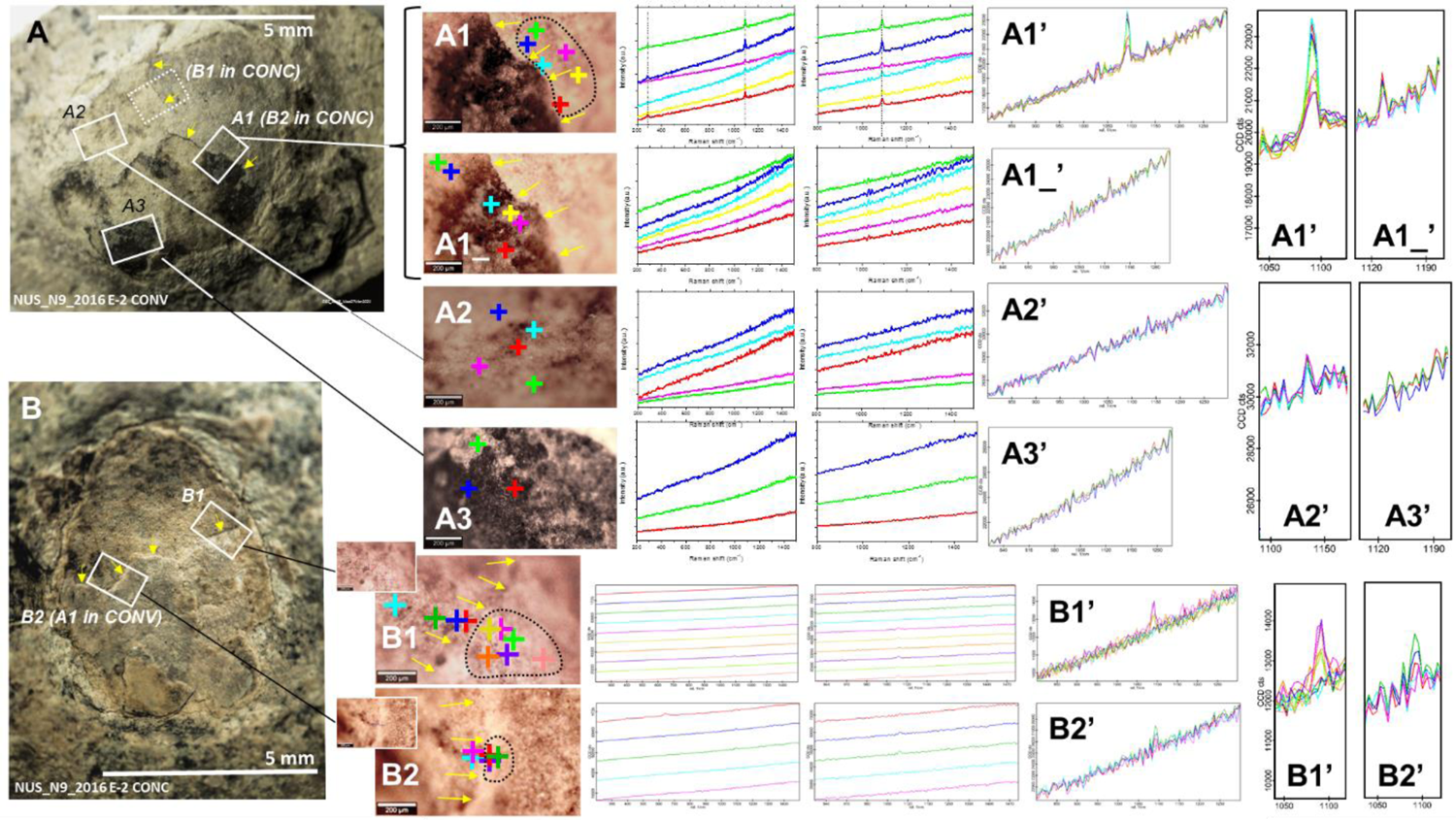
Detection of the eggshell and its limit in the convex surface of the NUS_N9_2016 E-2 egg, which preserved part of the eggshell, and in the corresponding positions in the concave surface of a fragment that was detached from the egg. **A**: Convex surface of the egg NUS_N9_2016 E-2. Areas **A1**, **A2** and **A3** were subjected to RAMAN spectroscopy. Signature calcite modes are consistently observable in **A1’** but not in **A1_**, **A2** and **A3**. **B**: Concave surface of the fragment detached from the convex surface of the NUS_N9_2016 E-2 egg via preparation. Areas **B1** and **B2** correspond, respectively, to an area indicated in dotted line and to area A1 in A. Calcite modes were predominantly and consistently detected only on the side of B1 and B2 where points are surrounded by a block dotted line. These findings were confirmed by EDS mapping as shown in Figures S43-S50.

**Figure S43.**
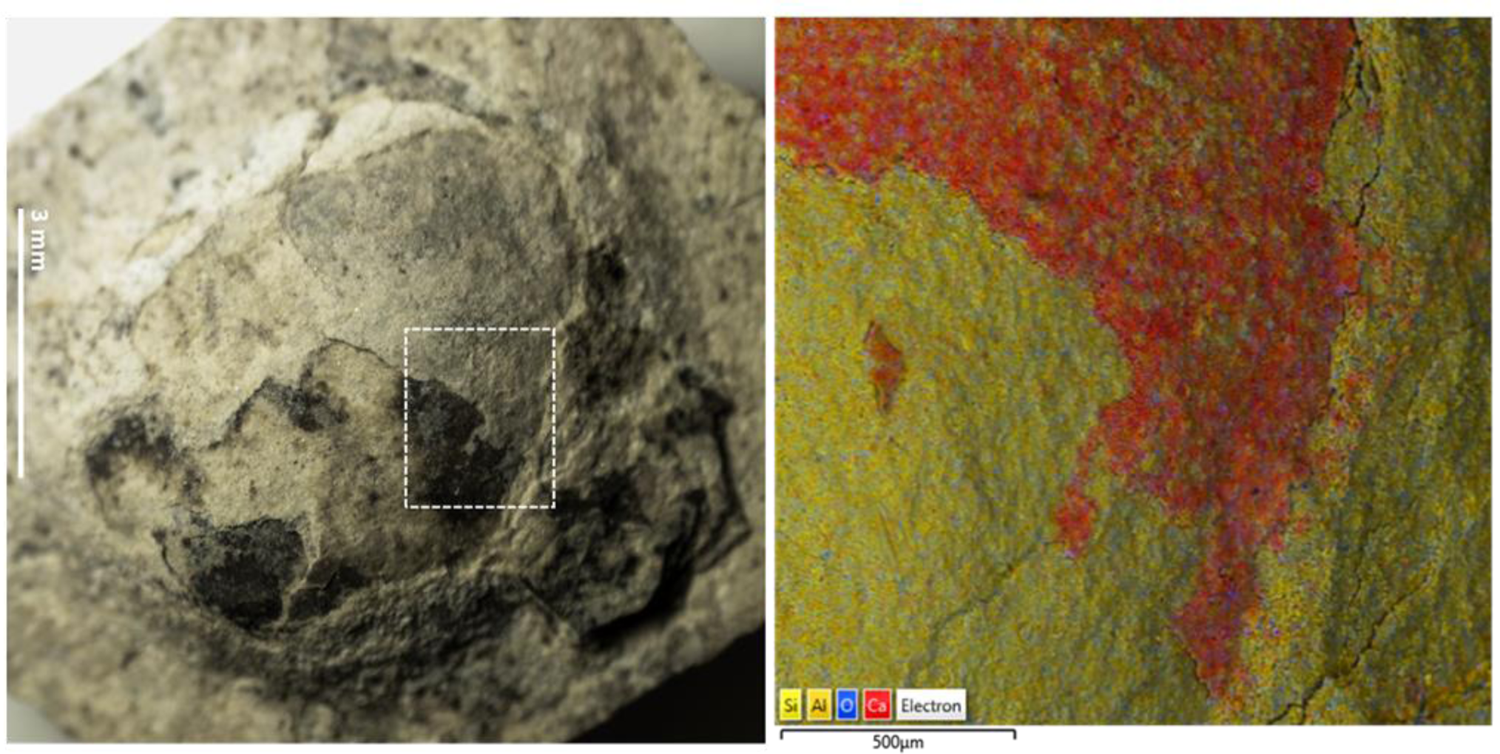
Detection of the eggshell on the convex surface of the NUS_N9_2016 E-2 egg via EDS mapping. Area of the convex surface of NUS_N9_2016 E-2 (left) analyzed via EDS mapping (right). The mapped area includes Area A1 shown in Figure S42, but is significantly larger. Calcium is shown in red and silica in yellow in the EDS map.

**Figure S44.**
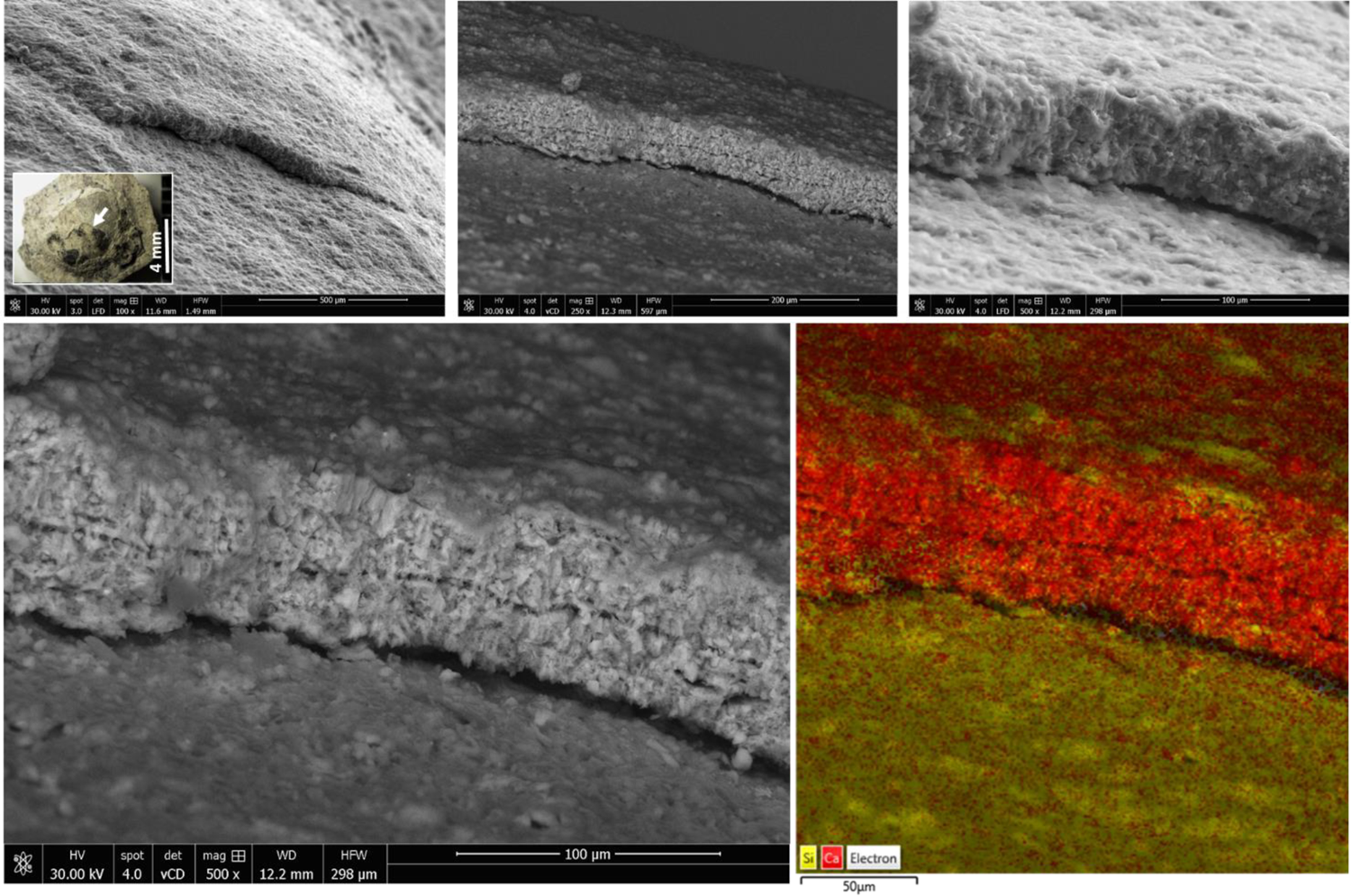
Detection of the eggshell on the convex surface of the NUS_N9_2016 E-2 egg via SEM and EDS mapping. The area of the convex surface of NUS_N9_2016 E-2 indicated by the arrow in the inset on the top left is progressively enlarged in SEM images and analyzed via EDS mapping (bottom). Calcium is shown in red and silica in yellow in the EDS map.

**Figure S45.**
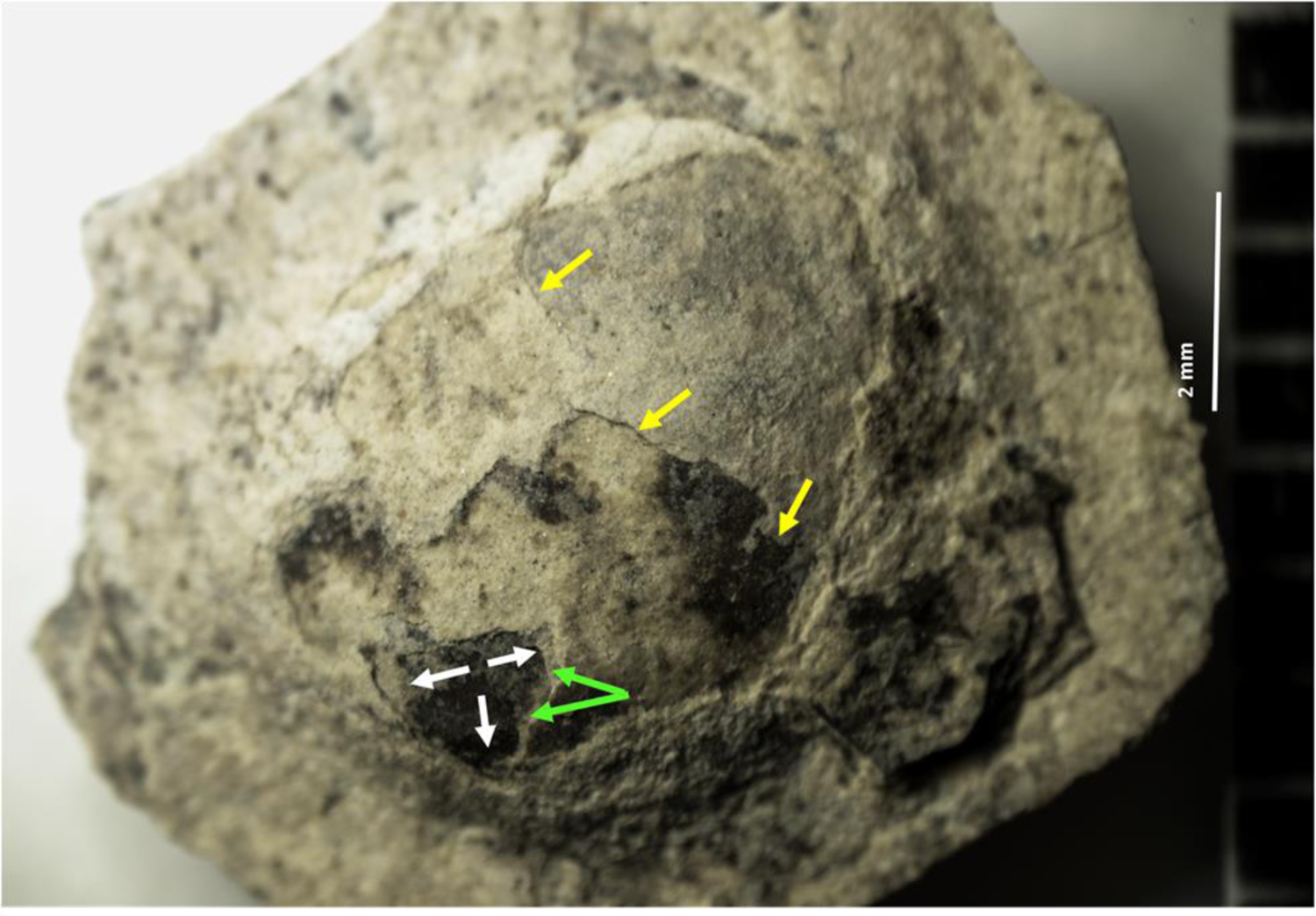
Areas of the convex surface of the NUS_N9_2016 E-2 egg where the two calcified layers of its eggshell were detected. The yellow areas indicate the limit of the top layer of the eggshell. Within the dark area delimited by the white arrows, which corresponds to a depression on the surface of the egg, the inner calcified layer of the eggshell is indicated by the green arrows (see also Figures S46-S50).

**Figure S46.**
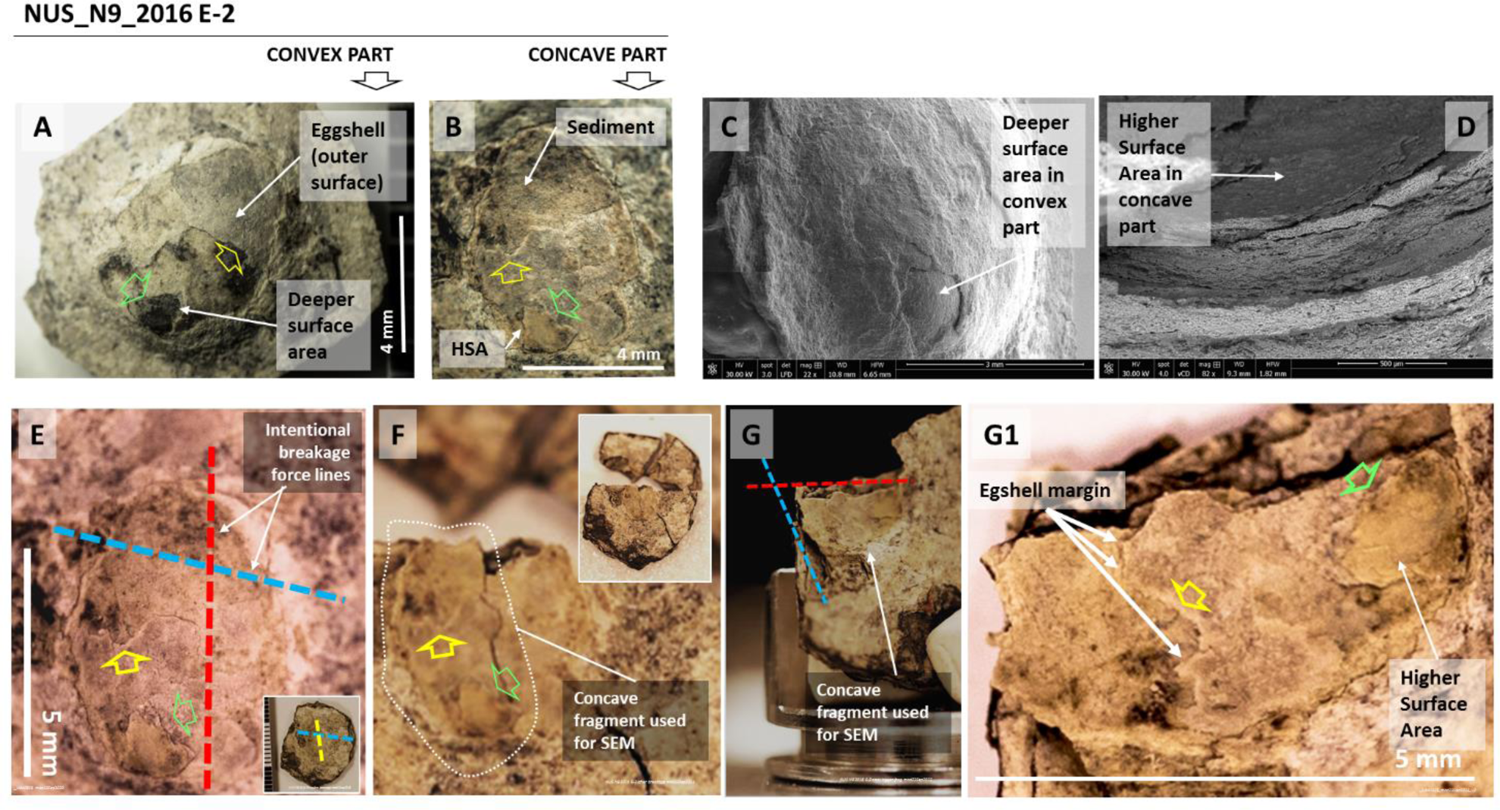
Approach to the study of a deeper area on the convex surface of the NUS_N9_2016 E-2 egg. A deeper area on the convex surface of the egg, indicated by the green open arrow (**A**), matches a higher surface area (HSA), also indicated by an open green arrow, on the concave surface of the fragment that was detached from the 3D-preserved egg (**B)**. The edge of the outer layer of the eggshell is indicated by an open yellow arrow in A and B. The deeper area on the convex surface of the egg under SEM is indicated in **C**. The higher surface area on the concave surface of the detached fragment under SEM is indicated in **D**. Two fractures were intentionally made on the concave surface of the detached fragment from NUS_N9_2016 E-2 along the red and blue dashed lines shown in **E**. The edge of the outer calcified layer of the eggshell is indicated by an open yellow arrow. **F**: Surface of the concave fragment after its fracture along the red dashed line shown in E. **G**: Surface of the concave fragment after its fracture along the red and blue dashed lines shown in E, already mounted on a SEM stub. **G1**: magnified view of the concave surface of the detached fragment already fractured along the red and blue dashed lines shown in E, indicating the edge of the outer calcified eggshell layer (yellow arrow) and the edge of the inner calcified layer (green arrow) on the higher surface area of this fragment.

**Figure S47.**
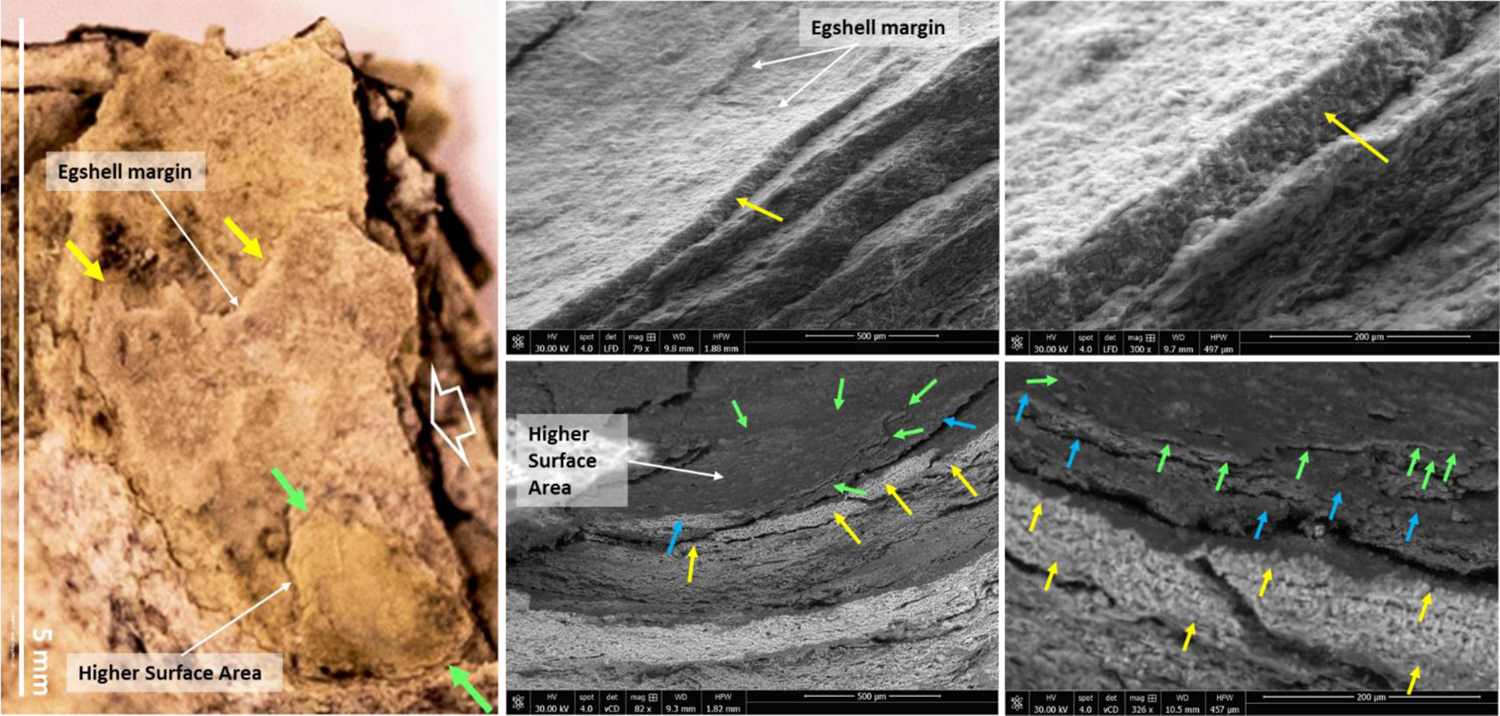
Different eggshell layers of the NUS_N9_2016 E-2 egg. Fractured concave surface of the detached fragment (right), indicating the edge of the outer calcified eggshell layer (yellow arrow) and the edge of the inner calcified layer (green arrow) on the higher surface area of this fragment. The outer calcified layer is observable under SEM on top as indicated by the yellow arrow and the inner calcified layer is observable under SEM on the bottom as indicated by the green arrow. The middle layer is indicated by blue arrows (see also Figure S48).

**Figure S48.**
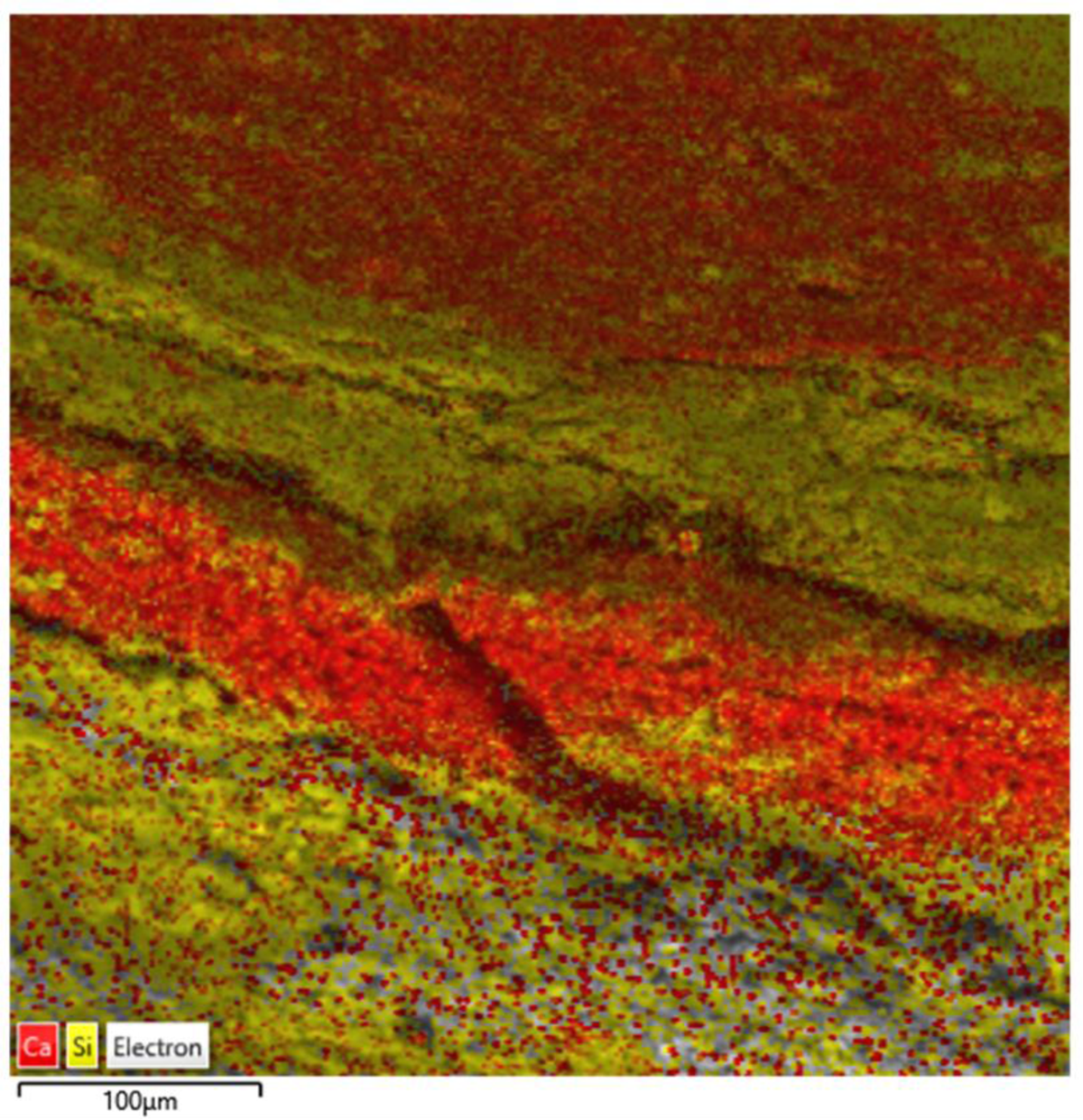
Two different calcified layers in the ornithuromorph NUS_N9_2016 E-2 egg. Red indicates calcium rich layers and yellow indicates silica rich areas. The red layer on top is the inner calcified layer, the red layer across the center of the image is the outer calcified layer of the eggshell. Between the two calcified layers resides the middle non-calcified layer.

**Figure S49.**
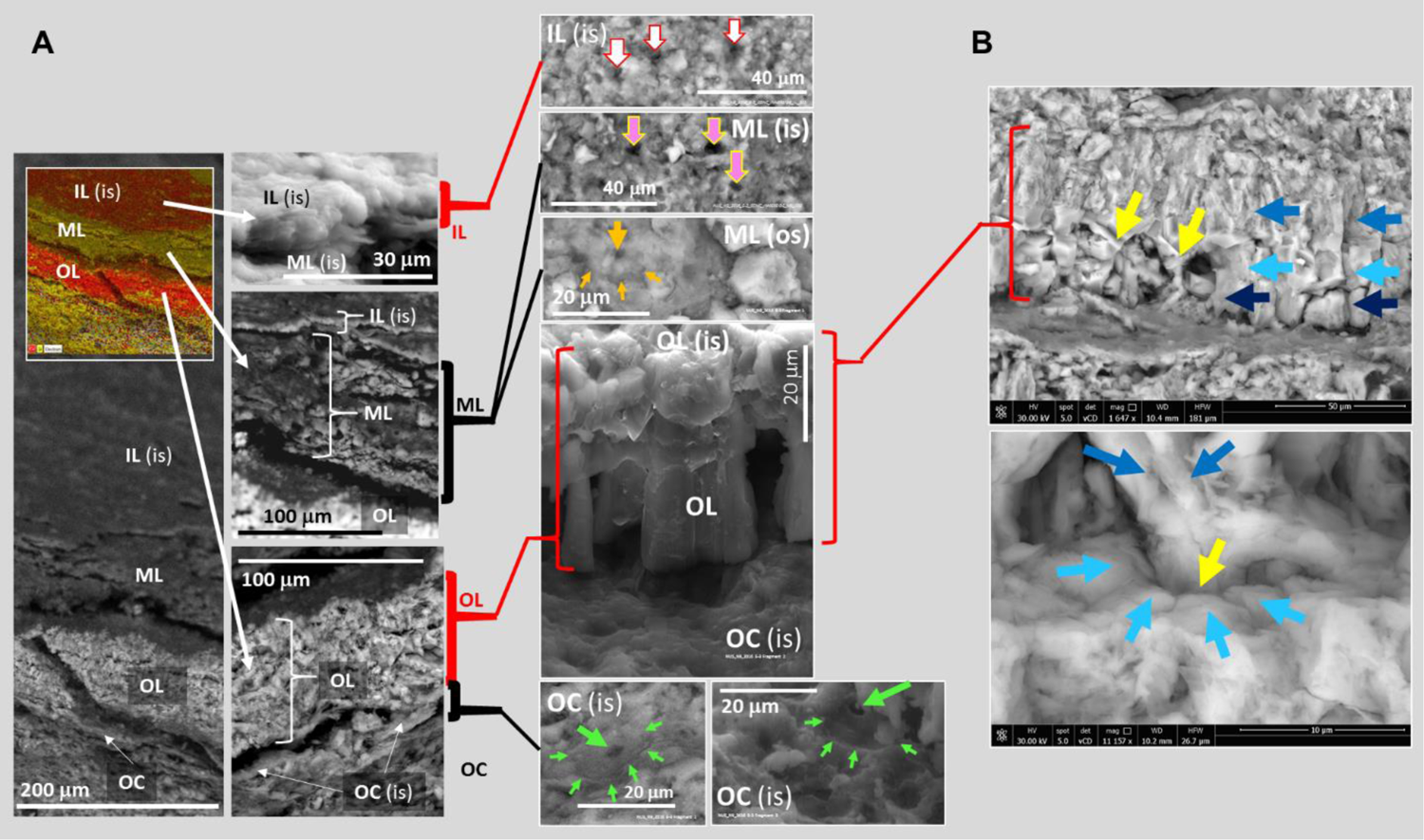
Different layers of the ornithuromorph eggshell. **A:** an inner calcified layer (IL), a middle non-calcified layer (ML), an outer calcified layer (ML) and an outer non-calcified cuticle (OC) are indicated in a SEM image and in a EDS map (inset) of the inner surface (is) of the concave fragment detached from the NUS_N9_2016 E-2 egg. Red indicates calcium rich layers and yellow indicates silica rich areas in the EDS map. Amplified SEM images of each layer are provided. Perforations are indicated by arrows of different colors. Some perforations seem to be encircled by a wider circular margin, as indicated in the outer surface (os) of the middle layer (ML) and in the inner surface of the outer cuticle (OC). **B:** detail of the outer calcified layer shown a prismatic outer layer, which appears to derive from an inner nucleation point, and how this layer appears to be fractured into two or three sublayers represented by arrows in different tones of blue (work in progress).

**Figure S50.**
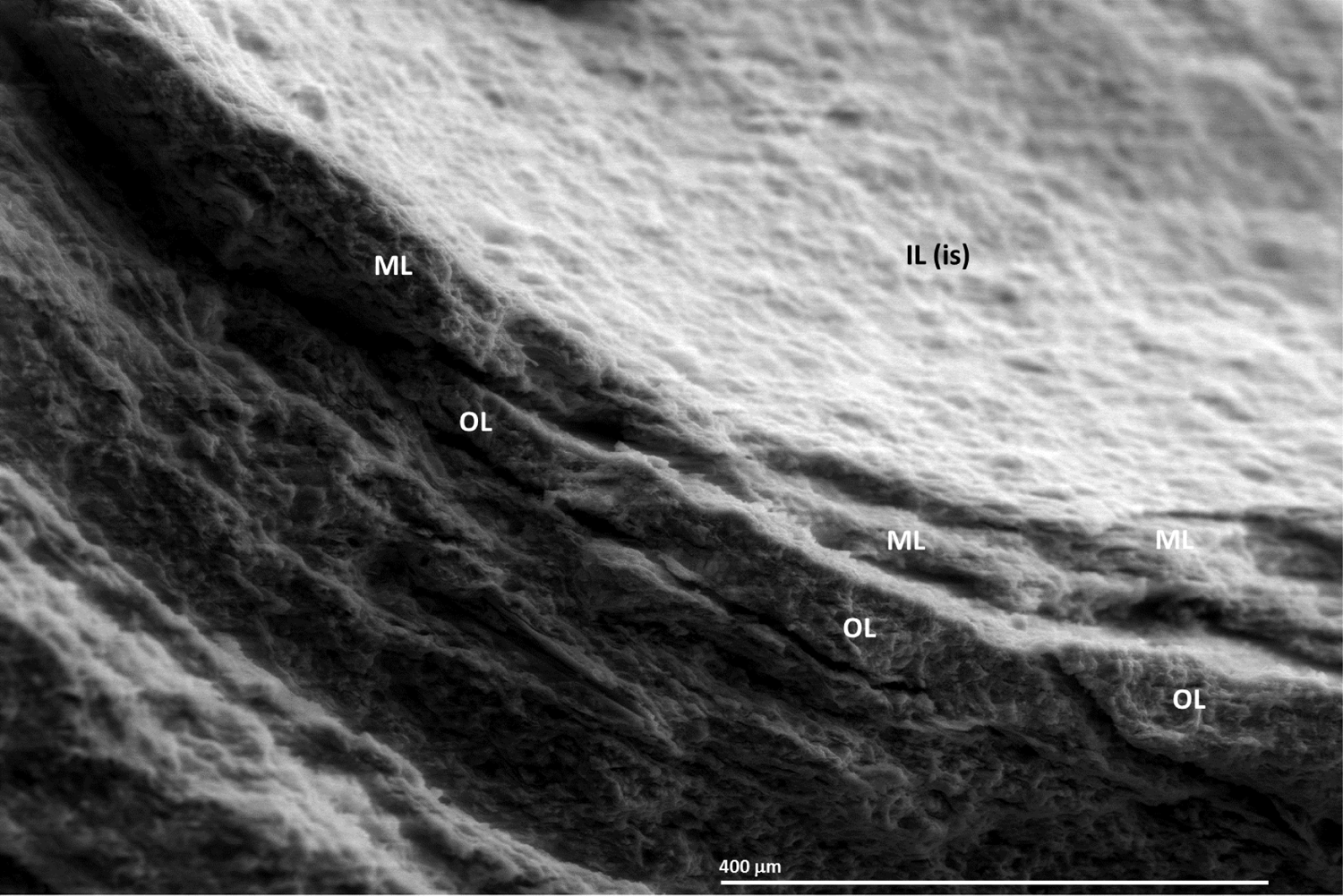
Different layers of the ornithuromorph eggshell. Inner surface (is) of the inner calcified layer (IL), middle non-calcified layer (ML), and outer calcified layer (ML) of the eggshell observable in the concave fragment detached from the NUS_N9_2016 E-2 egg.

**Figure S51.**
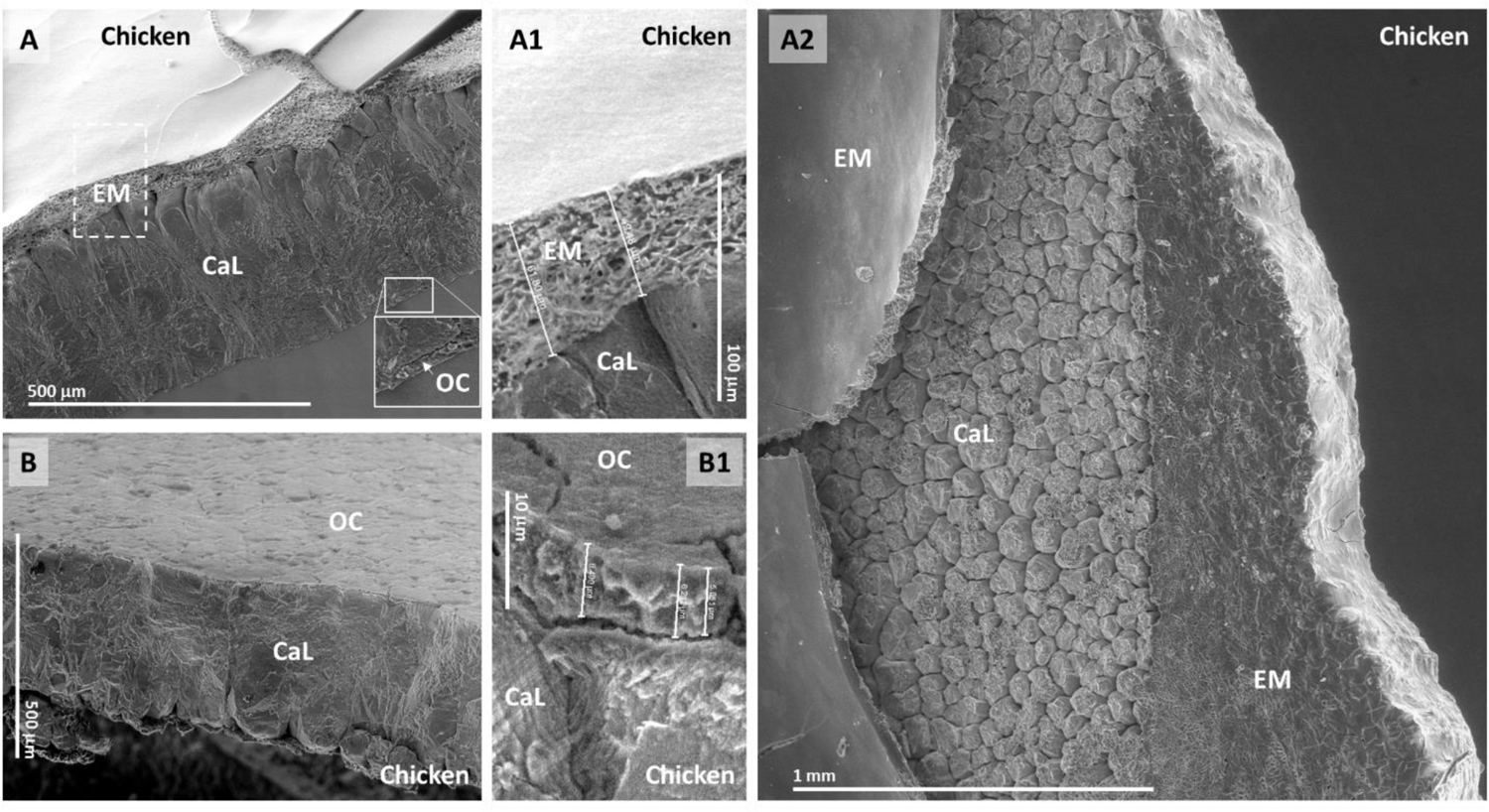
Chicken eggshell. Images of a chicken eggshell exposed to air for a few weeks under SEM are provided. **A**: SEM view of the inner side of the chicken eggshell with identification of the eggshell membranes (EM), calcified layer of the eggshell (CaL) and outer cuticle (OC), magnified in **A1** to put the eggshell membranes into evidence. Another area of the inner surface of the same eggshell is shown in **A2** to demonstrate how the inner eggshell layer represented by the eggshell membranes is not durable, resulting in exposure of the calcified layer and its calcite prisms. **B**: Magnified view of the outer cuticle that is external to the calcified layer, further magnified in **B1**.

**Figure S52.**
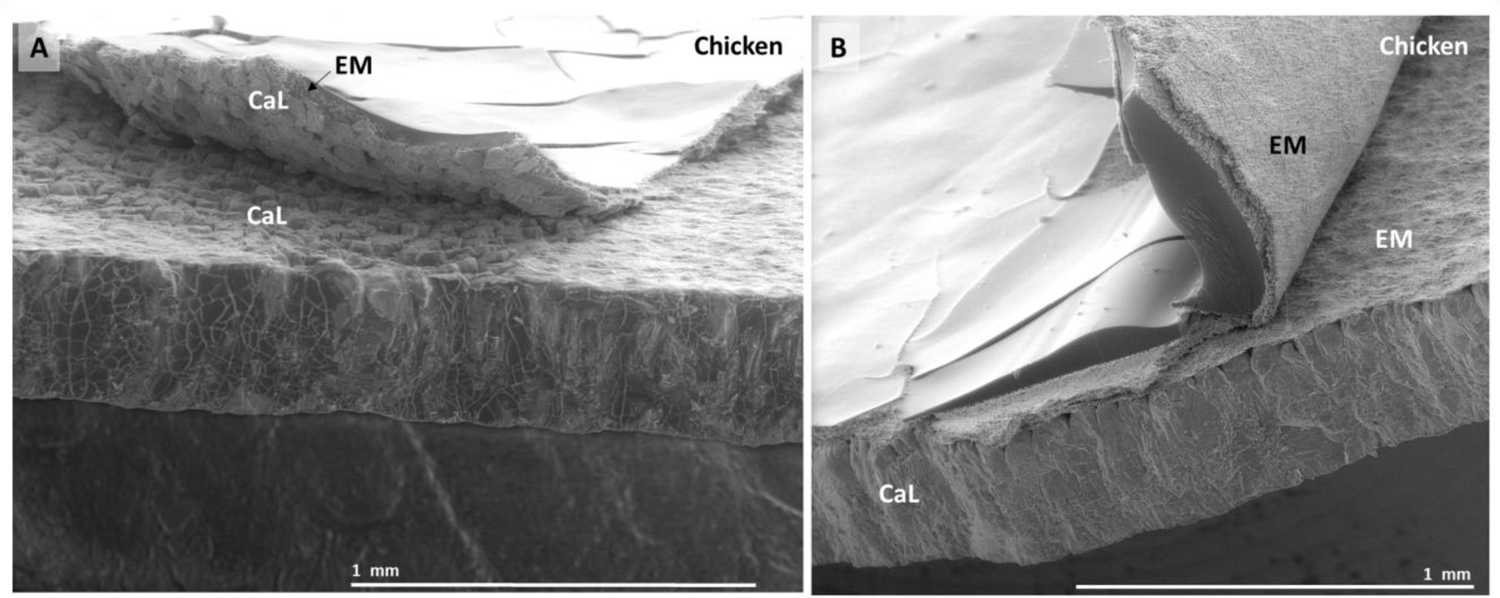
Detachment of the eggshell membranes of the chicken eggshell from natural decay. The eggshell membranes are shown to detach from the calcitic layer, sometimes to the point of causing fractures in the calcitic layer (**A**), other times separating part of the eggshell membranes from each other (**B**).

**Figure S53.**
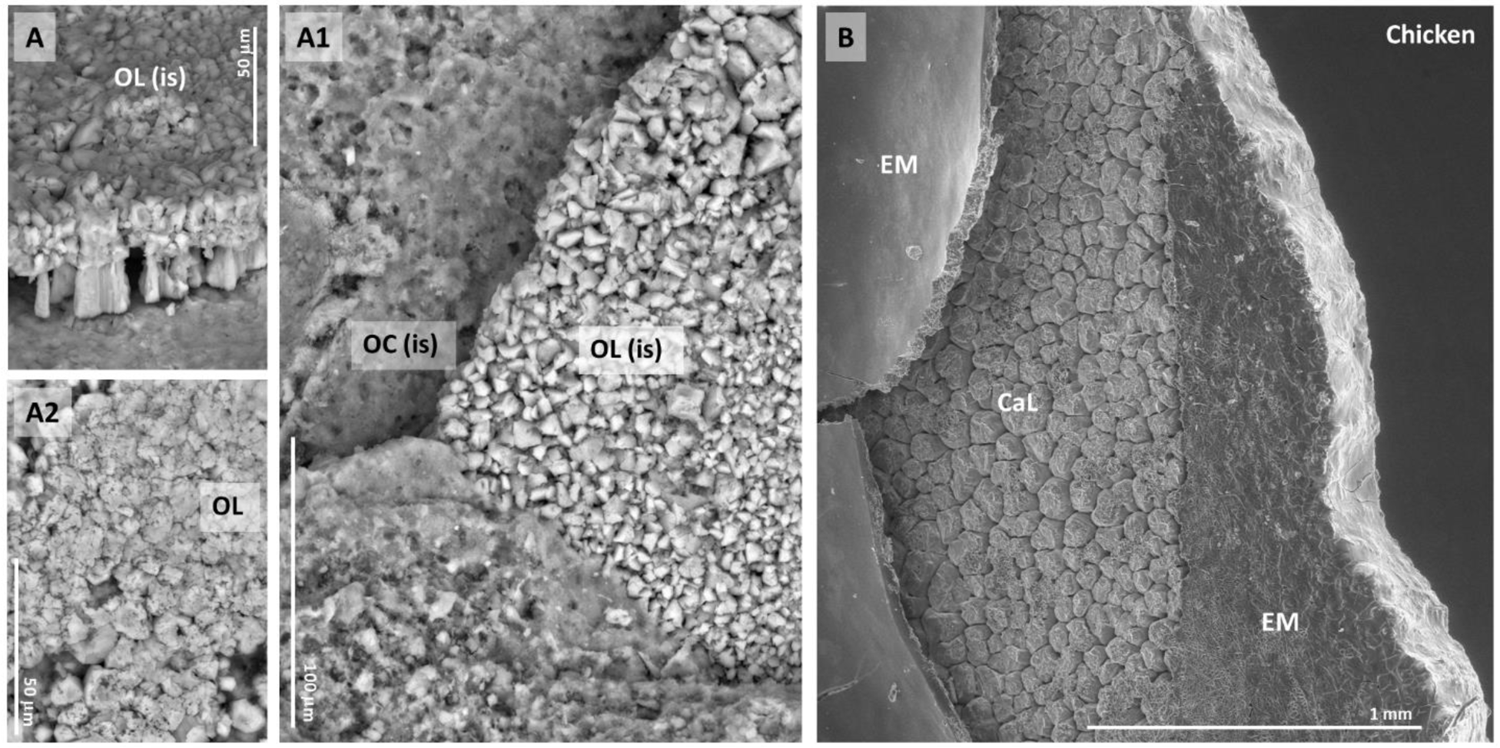
Appearance of the fossilized ornithuromorph eggshell in comparison with naturally decayed chicken eggshell. The calcitic outer layer (OL) of the fossilized ornithuromorph eggshell is shown in **A**, **A1** and **A2**, as well as the outer cuticle (OC), both from their inner side (is) in A and A1. Naturally decayed chicken eggshell is shown in **B**, with separation of the eggshell membranes (EM) from the calcitic layer (CaL). Note the similar appearance of the calcite prisms.

**Figure S54.**
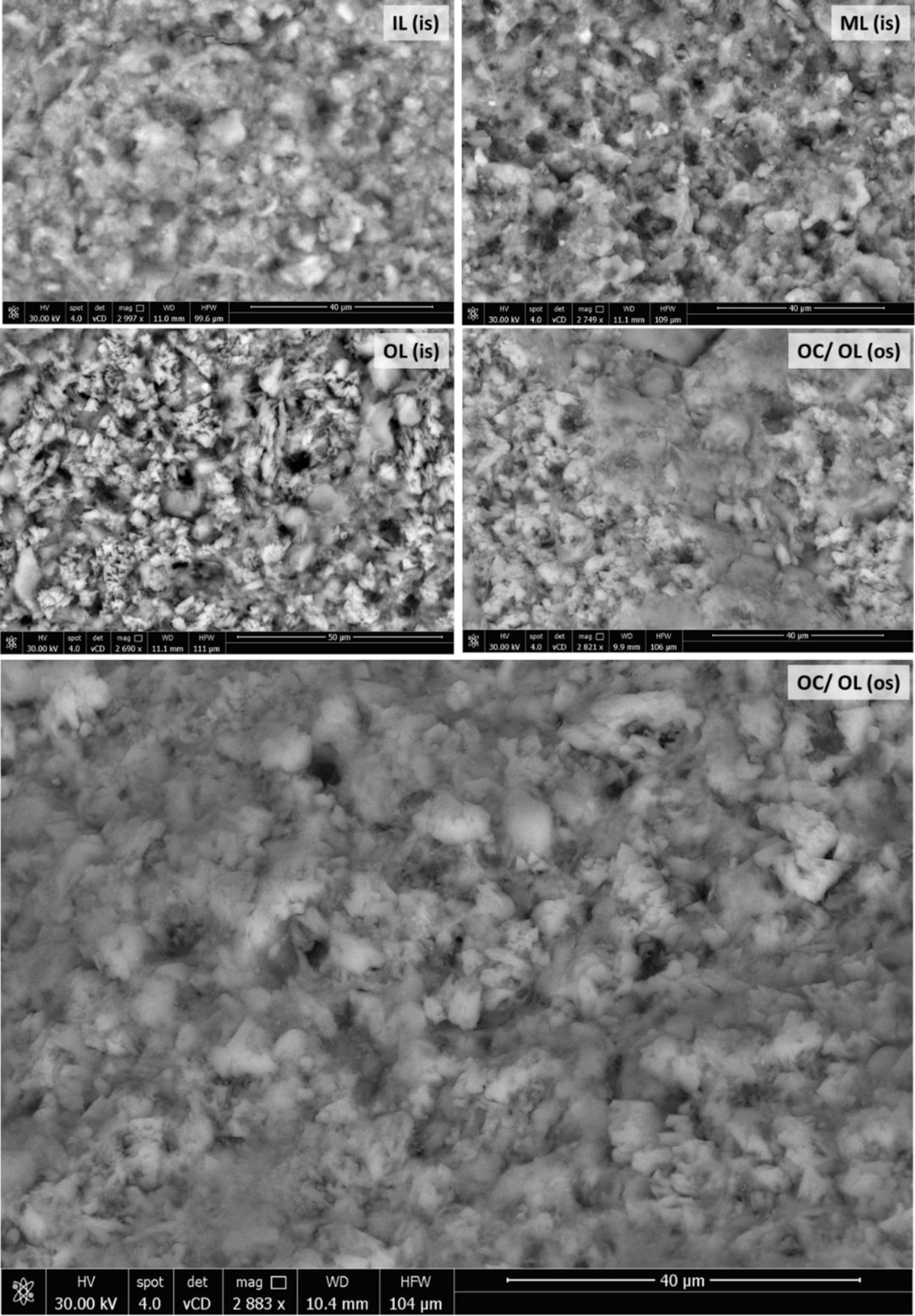
Perforations in the ornithuromorph eggshell under SEM. Perforations observable under SEM are shown in the inner surface (is) of the inner layer (IL), middle layer (ML) and outer layer (OL), and on the outer surface (os) of the outer layer and outer cuticle (OC /OL). These perforations are of similar diameter, below 4 m, and relatively regularly spaced, at 10 – 40 m intervals. It remains to be determined if these are artifacts.

**Figure S55.**
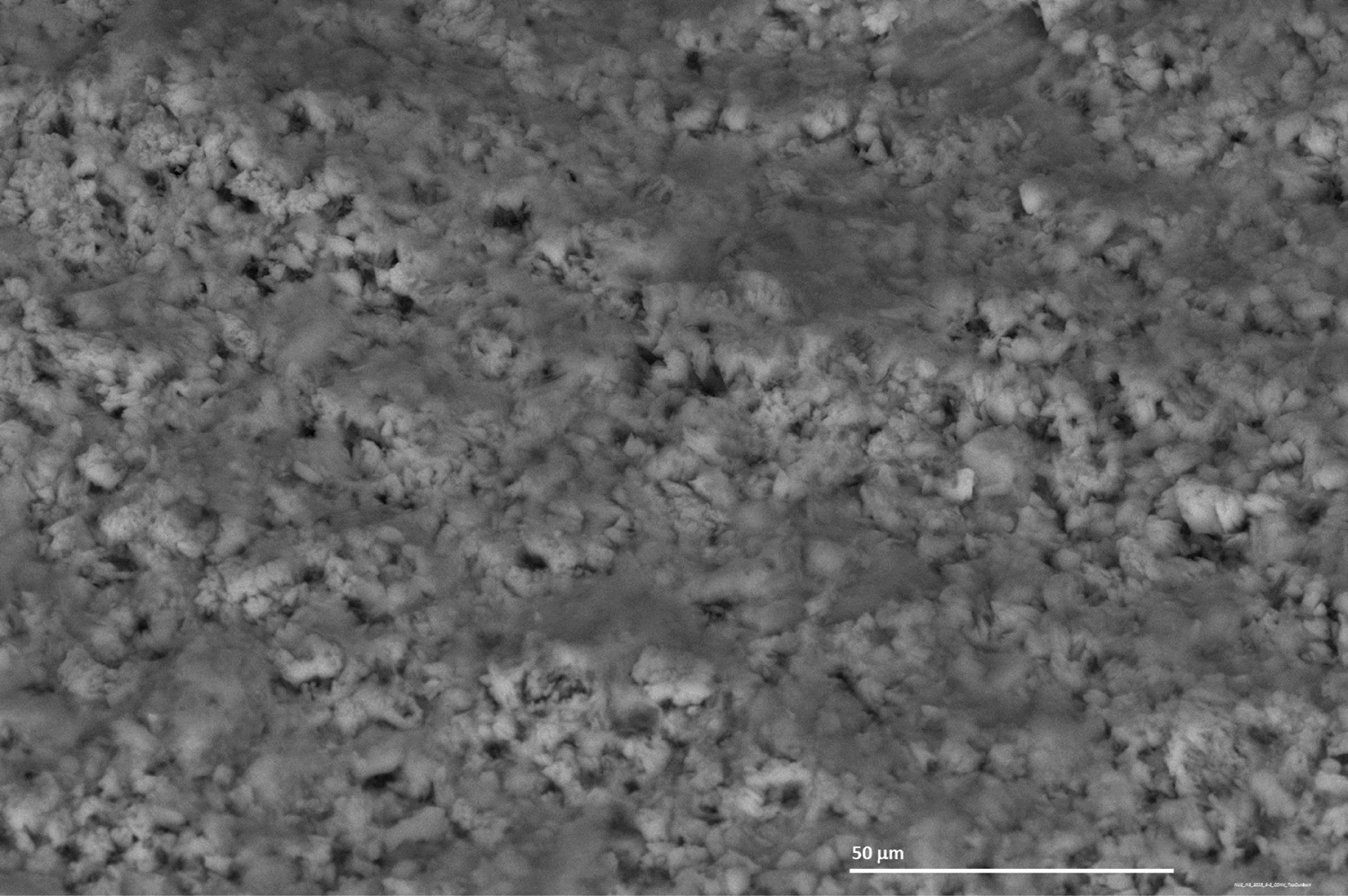
Appearance of the outer surface of the external calcitic layer of the ornithuromorph eggshell under SEM.

**Figure S56.**
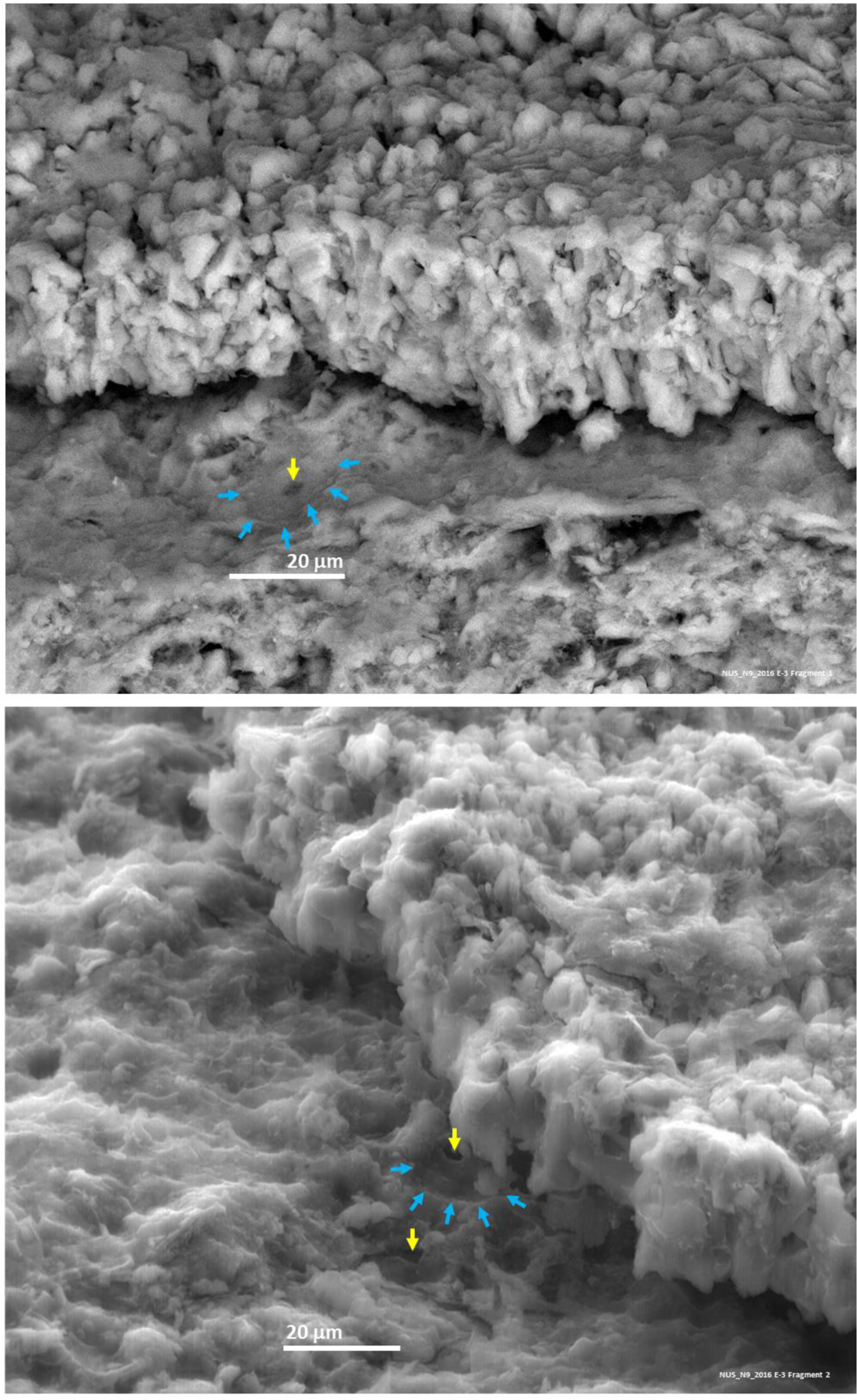
Perforations in the inner surface of the outer cuticle of the ornithuromorph eggshell. The perforation is indicated by a yellow arrow and a ringed or donut-shaped periphery is indicated by blue arrows. The possibility of an arrangement of the prismatic layer into rings is under investigation.

**Figure S57.**
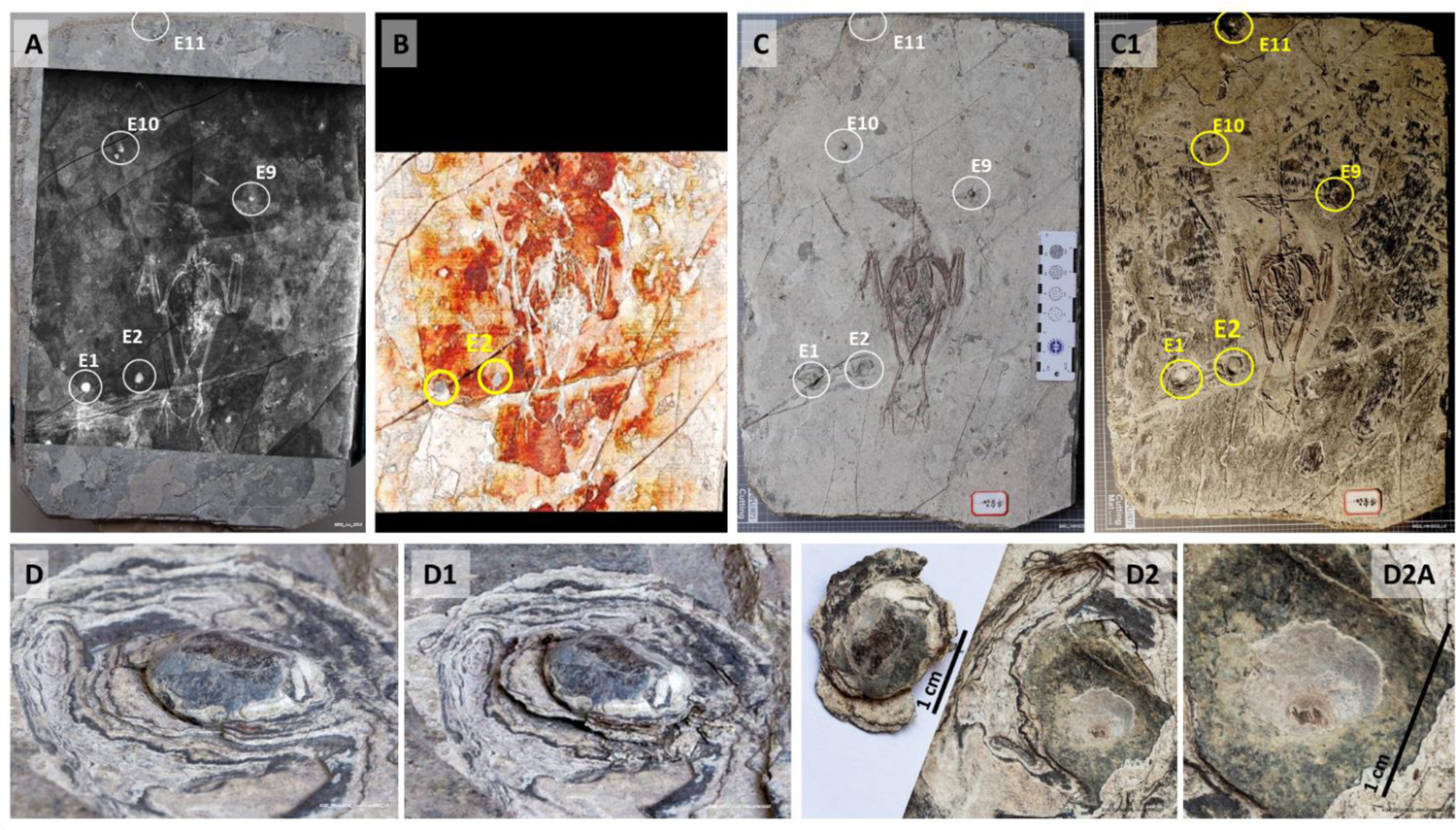
Additional fossil preparation of EHG_J1_2016 exposed a heterogeneous fossilized environment compatible with that of a fossilized nest. Guided by X-rays (**A**) and CT scan data (**B**), additional fossil preparation was undertaken to the level of detachment of 5 egg-candidate structures identified via surface irregularities and radiologic means (E1, E2, E3, E9 and E10; A, **C** and **C1**), exposing a heterogeneous fossilized environment (C1) that explains the 3D CT scan image captured in B by showing material of different density in the immediate surrounds of the bird skeleton. EHG_J1_2016 E2 is shown before detachment (**D**), at the stage of detachment (**D1**) and after detachment from the fossil, in addition to the area of the fossil it detached from (**D2** and **D2A**).

**Figure S58.**
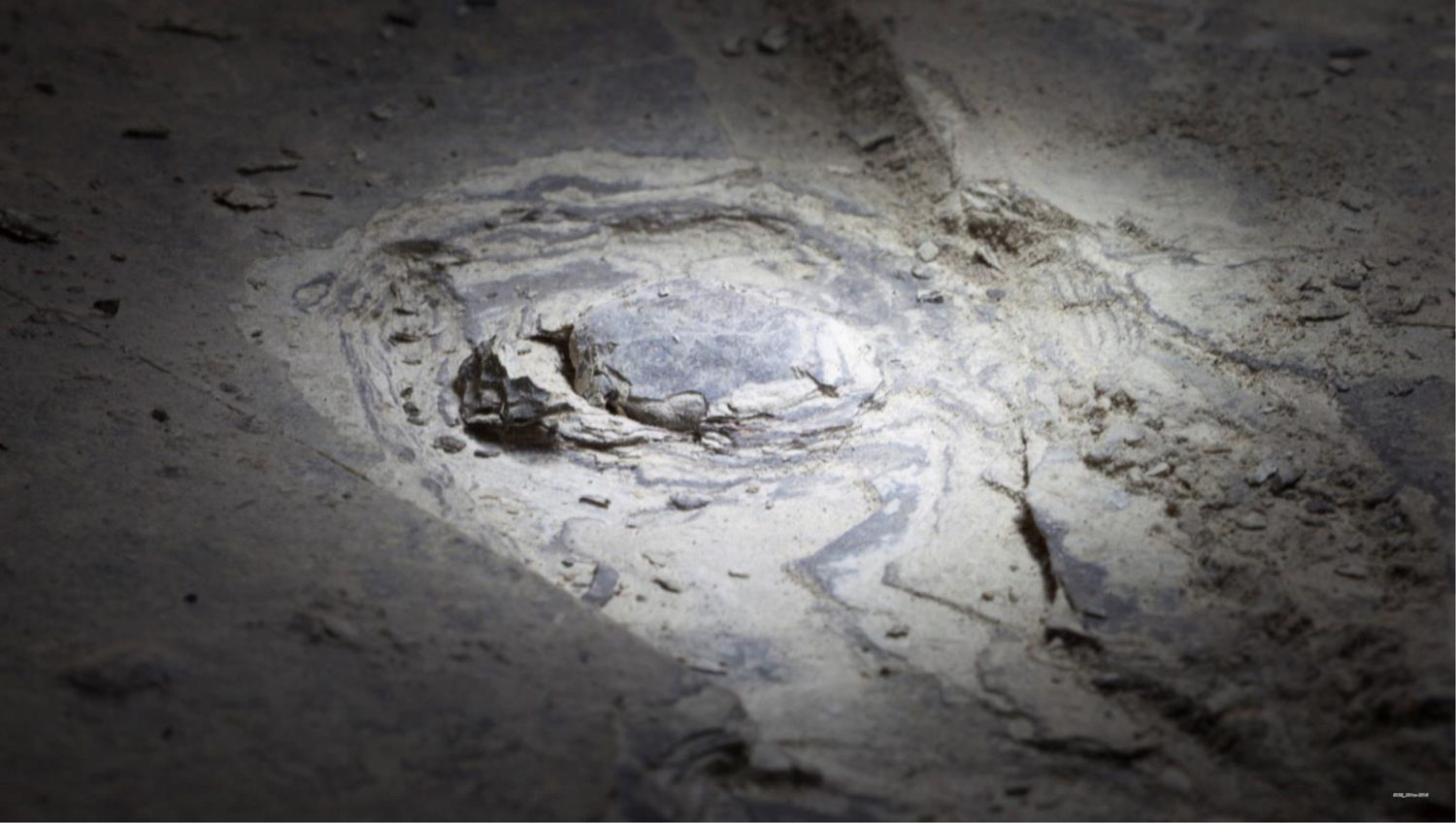
Step of the extraction of the EHG_J1_2016 E1 egg from the fossil slab of the EHG_J1_2016 ornithuromorph specimen as seen under stereomicroscopy.

**Figure S59.**
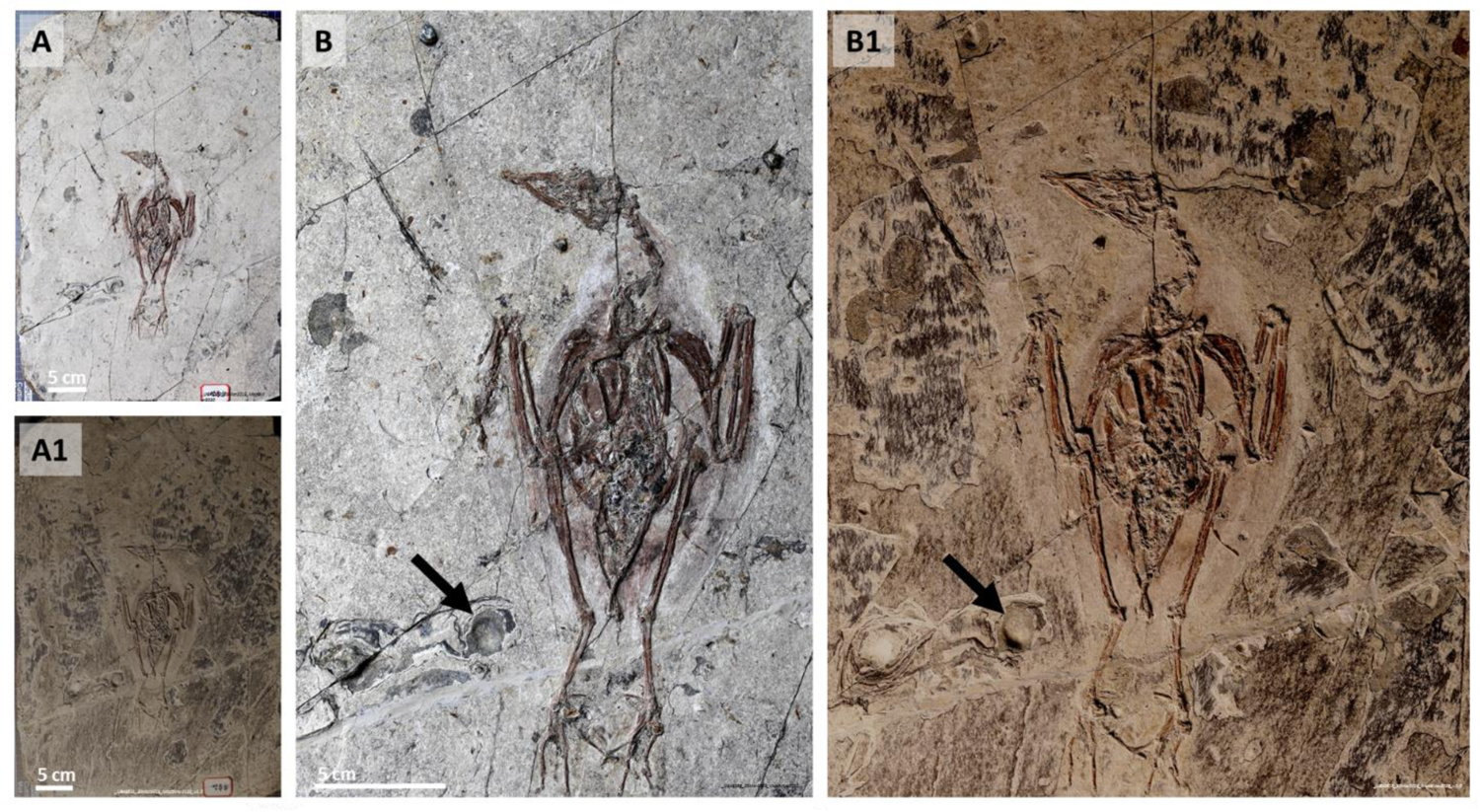
Fossil of specimen EHG_J1_2016 before and after additional preparation. Sediment was removed from the fossil (**A**) until the slab surface level at which detachment of putative eggs identified by X-rays and CT scans occurred (**A1**). The magnified view before (**B**) and after (**B1**) preparation shows the concavity left behind by the removal of EHG_J1_2016 E2 (arrow in B and B1).

**Figure S60.**
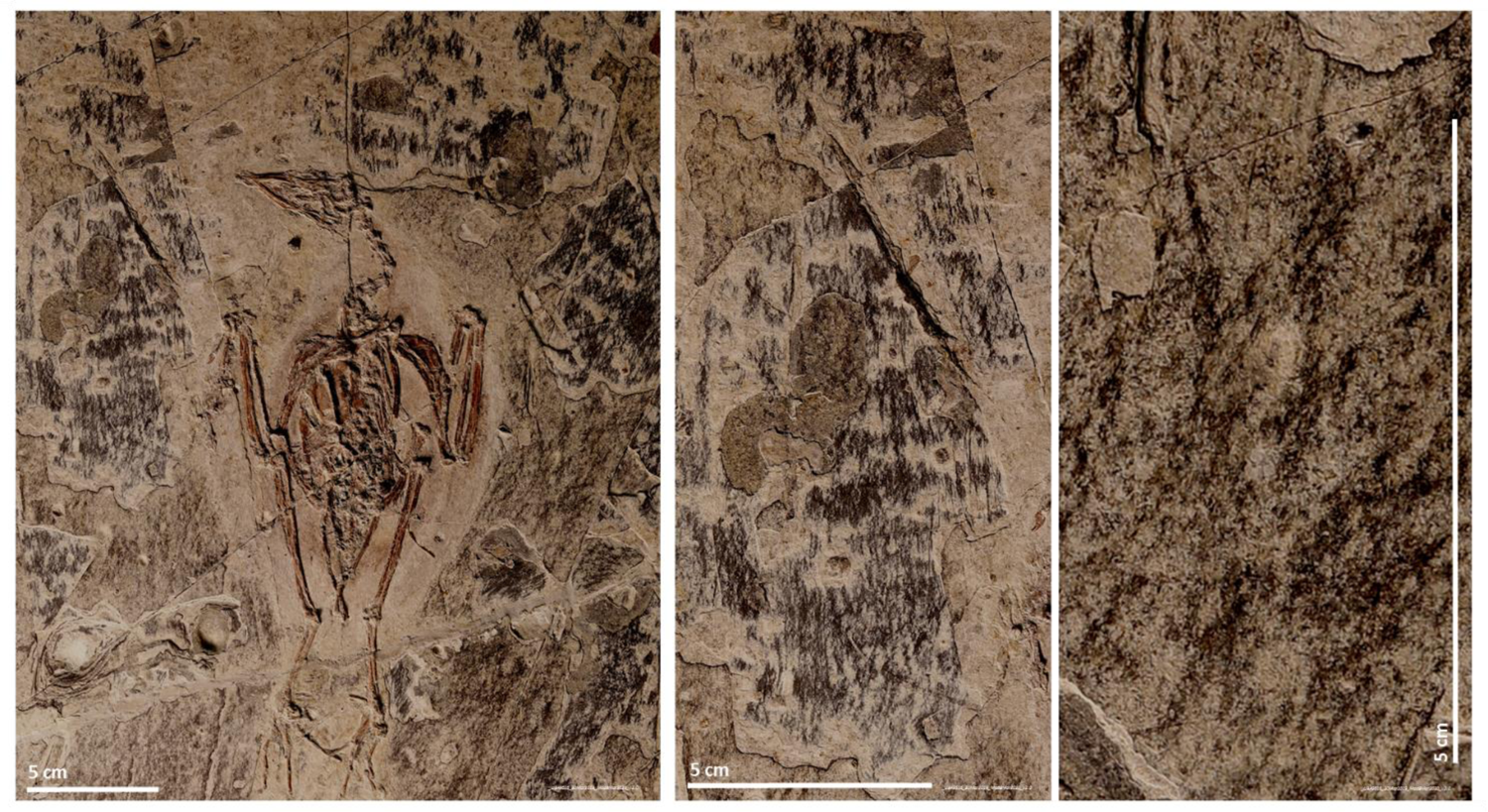
Details of the slab surface of EHG_J1_2016 specimen’s fossil after preparation. The removal of sediment down to the level at which eggs detached from the sediment resulted in the exposure of a heterogeneous picture compatible with that of fossilized vegetation.

**Figure S61.**
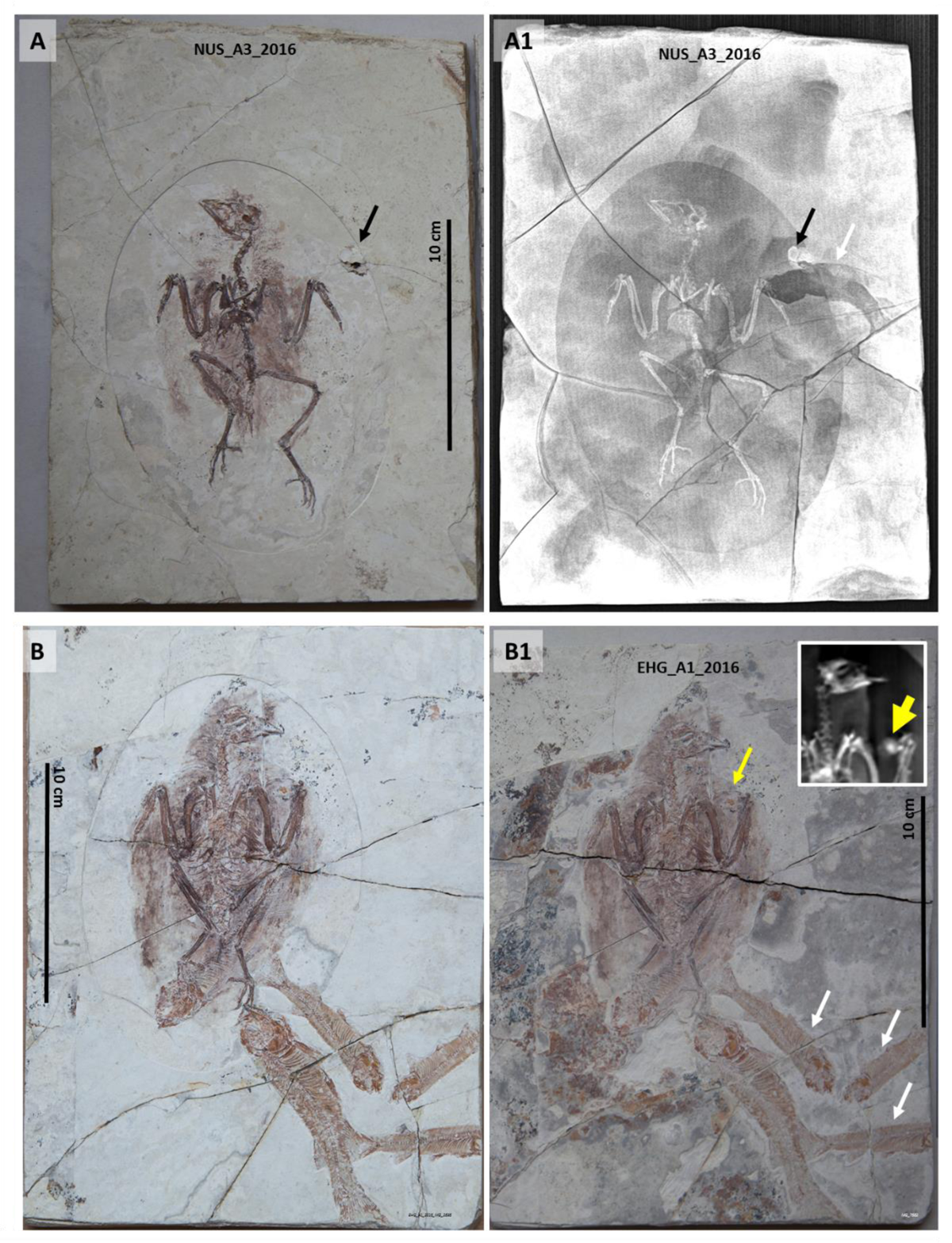
Two fossilized unspecified enantiornithes associated with fish skeletons and eggs. The fossil of NUS_A3_2016 is shown in **A** before additional preparation and under CT scanning (**A1**). The fossil of EHG_A1_2016 is shown before (**B**) and after (**B1**) additional preparation. White arrows point at fish skeletons in both fossils. Black arrows point at a small cluster of eggs associated with NUS_A3_2016 (see also Figure S62). Yellow arrow points at an egg associated with EHG_A1_2016, shown under CT scanning in the B1 inset.

**Figure S62.**
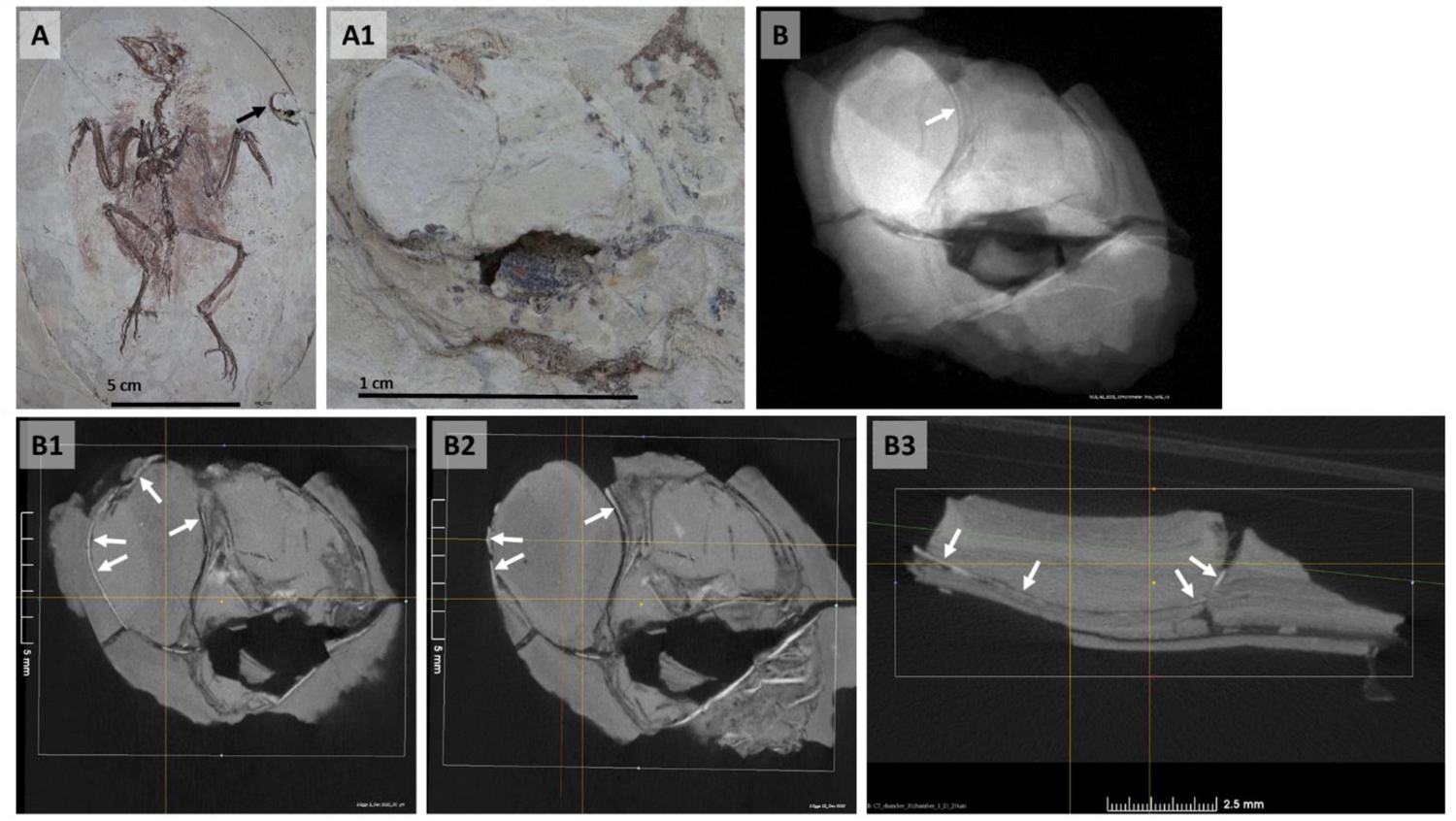
Egg cluster associated with the NUS_A3_2016 fossil. The unspecified enantiornithe is shown under macrophotography (**A**). The egg cluster associated with it (arrow) is amplified (**A1**) and shown under a 10 μm resolution X-ray (**B**) and 2D CT scan sections in coronal (**B1** and **B2**) and sagittal (**B3**) views. White arrows point at the eggshell.

**Figure S63.**
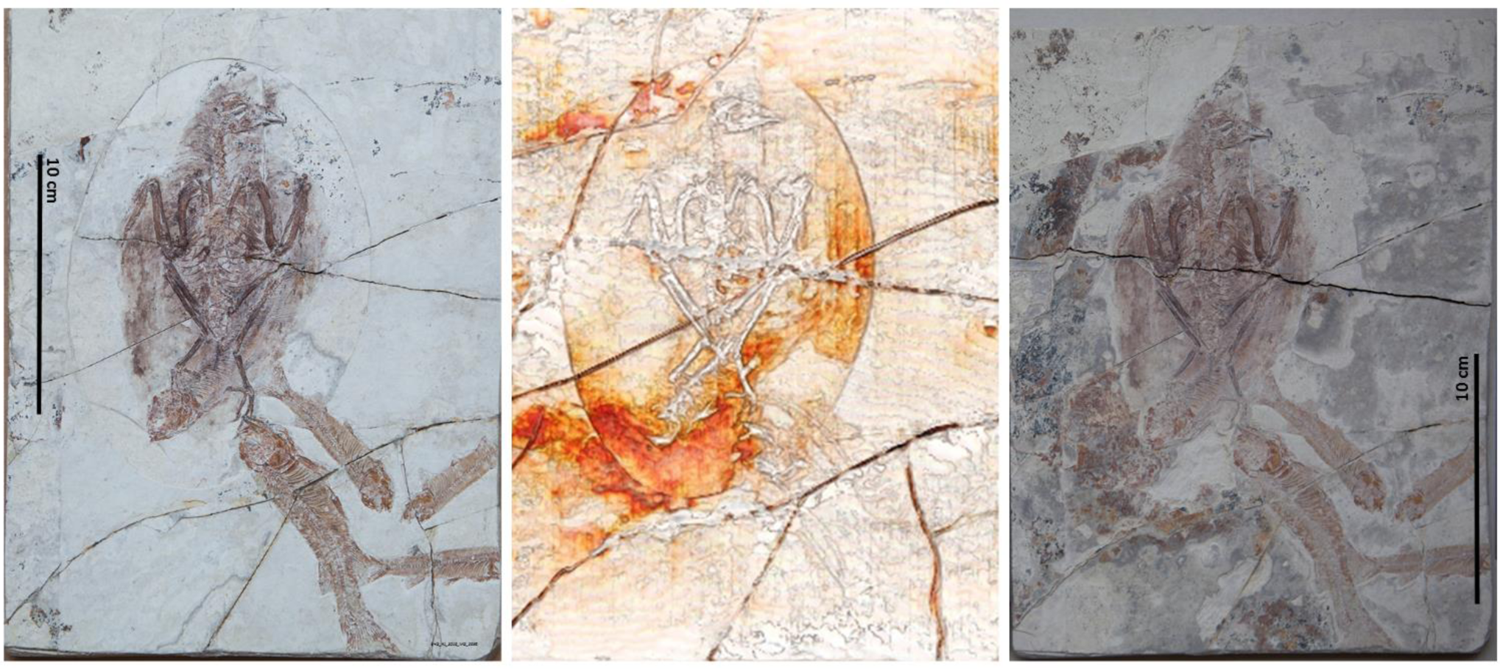
Unspecified enantiornithe EHG_A1_2016 association with fish skeletons. The fossil is shown before (left), under CT scanning (center) and after additional preparation (right). A CT scan image was used to guide fossil preparation and expose fossilized vegetation.

**Figure S64.**
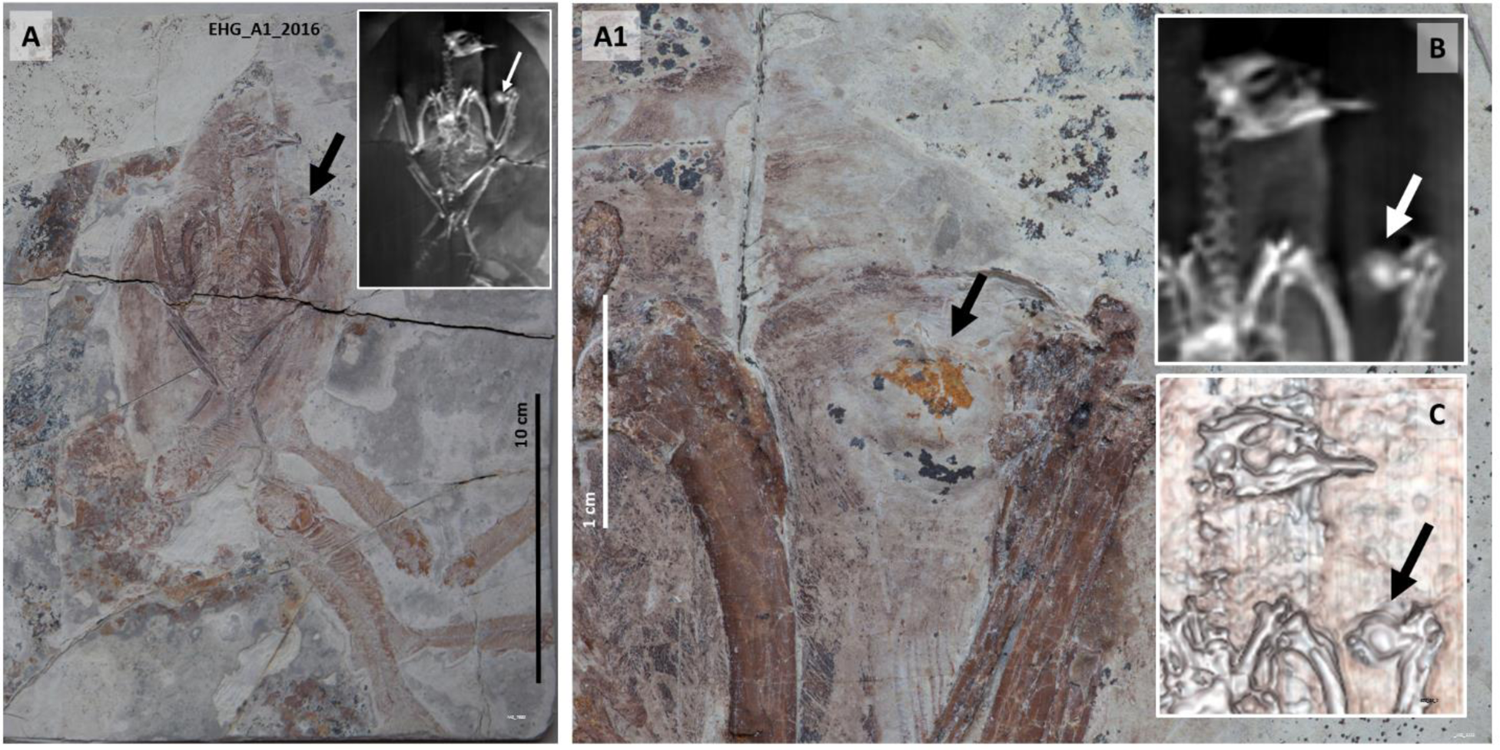
Unspecified enantiornithe EHG_A1_2016 in dorsal view associated with an egg and fish skeletons atop fossilized vegetation. An X-ray image is shown in the inset in **A**. The location of the egg (yellow arrows) in the inner space anterior to the folded right wing is shown under macrophotography in **A1**, a coronal CT scan view (**B**) and a 3D CT scan image (**C**).

**Figure S65.**
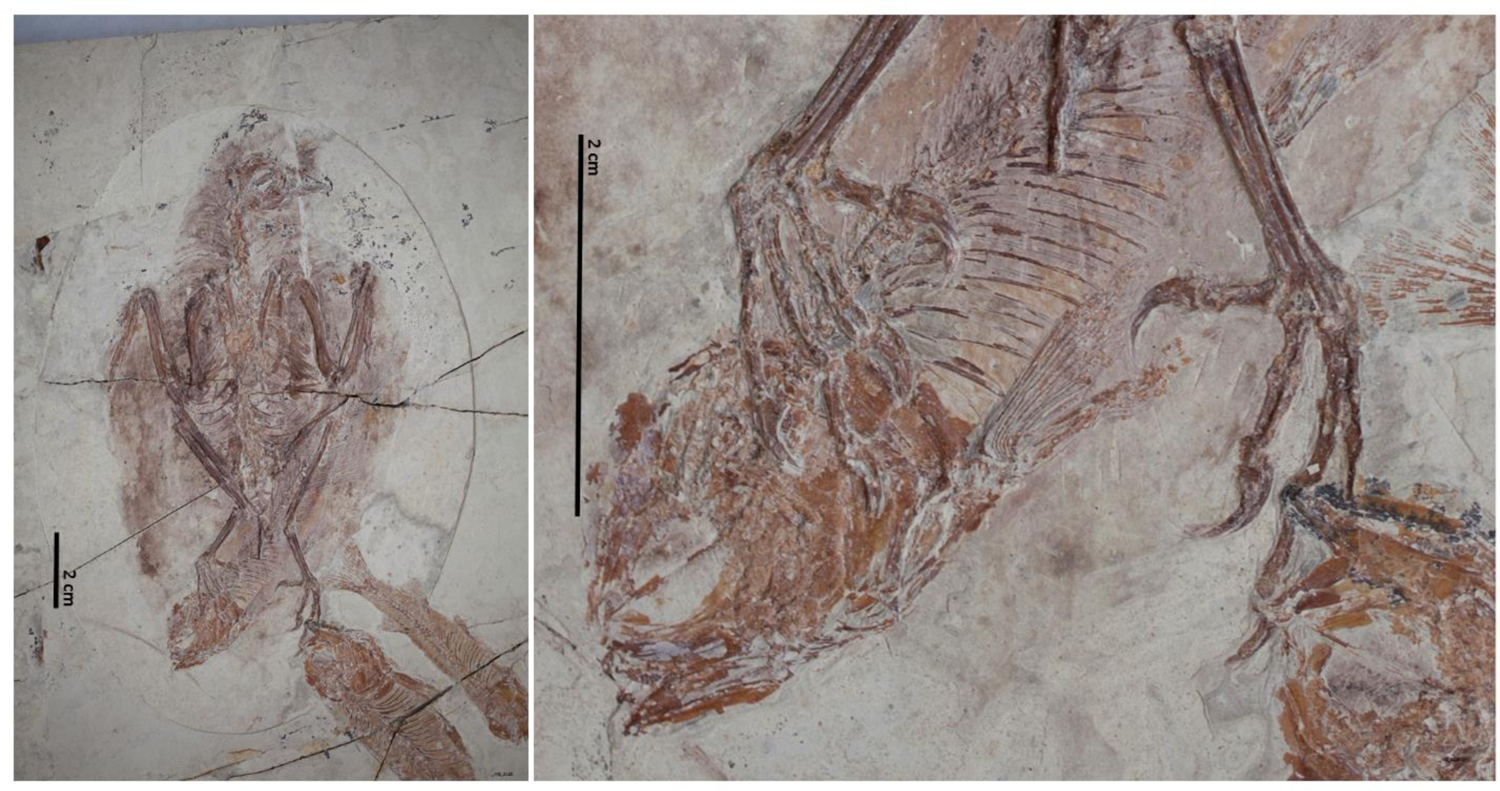
Unspecified enantiornithe EHG_A1_2016 has its left foot atop one fossilized fish skeleton and its right foot below another. The bird is in dorsal view. Its fossilized pose is compatible with a nesting posture.

**Figure S66.**
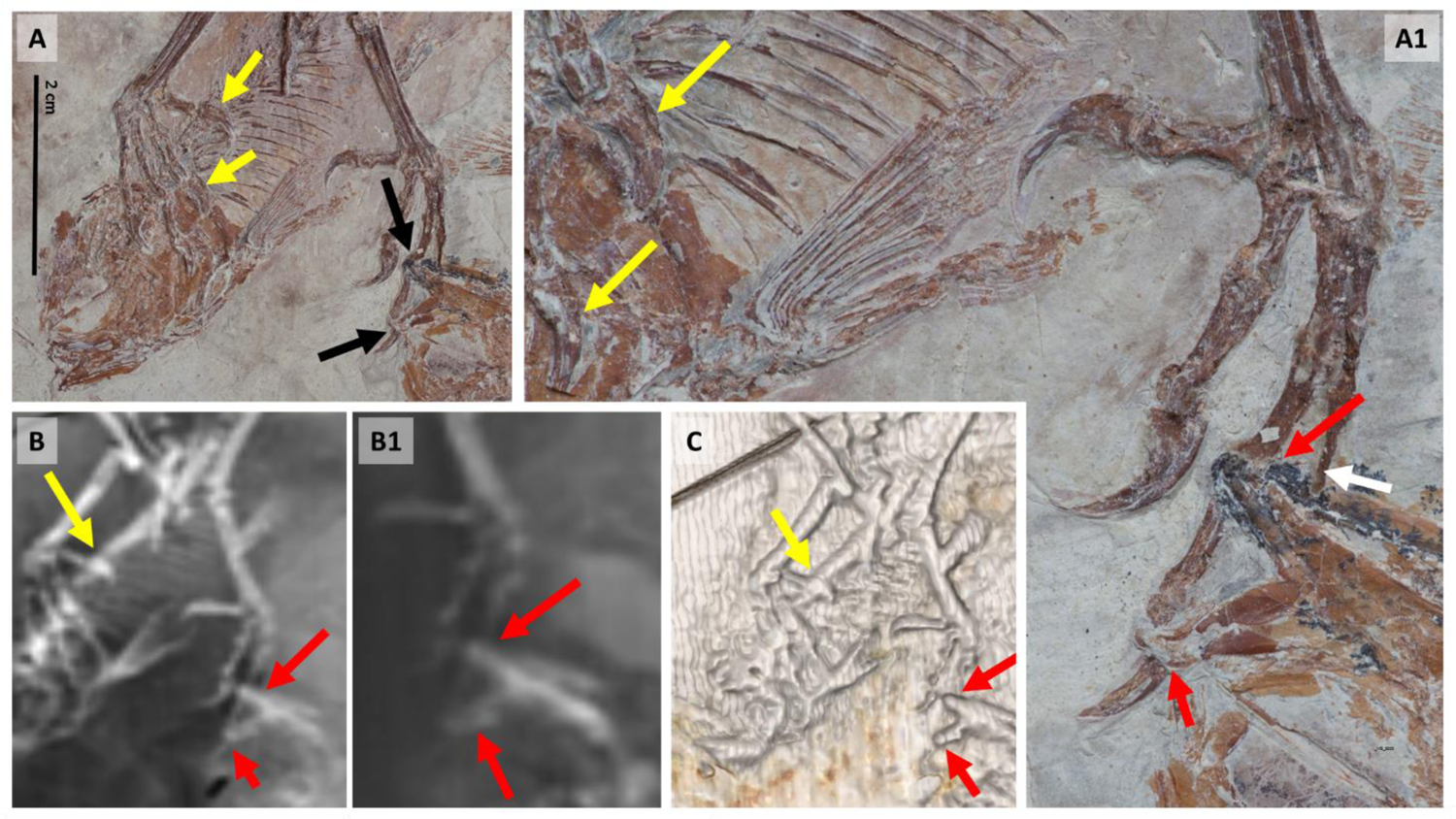
Unspecified enantiornithe EHG_A1_2016 has its left foot atop one fossilized fish skeleton and part of its right foot below another. The bird’s left foot fingers are dorsal to a fish skeleton (yellow arrows) whereas the right foot’s third finger is ventral to another (red arrows). It is unclear if a different right foot finger (white arrow) is dorsal to the same fish skeleton. 2D (bottom left and bottom center) and 3D (bottom right) CT scan images are shown.

**Figure S67.**
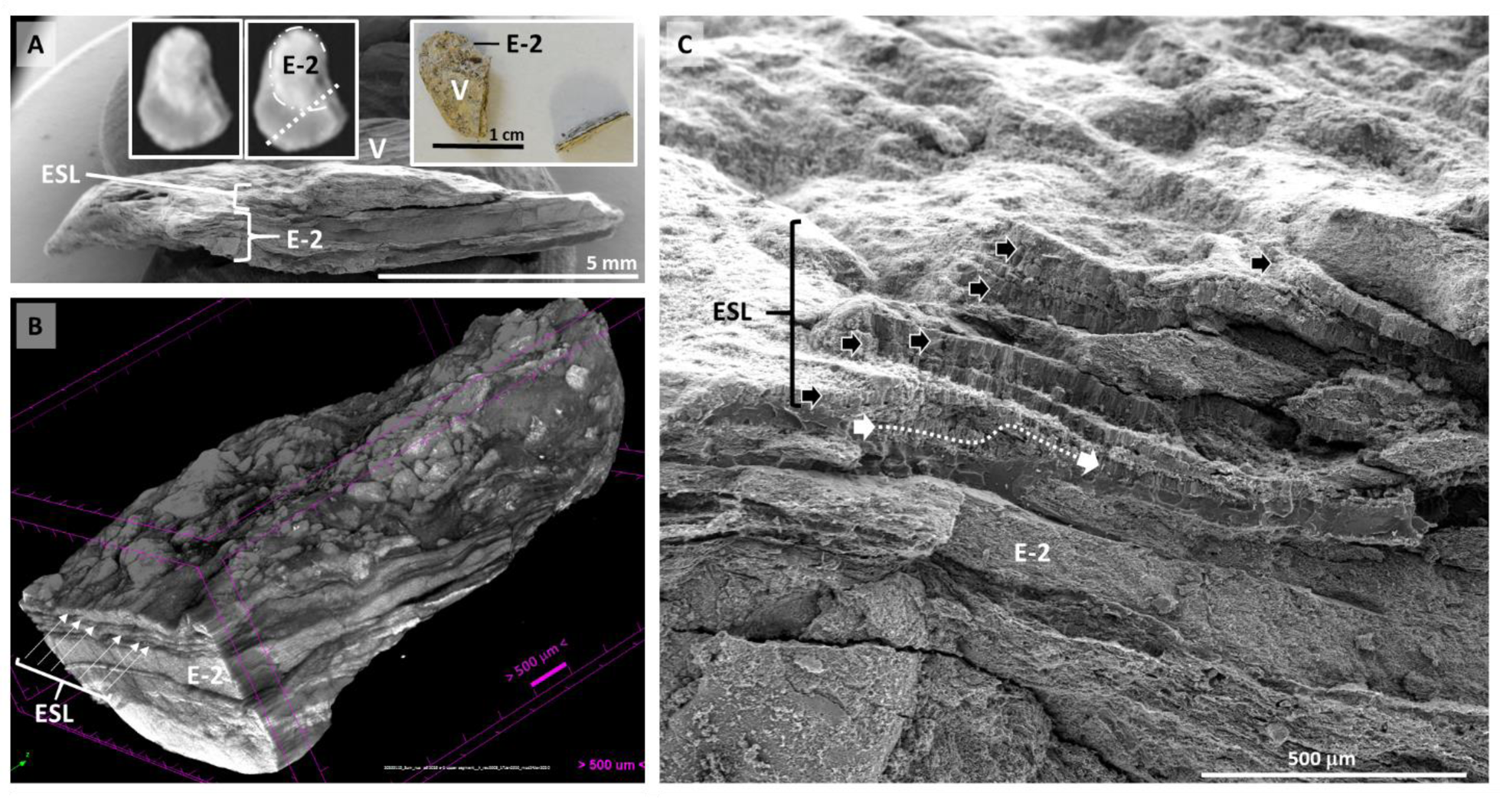
Layers of eggshell remnants associated with an egg extracted from the NUS_A6_2016 fossil. **A**: Radial surface of the fractured NUS_A6_2016 E-2 egg. NUS_A6_2016 E-2 (E-2) is shown encircled by a dotted and dashed line (left inset) before being fractured along the dashed white line (center and right insets). The sediment adjacent to the egg proper is shown to be composed of thin layers in a CT scan at 3 μm resolution (**B**), which represent extra eggshell layers (ESL) adjacent to the fossilized egg (E-2) as seen under SEM (**C**). Extra eggshell layers are indicated by black arrowheads. The NUS_A6_2016 E-2’s eggshell is tentatively indicated by the white arrowheads and dotted line.

**Figure S68.**
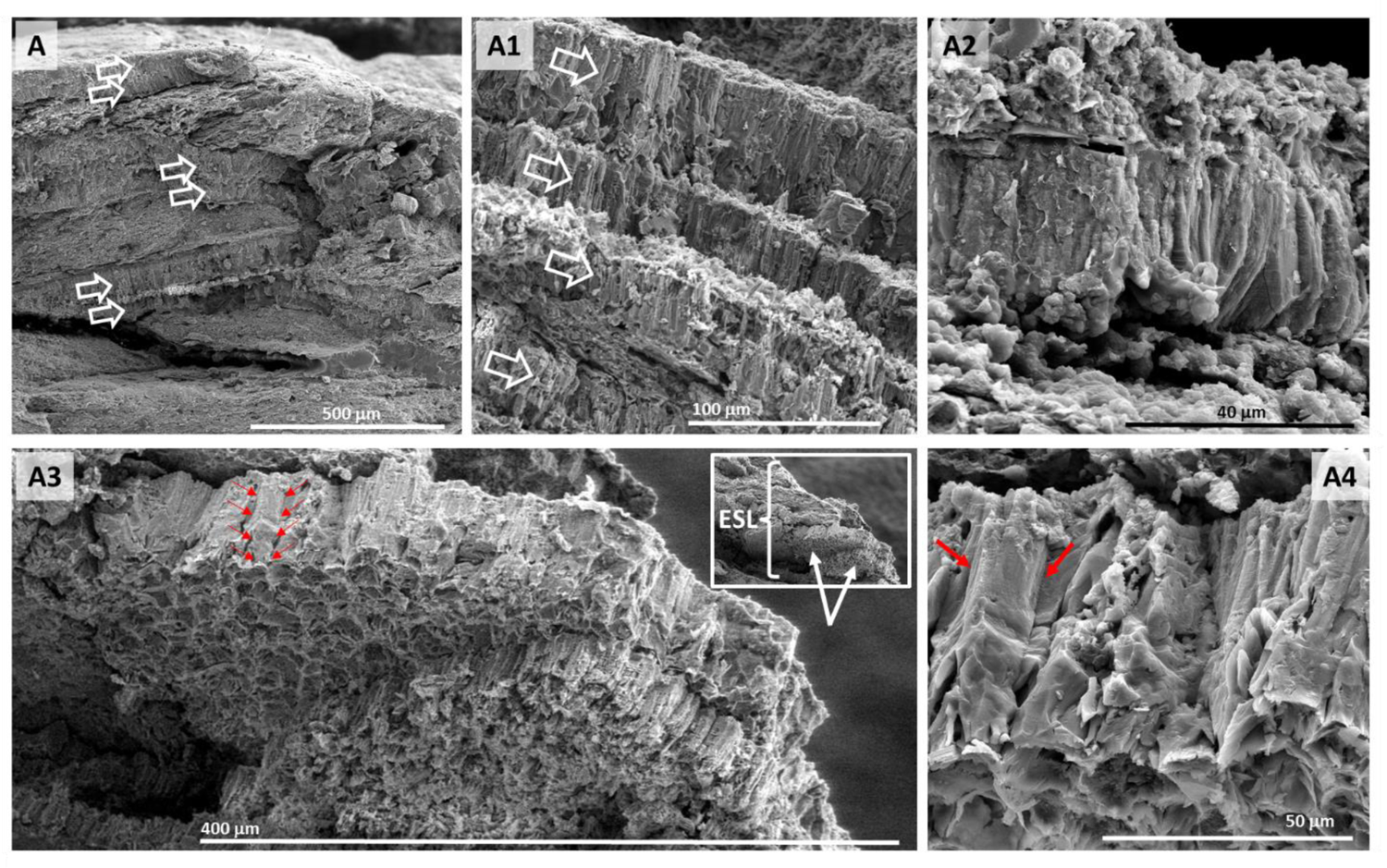
Layers of eggshell remnants associated with an egg extracted from the NUS_A6_2016 fossil. Extra eggshell layers adjacent to the NUS_A6_2016 E-2 egg are shown under SEM and indicated by open arrows in A and A1. A close-up view of one of these extra layers is shown in A2. A side view is shown in A3 and magnified in A4. Red arrows delimit an eggshell prism.

**Figure S69.**
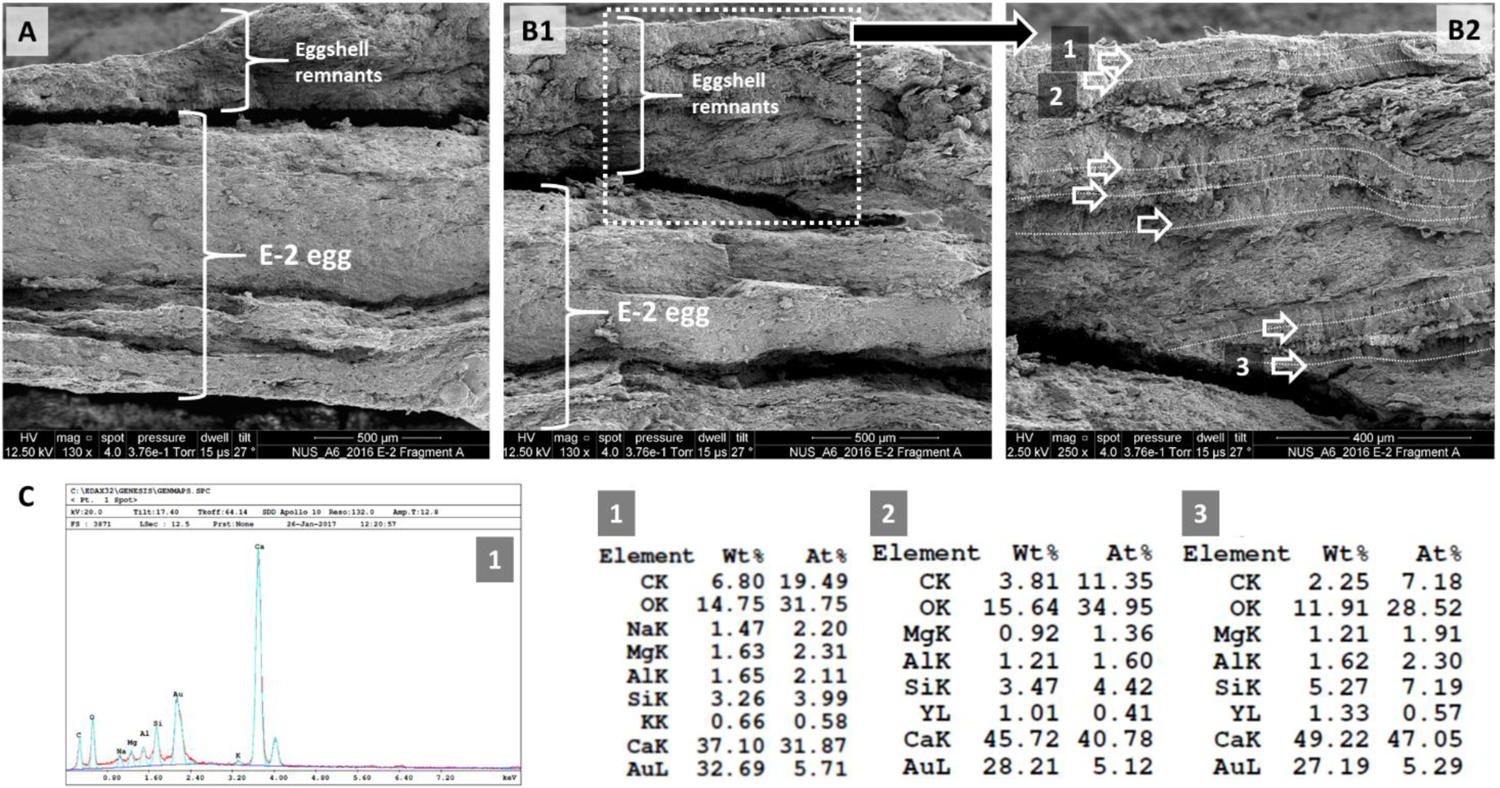
Layers of eggshell remnants. Extra eggshell layers adjacent to the NUS_A6_2016 E-2 egg under SEM and EDS. Layers indicated by numbers 1, 2 and 3 under SEM (top) are shown under EDS (below).

**Figure S70.**
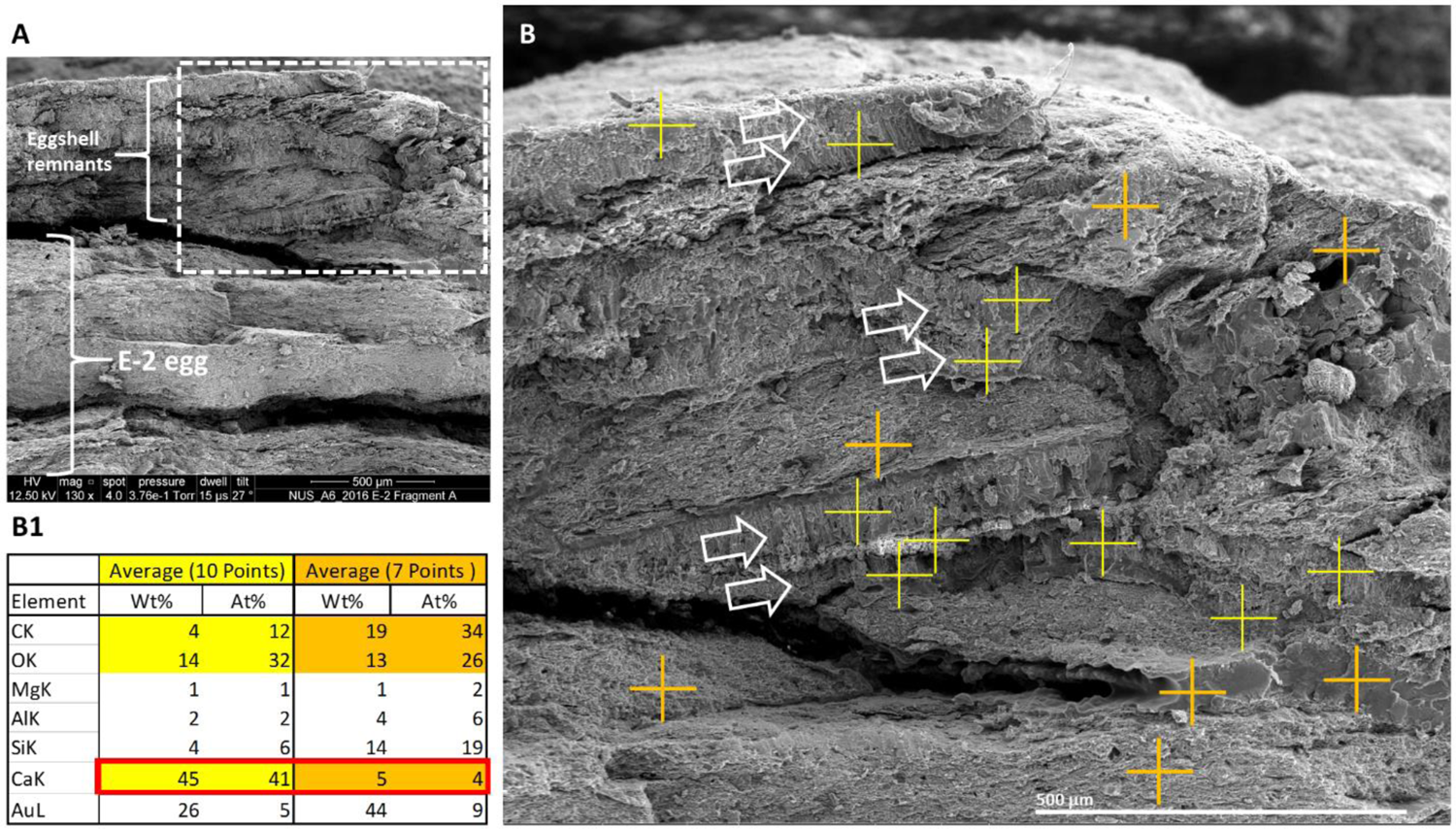
Multiple layers of eggshell remnants in the NUS_A6_2016 fossil confirmed by EDS analysis. The sediment adjacent to the NUS_A6_2016 E-2 egg (**A**) was studied further. Eggshells identified by white open arrows were studied by EDS (**B**). Yellow crosses reside within eggshells; orange crosses in the sediment. Average EDS readings of the points shown in B are shown in table format in **B1**. Calcium readings are outlined by a red rectangle.

**Figure S71.**
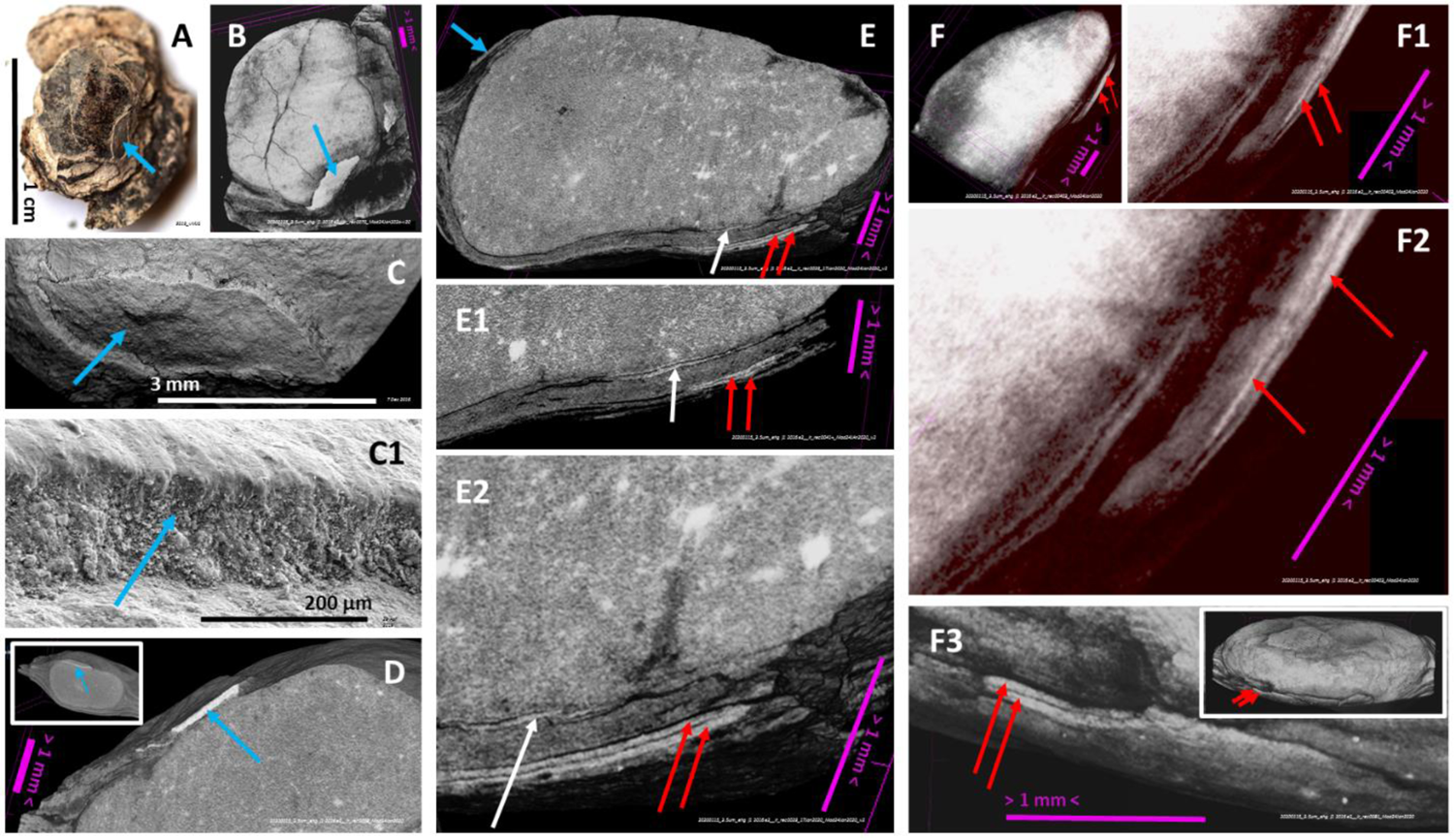
Eggshell remnants adjacent to EHG_J1_2016 E2 egg. Eggshell remnant (blue arrow) adjacent to EHG_J1_2016 E2 under macrophotography (**A**) and a 3D CT scan image at 3.5 μm resolution (**B**), SEM (**C**, amplified in **C1**) and virtual cuts of 3D views of CT scans at 3.5 μm resolution (**D**–**E**). This eggshell remnant was analyzed via EDS (data not shown). Other eggshell remnants (red arrows) were identified via virtual cuts (**E1** –**E2)** and 3D views of CT scans (**F**–**F3**), also at 3.5 μm resolution, in comparison with the eggshell closer to the egg proper (white arrow in E-E2).

**Figure S72.**
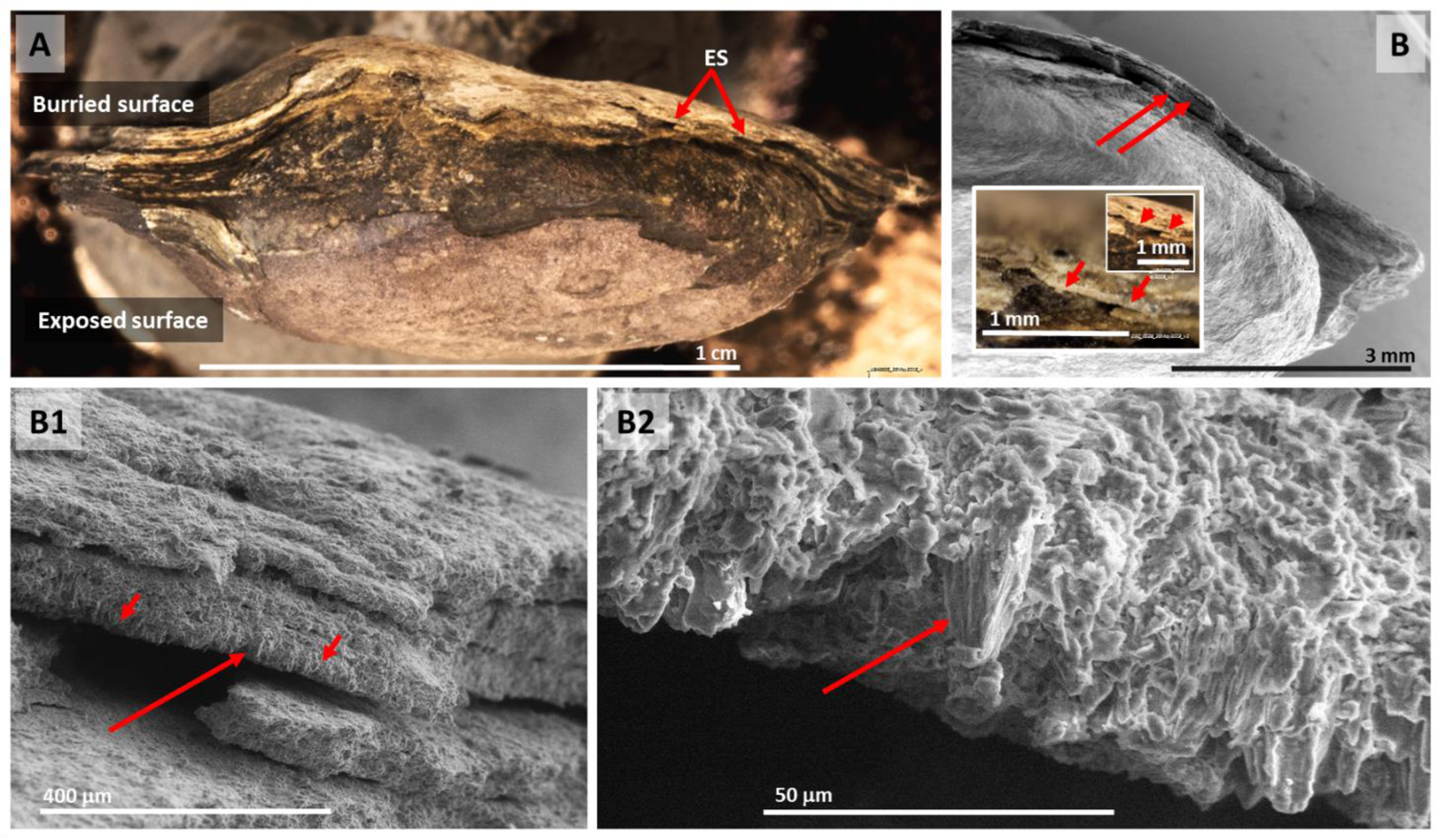
Eggshell remnants adjacent to the EHG_J1_2016 E2 egg. The same eggshell remnants (ES) adjacent to EHG_J1_2016 E2 indicated by red arrows in Figure S71 are shown here under macrophotography (**A** and **B**) and SEM (B). A calcium-rich crystal within one of these eggshell remnants (determined by EDS, data not show) is indicated by the longer arrow in **B1** and **B2**.

**Figure S73.**
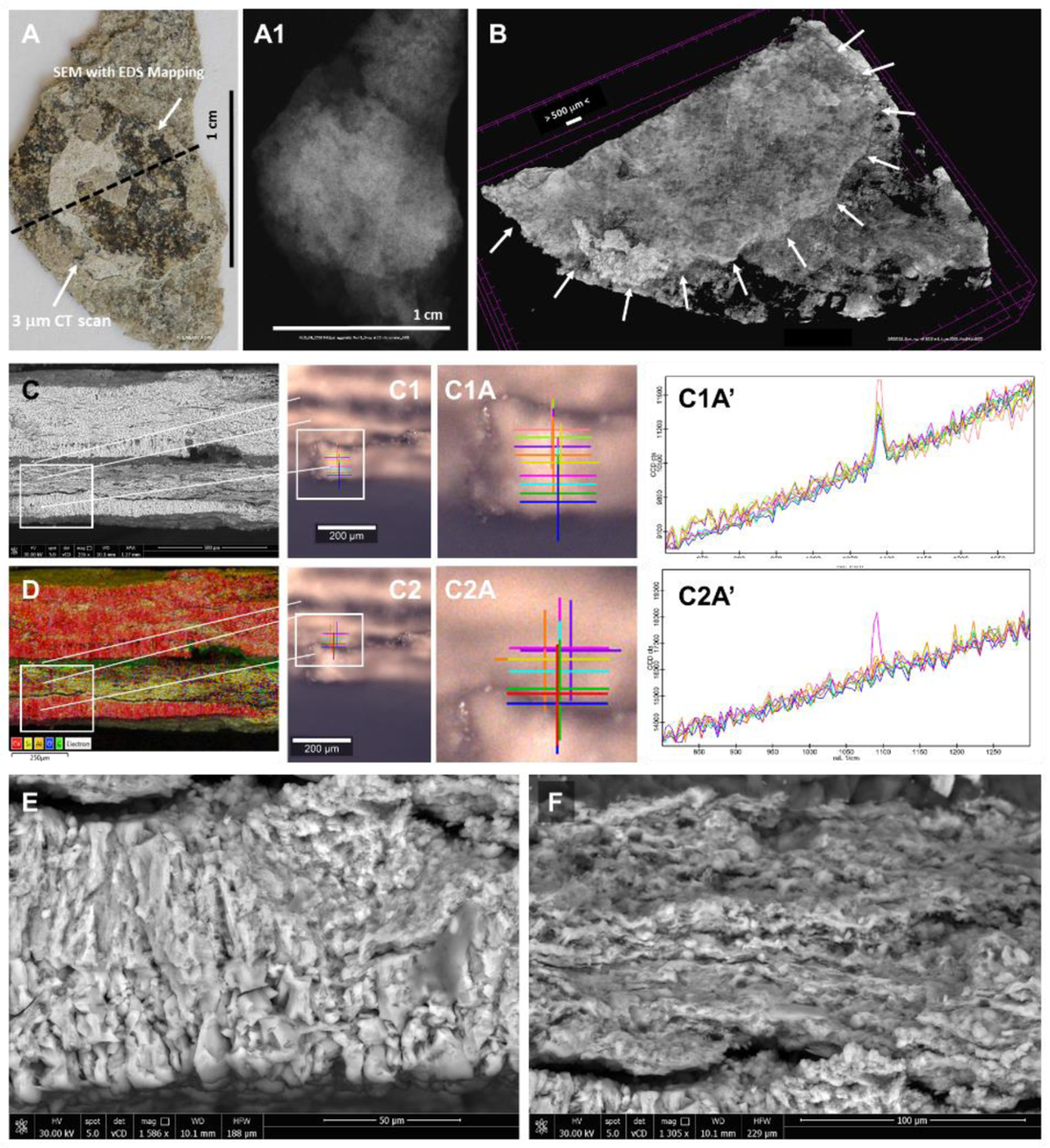
Isolated eggshell NUS_N9_2016 E-3 and associated extra eggshells. The NUS_N9_2016 E-3 eggshell was identified for its appearance (**A**), which was confirmed via X-ray at 3 μm resolution to conform to the size and 2D outline characteristic of the ornithuromorph eggs described in this manuscript (**A1**). It was fractured (see also Figure S74) and submitted to micro-CT scanning (**B**). Its radial surface (**C**) was submitted to RAMAN spectroscopy (**C1-C2)**, from the eggshell surface, which corresponds to the bottom of the white rectangle in C. This rectangle includes the sediment adjacent to the eggshell (yellow area of the rectangle in **D**; see Figure S75 for RAMAN spectroscopy results of the remaining radial surface of this sample). Characteristic calcite signature modes were observed in the eggshell (the calcite mode at ≈1091 cm^-1^ is consistently shown in all points measured in C1A’ but only in one isolated point from the adjacent sediment layer analyzed in C2A’). The entire radial surface of the sample was analyzed via EDS mapping (D), showing the separation of the NUS_N9_2016 E-3 eggshell, which corresponds to the calcium-rich (red) bottom layer, from the adjacent sediment (yellow layer above it, which is silica rich), a carbon-rich layer (green) and several layers of extra eggshells (top layers in red above them, already outside the white rectangle) intermixed with sediment. Magnified views of the NUS_N9_2016 E-3 eggshell and adjacent sediment under SEM are respectively shown in **E** and **F** (see also Figures S74-S77).

**Figure S74.**
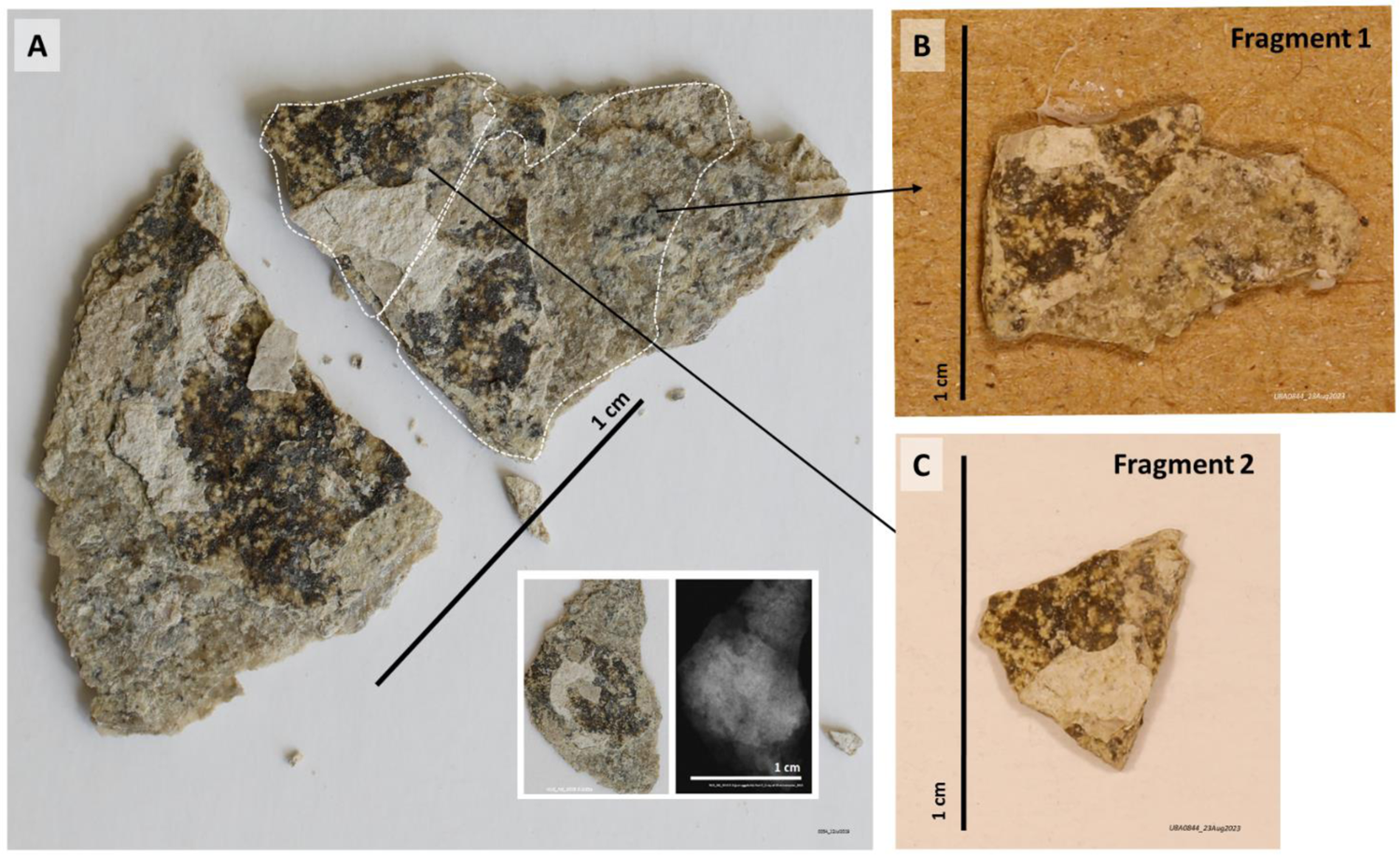
NUS_N9_2016 E-3 eggshell remnant. **A**: Before (inset, under macrophography and CT scan analysis) and after initial breakage, and subsequent intentional breakage of one of its halves into Fragment 1 (**B**) and Fragment 2 (**C**). The latter were used for further study of their freshly exposed radial surface under SEM and EDS.

**Figure S75.**
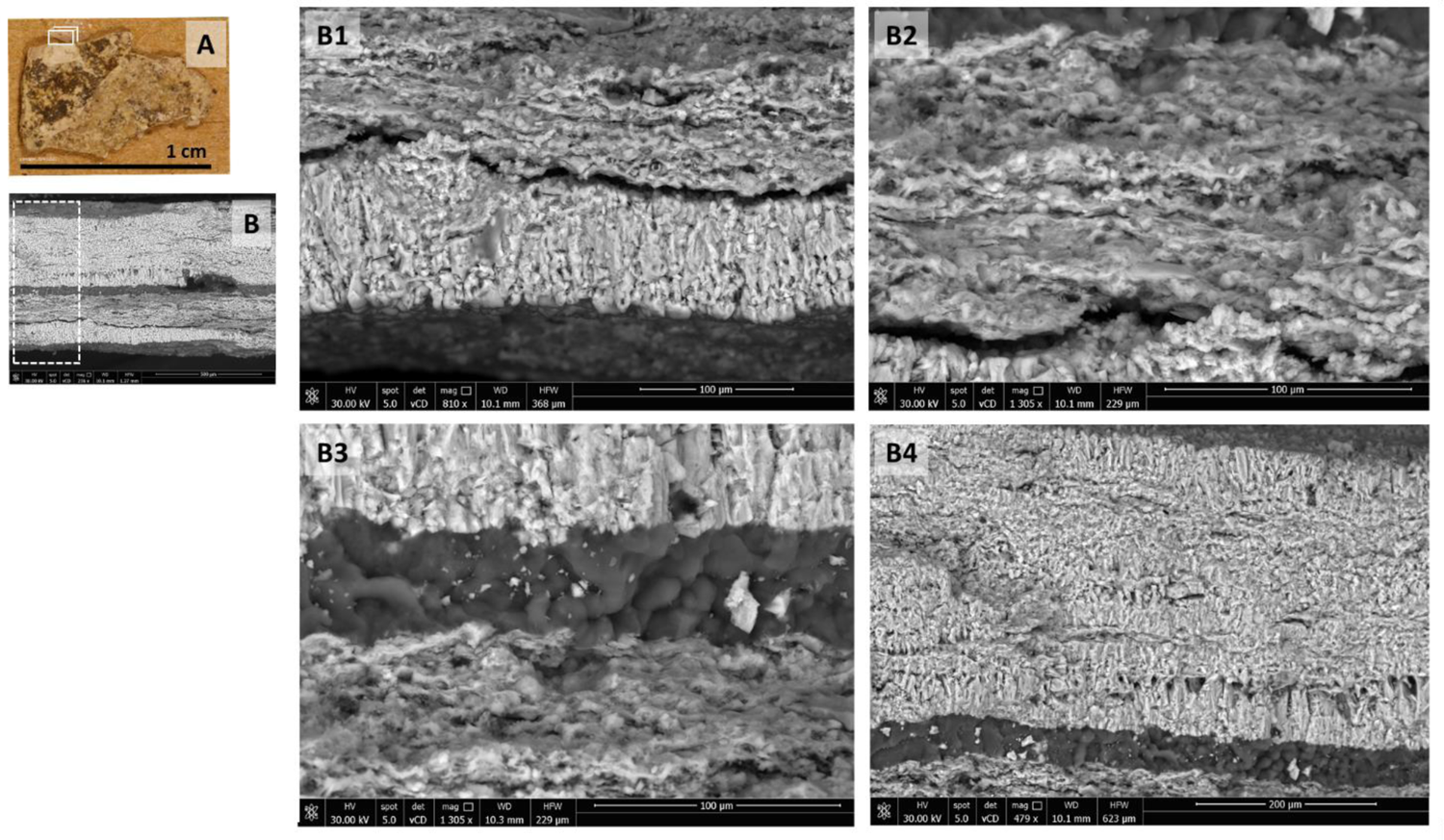
Extra eggshells adjacent to the NUS_N9_2016 E-3 eggshell. The radial surface of fragment 1 was studied in the area that corresponds to the white rectangular prism shown in **A**. The entire cross section of the fragment in that position is shown in **B** under SEM. Magnified views of the dashed rectangle represented in B are shown, from bottom up, in **B1**, **B2**, **B3** and **B4**. B1 corresponds to the NUS_N9_2016 E-3 eggshell, B2 to sediment adjacent to the NUS_N9_2016 E-3 eggshell, B3 to a carbon rich layer and B4 to extra eggshell layers (see also Figure S77).

**Figure S76.**
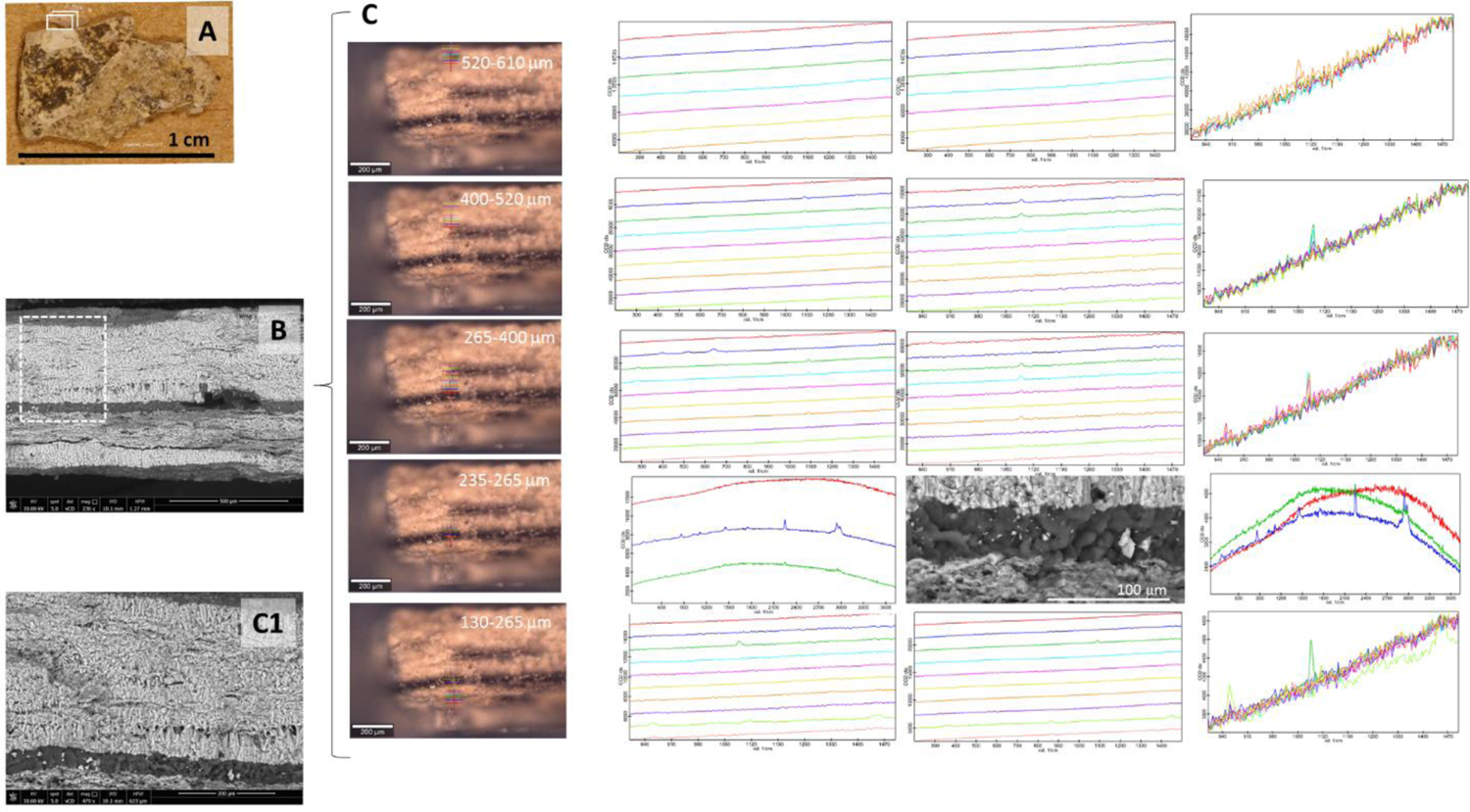
Extra eggshells adjacent to the NUS_N9_2016 E-3 eggshell. In addition to the layers of the radial view of fragment 1 that correspond to NUS_N9_2016 E-3 eggshell and the sediment that is immediately adjacent to it, already shown in Figure S73, the remainder of the radial section of the said fragment represented by the white prism in **A**, which corresponds to the dashed rectangle in **B**, were studied via RAMAN spectroscopy at intervals indicated as indicated in **C**. Characteristic calcite signature modes were consistently observed within intervals of approximately 100 m interspersed by interruptions of variable size in calcite signature modes (the calcite mode at ≈1091 cm^-1^ is shown; see also Figure S77). A carbon-rich layer was observed 235-265 m below the slab surface. An SEM view of the part of the section studied in C is shown in **C1**.

**Figure S77.**
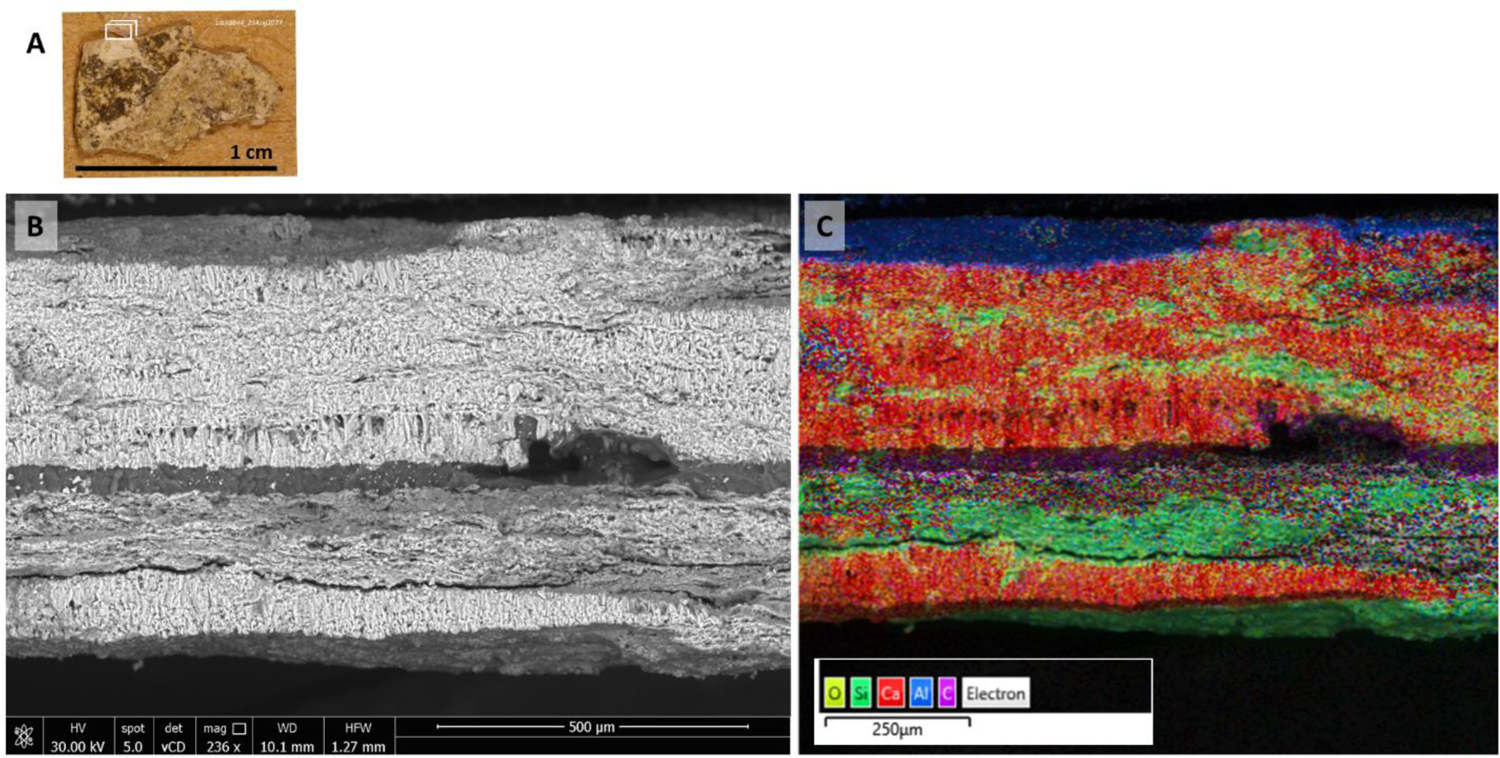
Extra eggshells lie beneath the NUS_N9_2016 E-3 eggshell. The radial surface of Fragment 1 of the NUS_N9_2016 E-3 eggshell (see Figure S74) at the location indicated by the white rectangle in **A**, which was analyzed via RAMAN spectroscopy in Figures S73 and S76, with details of its surface under SEM shown in Figure S75, is shown under SEM (**B**) and EDS mapping (**C**). In the latter, calcium is shown in red, silica in green and carbon in purple.

